# *E. coli* prepares for starvation by dramatically remodeling its proteome in the first hours after loss of nutrients

**DOI:** 10.1101/2024.02.29.582700

**Authors:** Théo Gervais, Bjoern Kscheschinski, Michael Mell, Nevil Goepfert, Erik van Nimwegen, Thomas Julou

## Abstract

It is widely believed that due to nutrient limitations in natural environments, bacteria spend most of their life in non-growing states. However, very little is known about how bacteria change their phenotype during starvation and what controls the concentration of different gene products inside the cells. Here we used microfluidics with quantitative fluorescence microscopy to quantitatively monitor growth and gene expression in many independent single-cell *E. coli* lineages as cells were switched from exponential growth to carbon starvation.

In contrast to the hypothesis that stationary phase at the population level may reflect a balance between continuing growth and death in different sub-populations, we found that all cells immediately enter growth arrest, that cells further in their cell cycle subsequently undergo reductive division, and no cell death occurs for more than two days. Second, we observed dramatic time-dependent changes in protein production that are highly homogeneous across single cells. Some promoters shut off protein production immediately, some show a slow exponential decay of production on a 10 h time scale, while others exhibit a transient burst of increased production before decaying exponentially at different rates. Notably, the reduction in protein production 30-60 h into starvation relative to production in exponential phase varies by more than two orders of magnitude across promoters and is highly correlated with production in the first 10 h of starvation.

Control experiments show that protein degradation itself also decays exponentially and using mathematical modeling we show how the fold-change in a gene’s protein concentration between exponential phase and late starvation depends on the size of the expression burst at the onset of starvation, the rate of subsequent production decay, and the rate of degradation decay. For many genes, the expression in late starvation is driven by production during the first 10 h. Finally, we establish that this expression program at the onset of starvation is critical for cell viability. In particular, by inhibiting gene expression during different periods of starvation, we show that tolerance to stress later in starvation is determined by gene expression occurring during the first 10 h.

Our study provides a foundation for quantitative studies of bacterial starvation by uncovering a gene expression program that fundamentally remodels the proteome during the first 10 h of starvation, is highly homogeneous across single cells, sets the proteome later in starvation, and is crucial for stress tolerance.

## Introduction

As it is relatively easy to observe the bulk exponential growth of bacteria in the laboratory and to measure the growth rates of different strains in different environments, most bacteriological studies have traditionally focused on exponentially growing cells. Differences in growth rate are also of obvious evolutionary importance in the wild, as faster-growing bacterial strains are able to consume available resources before other competing strains.

However, it is generally believed that in their natural environments, bacteria are often starving for nutrients and may thus likely spend most of their time not growing, and eventually dying of starvation or stresses present in the environment. Indeed, if the total bacterial population on earth remains roughly constant, the total amount of bacterial growth and death must be balanced.

Hence, one would *a priori* expect the selection pressure for survival, and for surviving starvation in particular, to be as important as the selection for fast growth. Although not much is known about the mortality dynamics of starving bacteria, it has been observed that different bacterial species die at different rates [1], suggesting that differences in bacterial physiology can have a strong impact on mortality. This argues that the study of the states of bacteria in non-growing conditions, and how these affect survival rates is as important as the study of bacteria in exponentially growth conditions.

The study of the behavior of growth-arrested bacteria is also highly relevant from a medical point of view. First, various pathogenic bacterial species induce their virulence genes at the transition to stationary phase [2–5]. Second, bacteria in stationary phase have higher levels of tolerance to antibiotics [6–8], and the fraction of persister cells is much higher in stationary phase cultures than in exponentially growing ones [9, 10].

A straightforward way to induce non-growing states in the laboratory, at least at the population level, is to simply let bacterial populations grow until they exhaust their nutrients, after which they enter ‘stationary phase’ where the population size initially remains constant and then decreases due to cell death. This stationary phase is therefore a practical setting for studying bacterial starvation.

Despite the importance of understanding bacterial states in stationary phase both from an evolutionary and medical perspective, relatively little is currently known. Previous studies of stationary phase have revealed that, in general, cells become much less active: their energy level sink [11–13], their DNA becomes coated by the protein Dps which is not expressed in growing cells [14], and the composition of the cytoplasmic membrane and the cell wall changes to make them sturdier [15, 16]. However, it is also known that the precise physiological state of the cells during starvation depends on the physico-chemical conditions (*e*.*g*. limiting nutrient, osmolarity, pH, *etc*.) [15, 17], and also on how quickly bacteria transition from growth conditions to growth arrest or how long the bacteria have been in starvation.

Here we focus on quantifying the time evolution of gene expression as cells switch from exponential growth into carbon starvation, and subsequently during the first few days of starvation. Since bulk expression measurements mask variability at the single-cell level and cell-to-cell variability is believed to be higher in stationary phase [17, 18], we here use high-throughput microfluidics in combination with time-lapse microscopy to quantify the time evolution of gene expression of a substantial collection of transcriptional reporters at the single-cell level. In a previous study, it was reported that the expression of a synthetic inducible promoter exhibited constant gene expression, at ≈ 10% of the level in exponentially growing cells, over 50 h in stationary phase [19]. This indicates that, at least for some genes, there is sustained gene expression activity deep into stationary phase, but it is currently unclear to what extent this observation generalizes to different genes, and how the time dynamics of expression varies across single cells.

Using a microfluidic device that allows a precisely controlled switch in environment [20], we tracked many independent lineages of single cells as they were switched from growing exponentially on glucose to a media without a carbon source, and found that all cells immediately enter growth arrest upon removal of the carbon source. After going into growth arrest, cells that are further along in their cell cycle undergo reductive division, causing the average cell size of the population to decrease. In addition, we found that all cells remain growth arrested without any cell death for more than two days in our microfluidic device.

We further found that gene expression undergoes dramatic changes upon entry into starvation with a wide variety of behaviors across promoters. Some genes cease protein production immediately, some slowly fade in protein production on a time scale of approximately 10 h, and others show a transient increase in expression, after which production also slowly fades away over time, decaying approximately exponentially. In addition, we found that protein degradation also decays exponentially with time and, using mathematical modeling, we show how the protein concentrations in late starvation are determined by the combination of growth arrest (which halts dilution), the transient increase in expression, the rate of exponential decay in protein production, and the exponential decay in protein degradation. Notably, except for genes such as ribosomal genes that immediately halt protein production, the sustained expression over the first hours in starvation in combination with growth arrest causes many genes to increase protein concentration compared to protein concentration during exponential growth, even though their production was never upregulated. In general, for most genes the protein concentration deep in starvation is determined by the expression in the first hours after growth arrest. Thus, the expression program in the first hours of starvation is the main determinant of the proteome composition in late starvation.

Using a setup where a batch culture is flown through the microfluidic chip as it progresses from exponential phase to stationary phase, we also show that the gene expression response is similar between abrupt and gradual growth arrest. Finally, we demonstrate that the transient expression upon entry into starvation has major phenotypic consequences. In particular, we show that the expression changes early in starvation are crucial for conferring resistance to stress later in starvation.

## Results

In order to study gene expression under starvation with single-cell resolution, we used an integrated setup which combines microfluidics, time-lapse microscopy and automated image analysis [20]. After inoculation in the Dual Input Mother Machine (DIMM) microfluidic chip, *E. coli* cells were first grown in M9 minimal media supplemented with 0.2% glucose for 10 h and then switched to M9 minimal media without carbon source (referred to as “M9 zero”) for up to 60 h. To assess the survival of cells at the end of this starvation period, cells were then allowed to regrow in fresh media for an additional 10 h in some experiments. We used a multiplexed version of the DIMM chip that allows us to measure up to 8 strains in parallel, each carrying a different transcriptional fluorescent reporter (Fig. 1A).

**Figure 1.**
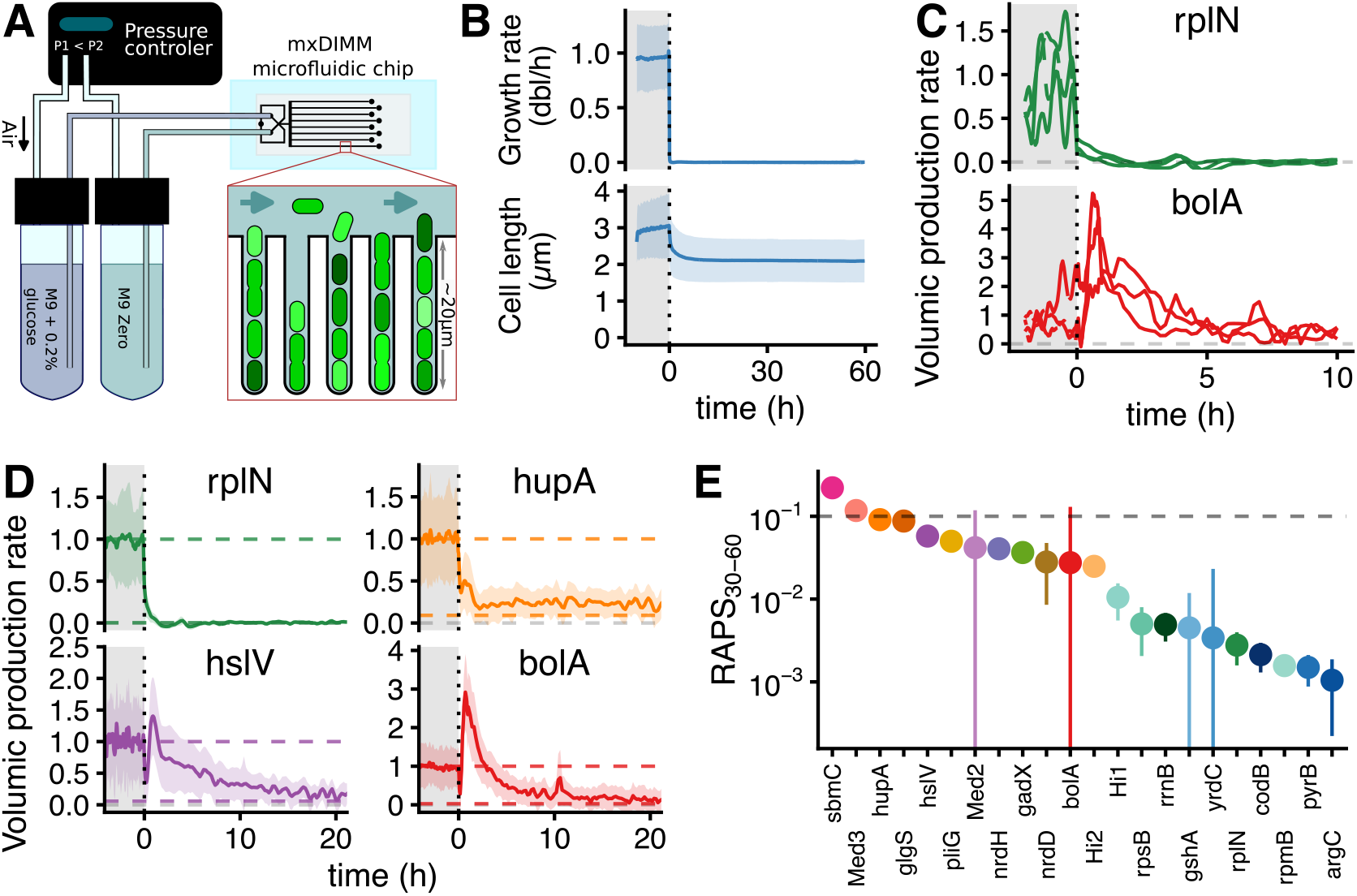
Characterisation of single-cell gene expression during starvation using microfluidics and time-lapse microscopy. (A) Size and fluorescence of individual *E. coli* bacteria were monitored as cells were switched from exponential growth (M9 minimal media + 0.2% glucose) to starvation conditions (“M9 zero” without carbon source) in a modified DIMM microfluidic chip [20], allowing measurement of up to 8 strains in parallel. (B) After switching to “M9 zero”, elongation stops immediately (top panel, Fig. S2), and average cell size decreases due to reductive divisions (bottom panel). Lines and ribbons show mean and mean ± s.d. across all single cells, respectively; time 0 was defined as the time of the media switch. (C) Examples of single-cell traces of the volumic production rate *q* for the rplN promoter (top panel) and the bolA promoter (bottom panel). The volumic production rate is defined as the amount of fluorescent protein produced from the transcriptional reporter per unit time and per unit cell volume. Note the consistency in the time-dependent response in production upon entry into starvation across the single cells. (D) Time-dependent volumic production rates for four example promoters (out of 22) illustrate the diversity of the responses upon entry into starvation (only the first 20 of 60 h are shown); lines and ribbons show mean and mean ± s.d. across all single cells, respectively, normalized by the average value during exponential growth. Coloured dashed lines are guides for the eye showing 1 (i.e. the production during exponential growth) and the average relative production rate in the last 30 h (Relative Average Production in Starvation between 30 and 60h, RAPS_30-60_). (E) Production rates late in starvation relative to production rates in exponential growth (RAPS_30-60_) for 18 native and 4 synthetic promoters vary over more than 2 orders of magnitude. Dots and error bars correspond to the mean and mean ± s.e.m, respectively. The horizontal dashed line indicates the value reported previously for a synthetic promoter in [19].

### Inference of instantaneous growth and protein production rates

The cell size and the total fluorescence were measured at each time point and for every individual cell in the experiment using our image analysis software deepMoMA (see SI 3). We used this data to estimate the instantaneous growth rate *⁁*(*t*) and volumic production rate *q*(*t*) as measures of growth and transcriptional activity of every cell at each time point *t*, respectively. Formally, we define the instantaneous growth-rate *⁁*(*t*) of a cell as the time derivative of the logarithm of cell volume

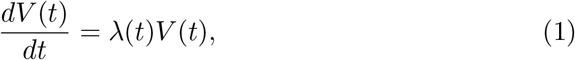

with *V*(*t*) the cell’s volume at time *t*. We further define the instantaneous volumic production *q*(*t*) as the rate at which fluorescent proteins are produced per unit time and per unit volume of the cell. That is, we assume that the total fluorescence *F*(*t*) of the cell evolves as

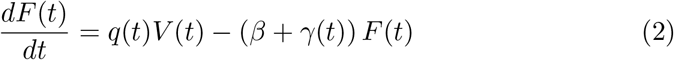

where *β* is the rate of photo-bleaching and *γ*(*t*) is the rate of degradation of fluorescent protein, which as we will see below, depends explicitly on time *t* during starvation.

Note that, because ribosome concentration is believed to be approximately constant independent of cell volume, and the fluorescent reporters of different promoters have essentially the same mRNA, the production rate *q*(*t*) is proportional to the product of the mRNA concentration of the reporter and the total protein production rate per mRNA.

There are several challenges in inferring the growth rates *λ*(*t*) and production rates *q*(*t*) from the raw measurements. First, the raw measurements of cell size and fluorescence are affected by significant measurement noise. For example, simply taking discrete derivatives between consecutive measurement time points of the logarithm of cell volume would result in estimates of growth rate that are far too noisy to be useful, and the same applies to inferring *q*(*t*) from changes in measured total fluorescence.

To address this challenge, we recently developed an advanced Bayesian inference procedure, called RealTrace, that removes measurement noise by assuming the growth and production rate are both smoothly fluctuating with time, without making specific assumptions about the functional form of these fluctuations. In particular, RealTrace uses maximum entropy process priors in which the smoothly fluctuating growth and production rates are characterized only by their mean, variance, and correlation time. For each reporter, RealTrace first fits the parameters of the priors separately for the exponential growth and starvation phases, and then uses these priors to infer posterior distributions for the instantaneous growth and production rates of every cell at every time point (Fig. S1, and SI section 5).

A second challenge is that RealTrace assumes that bleaching and protein-degradation are constant in time. However, we performed calibration experiments to determine protein degradation rates (Fig. S8, SI section 6) and these uncovered that the protein degradation rate is decreasing approximately exponentially with time during the starvation phase, i.e. *γ*(*t*) = *γ* _0_*e*^*− ωt*^, if we define *t* = 0 as the start of the starvation phase. To infer the volumic production rates *q*(*t*) we thus proceeded as follows. After estimating the time-dependent degradation rates *γ*(*t*) of the fluorescent proteins, we performed additional calibration experiments to estimate the photobleaching rates *β* of the fluorescent proteins (Table S6, SI section 7). We then ran RealTrace assuming fluorescence only decays through photobleaching, i.e. assuming the total fluorescence of the cell evolves according to 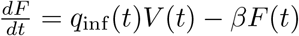. Because this inference neglects protein degradation, the inferred production rate *q*_inf_ (*t*) will systematically underestimate the true production rate *q*(*t*). But because we have estimated the time-dependent protein degradation rate *γ*(*t*), we can correct for this underestimation to infer the true production rate *q*(*t*) (SI section 8).

### Immediate growth-arrest and reductive divisions upon entry into starvation

The most readily observable signatures of starvation are the arrest of growth at the population level, and the decrease of average cell size. Although it has been reported recently that *E. coli* cells are all growth-arrested during stationary phase [21], our experiments are the first to measure the changes in growth at the single-cell level with high time resolution as cells are abruptly switched from exponential growth to carbon-free media.

We observed that, upon the switch from M9 + 0.2% glucose to M9 zero, all bacteria stop growing immediately (within the 3 minute microscopy acquisition time; Figs. 1B and S2A,B). In addition, after growth has completely ceased, a large fraction of cells (38%) undergoes division within the first 10 h, with most of these reductive divisions occurring early and their frequency decreasing with time (Fig. S2C), leading to a decrease of the average cell size with time (Fig. 1B). That average cell size decreases upon entry into stationary phase has been observed previously [21, 22], and this has been attributed to a different mechanism of cell size control during the last cell cycle(s). However, our data shows that reductive divisions occur after cell growth has already ceased completely. Moreover, we find that whether a cell will undergo a reductive division is well predicted by its size at the time of entry into starvation, i.e. as measured either by the cell’s absolute length or added cell length since birth (Fig. S2D). It is tempting to speculate that, perhaps, the cells that undergo reductive division are those in which DNA replication has already been completed.

In summary, in contrast to the hypothesis that stationary phase at the population level might reflect a balance between growth and death at the singlecell level, we find that all cells immediately arrest growth upon loss of carbon source, and that the reduction in average cell size results from the fact that longer cells undergo a division during the first hours of starvation. Moreover, the fact that we observe all cells stop growing immediately indicates that our setup used with M9 zero constitutes a powerful platform to study starvation at high time resolution, under controlled conditions.

### Dramatic and highly consistent remodeling of gene expression upon entry into starvation

We measured gene expression dynamics during starvation using transcriptional fluorescent reporters for a set of native and synthetic promoters selected as follows. For native promoters, we took advantage of previous flow cytometry measurements [23] to pick 18 promoters spanning a broad range of expression and of fold-changes between growth and stationary phase (Fig. S3). In order to quantify expression of constitutive genes, we also used 4 synthetic promoters that were obtained by screening random sequence fragments for their expression level during exponential growth (either “medium” or “high”) [24], and for which we verified the absence of regulatory inputs other than sigma factor binding sites (RpoD/RpoS).

In previous works it has been observed that, when conditions change, different subpopulations of single cells exhibit highly variable responses in gene expression [25–27] and we thus anticipated that the expression responses in single cells upon switch to starvation would also exhibit large variability. However, we not only find that upon entry in starvation, the protein production rates of different promoters undergo dramatic changes that are highly diverse across promoters, but that these changes are also highly consistent across single cells. As an example, Fig. 1C shows single-cell production traces for the ribosomal protein promoter rplN and the rpoS target promoter bolA (Fig. S4 shows example single-cell production traces for all 22 promoters). The observed coherence of the responses across single cells suggests that *E. coli* has evolved a tightly controlled regulatory program in response to starvation.

Given how reproducible the time-dependent responses are across single cells, we chose to first analyze the responses across promoters based on their time-dependent means and standard deviations across the population, and will return to analysis of the variability across single cells later on. Figure 1D shows the volumic protein production rate across time for 4 native promoters representative of the diversity of the dynamics that we observe across the 22 promoters (Fig. S5A). Note that values are scaled to the average volumic production rate during exponential growth and that our set of promoters covers a range of absolute volumic production rates that spans more than 2 orders of magnitude during growth, and more than 3 orders of magnitude during starvation (Fig. S5B).

Overall, protein production rates change dramatically at the onset of starvation. As expected, we observed that mean volumic protein production rate after prolonged starvation is always lower than it was during exponential growth. As can be seen from Fig. 1D, production patterns are diverse and rather complex upon entry in starvation: Certain promoters, such as the ribosomal promoter rplN, undergo a sharp down-regulation of production that almost stops their production within an hour of the switch. Other promoters, such as hupA, see a sharp initial drop in production, followed by a slower decay, and even a very long sustained production at a lower level. Finally, promoters such as hslV and bolA show a transient burst of variable amplitude in production, increasing production over its value during exponential growth, followed by a slower decay in production that proceeds at different rates for different promoters.

### Slow protein production decay at variable rates across promoters

In a previous study, it was found that one promoter exhibits approximately *Constant Activi*ty *in Stationary Phase* (CASP) at the population level over a 50 h period [19]. In line with this observation, we also find that many promoters still have detectable protein production 60 h after the switch to starvation, i.e. at the end of our experiments (Fig. S6A). However, rather than a constant production, we observed that the activity of all promoters keeps decreasing slowly over this timescale and can often be well fit by a mixture of two exponential decay functions (SI section 9 and Fig. S6B).

To compare the activity of different promoters during starvation, we decided to measure the average volumic production rate over defined time windows, relative to the average rate during growth, e.g. RAPS_30-60_ refers to the Relative Average Production in Starvation during 30-60 h of starvation. If the decay in production were driven primarily by the overall reduction of energy or of building blocks availability, then we would expect all promoters to have similar RAPS_30-60_ (estimated at approx. 10% in [19]). In contrast, our data shows that RAPS_30-60_ varies by more than 2 orders of magnitude (from 20% down to 0.1%) in our set of 22 promoters (Fig. 1E and Fig. S5B). The fact that RAPS_30-60_ is so variable across promoters underscores the profound changes that the gene regulatory network undergoes during starvation.

### Time-dependent protein concentration changes

While protein production rates arguably define the current gene expression state of the cell, protein concentrations arguably define the physiological state of the cell, e.g. they dictate the rates at which different biochemical reactions occur within the cell. We thus also aimed to track how protein concentrations change across time for our promoters.

To obtain time traces of protein concentration for our fluorescent promoters (Fig. 2A) we numerically integrated our estimates of the true time-dependent protein production rates *q*(*t*) accounting also for the time-dependent protein degradation rate *γ*(*t*) (SI section 8 for more details on the calculation). As mentioned above, our calibration experiments on the degradation of fluorescen proteins show that degradation *rates* decrease exponentially, i.e.. *γ*(*t*) = *γ* _0_*e*^*− ωt*^, starting from initial values of degradation rate (γ_0_ ∈ [0.02 − 0.03] h^-1^, see Table S5). These rates are compatible with the overall protein degradation rate at vanishingly slow growth [28] and the marked slow-down of degradation during starvation is also consistent with the fact that the vast majority of protein degradation in *E. coli* is performed by ATP-dependent proteases [29,30] and that ATP levels decrease dramatically during starvation [12,13,31]. Furthermore, changes in the levels of metabolites related to protein degradation upon entry into carbon starvation also indicate that degradation slows down during starvation [11].

**Figure 2.**
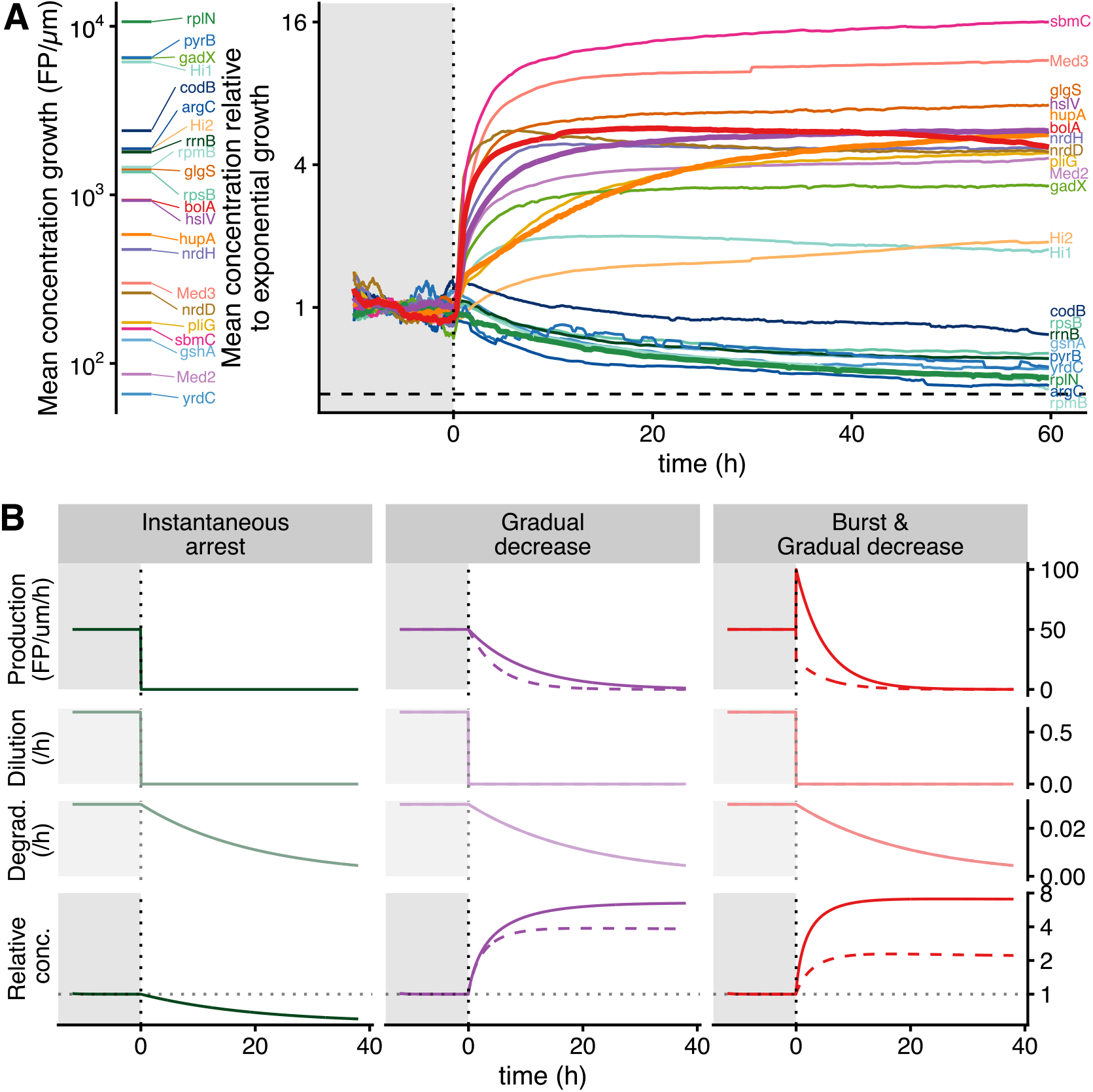
Dynamics of protein concentration during the switch from growth to starvation. (A) The protein concentrations corresponding to promoters used in this study span a broad range (ca. 100 fold) during exponential growth (left), and change in a promoter-dependent manner upon entry into starvation (right). Protein concentrations are measured in fluorescent protein per *µ*m (FP/*µ*m) and corrected for photobleaching (SI section 8). Time traces show averages across all cells, scaled to the average concentration during exponential growth and shown on an logarithmic scale; promoters shown in Fig. 1D are indicated with thick lines. The horizontal dashed line shows the minimal concentration that would be reached, theoretically, in the case of instantaneous arrest of both production and growth upon entry into starvation (SI section 12). (B) Examples of protein concentration dynamics (bottom row) resulting from the combined time dynamics of protein production (top row), dilution (second row), and protein degradation (third row) for three archetypical scenarios of production dynamics (columns). In all scenarios dilution drops to zero as growth stops immediately after the switch (Fig. 1.B) and degradation rates decay exponentially during starvation as measured experimentally (Fig. S8). On the left protein production halts immediately upon the switch, in the middle it decays exponentially (at two different rates), while on the right there is an initial sudden change in production, followed by exponential decay. All parameters of the model are set after experimental observations (SI section 12). Note that the predicted concentration fold-changes are comparable to those observed experimentally in panel A.

Since the results in Fig. 2A use our measurements of degradation rates for the fluorescent proteins, extending these results to the native proteins of these reporters requires the assumption that the degradatation rates of these proteins are similar to that of the fluorescent proteins. To validate this assumption, we used quantitative proteomics to measure the temporal dynamics of native protein levels in batch cultures (SI section 11). However, we found that the accuracy of the proteomics measurements was insufficient for this purpose (Fig. S7). Consistent with this observation, publicly available proteomics datasets comparing growth and starvation show dramatic variability between replicates and datasets, making it impossible to obtain reliable quantitative estimates of the change in concentration of individual proteins (Fig. S7). Based on available knowledge, we thus assume that the overall decrease of degradation due to reduced energy levels is the dominating effect for most genes, and that concentrations of fluorescent proteins used as transcriptional reporters are representative of the concentrations of the corresponding native proteins.

It is noteworthy that, among our set of promoters, there are almost none whose protein concentration remains stable across the switch, i.e. concentration either decreases due to degradation after a sharp production arrest, or increases more than 2-fold (Fig. 2A). Notably, synthetic promoters reporting constitutive gene expression (high expressors) are the closest to staying constant in concentration. For downregulated promoters, despite dilution stopping, the concentration can drop as low as ≈ 50% of their exponential levels due to protein degradation (Fig. 2A). In contrast, for promoters whose concentration increases, the relative change can be much larger, up to 12-fold across our set of 22 promoters. Strikingly, for those promoters, most of the changes seem to happen within the first 10 h of starvation, suggesting that the differences in production observed after prolonged starvation, *i*.*e*. RAPS_30-60_, are not playing an important role in the final concentrations reached at the end of the experiment.

Perhaps counter intuitively, even for promoters that do not exhibit a temporal burst in production but only undergo decreasing production, protein concentrations can increase substantially (i.e. more than 2-fold). Among the 4 examples of Fig. 1D, this is the case for both hslV and hupA (Fig. 2A right), although hupA takes far more time to reach the same fold-change. Mechanistically, this increase in protein concentration comes from the delay between the growth arrest (which is almost instantaneous) and the arrest of protein production (which is gradual and promoter-dependent). As a consequence, even for promoters for which protein production is reduced to very low levels during starvation, the protein concentration can nonetheless increase substantially during starvation.

### A simple mathematical model captures the diversity of observed concentration dynamics

Although the protein production time courses are highly diverse between promoters and differ in their details, it seems that most of these dynamics can be captured by two main variables: the amplitude of the initial burst of production and the delay between growth arrest and production arrest. We thus explored to what extent the diversity of protein concentration dynamics that we observe in our experiments can be captured by a simplified mathematical model (SI section 12). In this model all cells immediately arrest growth upon the switch to starvation (thereby ceasing dilution) and protein degradation decreases exponentially with time, i.e. with degradation rate 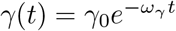 as observed in our calibration experiments (Fig. S8).

For the protein production rate *q*(*t*) we assume that, upon entry into starvation, the production rate undergoes an instantaneous fold-change of size *b*, i.e. with *b* > 1 corresponding to a burst in production and *b* < 1 to an instantaneous reduction in production. After this, we assume that the protein production *q*(*t*) decreases exponentially at a rate *ω*_*q*_ that may differ between promoters.

Several simple scenarios of this model are illustrated in Fig. 2B: in a first scenario, production stops instantaneously upon entry into starvation, i.e. *b* = 0. A second case consists of a simple exponential decay of the production rate after entering starvation, corresponding to a delayed production arrest (i.e. *b* = 1, and the figure shows two example values of *ω*_*q*_). Finally, upon entry into starvation there may be a transient burst of production (*b* > 1) or an immediate decrease in production (*b* < 1). As shown in Fig. 2B, the concentration dynamics resulting from the cessation of dilution, the time-dependent protein degradation, and these time-dependent production rates, recovers the diversity of protein concentration dynamics that we observe experimentally.

As shown in SI section 12, the fold change in protein concentration between exponential growth and late stationary phase can be calculated analytically for this model and depends only on 3 effective ratios of rates. First, the total amount of decay is set by the ratio *r*_γ_ = γ_0_/*ω*_*1*_. In particular, for a promoter that ceases production immediately upon entry into starvation, a fraction 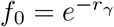 remains deep in stationary phase, corresponding to about 44% of the initial concentration as illustrated in the left column of Fig. 2B.

Besides *r*_*γ*_ the fold-change in concentration increases with the ratio *r*_*q*_ = *bλ /γ*_0_, which corresponds to the ratio of the fold-change in production *b* and the fold-change in degradation/dilution γ_0_*/λ*. Finally, the fold-change also depends on the ratio *r*_*t*_ = ω _*q*_*/ω*_*γ*_ of the time scales at which production and degradation decay.

Introducing a delay in the arrest of production is enough to produce a large increase of concentration (e.g. there is a 4-fold increase when the production decays with a characteristic time of 5h, Fig. 2B, middle column). The early burst of production can further tune the concentration fold-change observed between growth and starvation (e.g., 7-fold for bolA, right column of Fig. 2B).

Because bolA is a known reporter of RpoS activity, we hypothesize that the increase in concentration we observe for this promoter, and by extension for others that display the same dynamics, are the results of rpoS activity. To test this hypothesis, we studied the concentration levels and fold change for some of our promoters in a. ΔrpoS background. We find that promoters going down in concentration in the wild type display a slightly elevated concentration, while promoters going up in concentration during starvation in the wild type display a great decrease of concentration in the. ΔrpoS background (Fig. S9A). Surprisingly, knocking out rpoS did not always results in a decrease of the concentration fold change upon entry into starvation (Fig. S9B). Together, those results demonstrate the importance of RpoS on gene regulation during starvation, but suggest that other important regulation mechanisms are at play in this condition.

### Gene expression dynamics under progressive entry into starvation is similar to that under a hard switch

Our results so far concern the gene expression response upon a sudden ‘hard switch’ from exponential growth, to starvation conditions without a carbon source. Such an almost instantaneous depletion of nutrients might be rare in the natural environments of *E. coli*, and it is conceivable that the gene expression response might be quite different when the depletion of the nutrients is slower, perhaps allowing cells to sense the loss of nutrients.

To address this question, we did another series of microfluidic experiments where a culture growing in M9 + 0.02% glucose is flown through a mother machine microfluidic chip, so as to expose bacteria in the chip to the same physico-chemical conditions as in the culture flask, where nutrients are depleted due to the uptake by bacteria (Fig. 3A), following the same principle as in [19,21]. We observe that under this progressive entry into starvation, the instantaneous growth rates decrease over a longer period than for the hard switch. That is, while the growth rate dropped from its maximum value to zero in ca. 5 min in experiments with instantaneous switching, this transition takes ca. 45 min in these culture overflow experiments, which almost certainly indicates that nutrients are exhausted more gradually (Fig. S10A,B). It is interesting to contrast this timescale with the sharp angle of the growth curve observed at the population level in this media (Fig. 3B). For comparison of the gene expression dynamics between the two transitions, we define the *t* =0 start of starvation for the overflow experiments as the time when the instantaneous growth rate (averaged over all individual cells) has decreased by 95% (which corresponds to 0.055 dbl/h; Fig. 3B and Fig. S10A,B).

**Figure 3.**
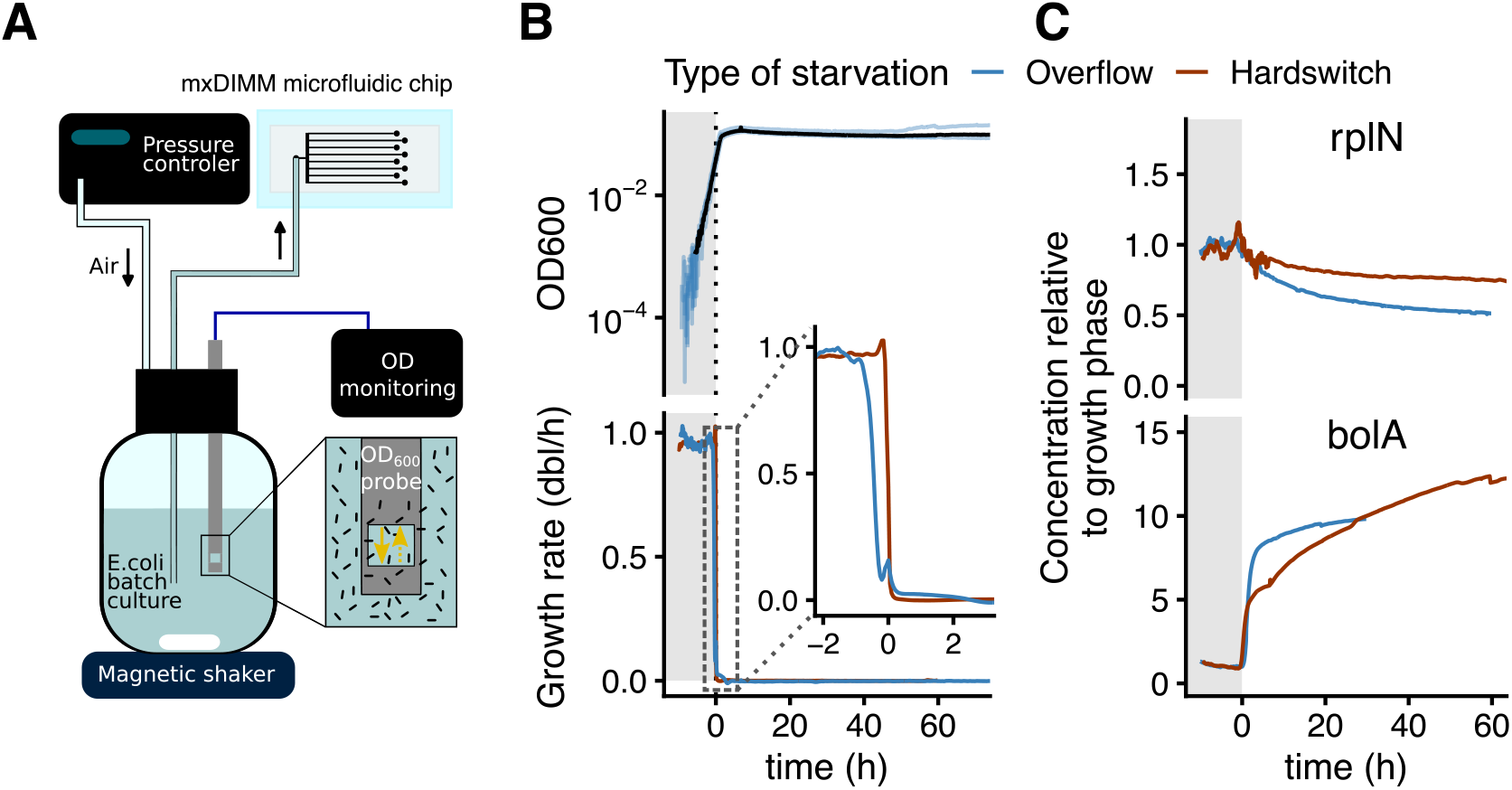
Temporal dynamics of gene expression is similar when the entry into starvation is gradual. (A) A batch culture of *E. coli* growing in M9+0.02% glucose was connected to a microfluidic chip to induce starvation of cells trapped in the growth channels of the multiplexed DIMM. (B) The OD_600_ in the flask was followed along with the average growth rate of cells in the chip to verify that the growth arrest was synchronized between flask and chip. Time 0 was defined as moment where average growth rate becomes lower than 0.055 dbl/h. Top: individual blue lines correspond to replicates, while the black line is the mean of those replicates. Bottom: average instantaneous growth rate across all cells, along time, measured in the two experimental setups, indicated with colors. The inset is a zoom on a 5 h time window spanning the entry into starvation. The growth arrest is more progressive in the overflow setup (blue line, duration ≈ 45 min; see Fig. S10A,B) than in the hard-switch setup (red line, duration ≈ 5 min; see Fig. S2B). (C) For two example promoters, the temporal dynamics and magnitude of the relative concentration, computed as in SI section 8, is similar between the two modes of entry into starvation (see also Fig. S10B and S11).

Although the growth variability across cells increases during the transition to growth arrest (Fig. S10B), which does not seem to happen when cells are switched abruptly (Fig. S2B), all cells eventually go into growth arrest, *i*.*e*. there is no subpopulation of cells which continues to grow for an extended period of time.

Despite this much more gradual entry into starvation, the dynamics of protein production rates (Fig. S10C) as well as the resulting dynamics of protein concentration (Fig. 3C and Fig. S11) are very similar to those measured in experiments with a hard switch. This suggests that our observations on gene expression in the experiments with instantaneous switching generalize to the entry into stationary phase in glucose-limited media, and probably beyond to a wide range of environmental transitions from growth to starvation.

### Gene expression changes during early starvation are highly consistent across single cells

Like the dynamics of volumic protein production (Fig. 1C), the dynamics of protein concentration is also highly consistent across single cells (Fig. 4A and Fig. S12A). To characterize the consistency in the response to starvation at the single-cell level, we quantify how much of the single-cell response is explained by the population average. For this, we measure the ratio *ρ* of the variance in the average protein concentration across time and the total variance of all protein concentration observations across all cells 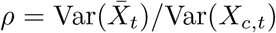, where *X*_*c,t*_ is the protein concentration in individual cell *c* at time point *t* and 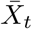 is the population average concentration at time Note that when the concentration fluctuates around a fixed value, *ρ* is close to zero, whereas in the limit that changes in the population average are much larger than single-cell variations *ρ* will approach its maximum value of 1 (Fig. 4B). Especially for promoters for which the protein concentration increases upon entry into starvation, *ρ* is much larger during the first 10 h of starvation than during exponential growth or later in starvation (Fig. 4C, Fig. S12). For bursting promoters (*e*.*g*. hslV, bolA), more than 50% of the observed variation across all cells during early starvation is captured by the population average behavior, while *ρ* is smaller for promoters only displaying a delayed arrest of production (e.g. 25% for hupA). Altogether, this analysis demonstrates that the gene expression response upon entry into starvation is highly consistent at the single-cell level, which in turn suggests that the dynamic properties of the response (i.e. the amplitudes of the burst and timescale of the arrest in volumic production for a given promoter) are tightly regulated.

**Figure 4.**
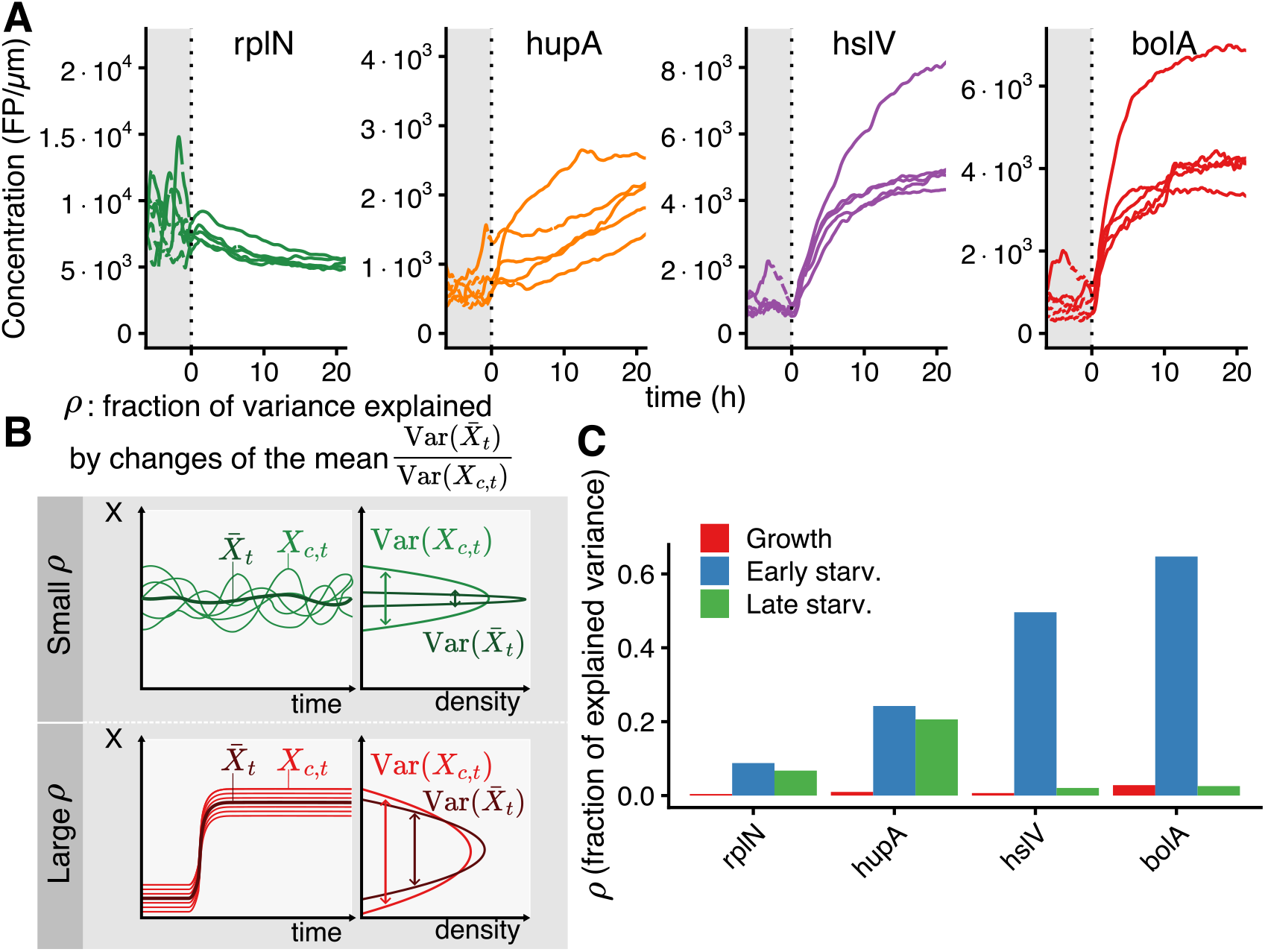
Concentrations of proteins change rapidly and homogeneously across bacterial population at starvation onset. (A) Protein concentration dynamics is highly similar across single cells and follows the average population dynamics; lines show protein concentration traces for randomly chosen single cells for 4 representative promoters. (B) To quantify the consistency of the protein concentration changes across single cells, we calculate *ρ*, the ratio of the variance in the population average protein concentration across time and the variance of all individual protein concentration measurements. Two limit cases illustrate the meaning of *ρ*. When individual cells fluctuate around a constant value, the population average variance is small compared to total variance (*ρ* ≈ 0; top row). In contrast, when all cells follow the same dynamics starting from slightly different values, the total variance is close to population average variance (*ρ* ≈ 1; bottom row). (C) The fraction of explained variance *ρ* is systematically higher during the first 10 h of starvation (“Early starv.”, blue), compared to exponential growth (“Growth”, red) and late starvation (“Late starv.”, green) behavior; corresponding values for all promoters are shown in Fig. S12.

### Protein concentrations late in starvation are set by production during early starvation

An important consequence of the fact that volumic production rates drop to low levels in late starvation is that protein concentrations deep in starvation will strongly depend on gene expression early in starvation, and perhaps even on production during exponential growth. To quantify the relative contributions from gene expression in different periods to the protein concentrations late in starvation we measured, for each time point in growth arrest, the fractions of proteins that derive from production in the exponential growth phase, during the first 10 h of starvation, and after the first 10 h of starvation. Figure 5 shows these proportions as a function of time in starvation for the same 4 example promoters displayed in previous figures. We see that for strongly downregulated promoters, such as rplN, the protein levels late in starvation are dominated by proteins produced during the exponential growth. For promoters with delayed production arrest (*e*.*g*. hslV) or with a burst (*e*.*g*. bolA), more than half of the proteins derive from production that happened during the first 10 h of starvation (blue segments in Fig. 5A). While for promoters with significant sustained expression (e.g. hupA), the proteins produced later in starvation can eventually come to constitute the majority (green area in Fig. 5A), although this is relatively rare (Fig. S13). These observations hold for all promoters displaying similar temporal dynamics of volumic production, and emerge from remarkably homogeneous behaviours across individual cells (Fig. S13).

**Figure 5.**
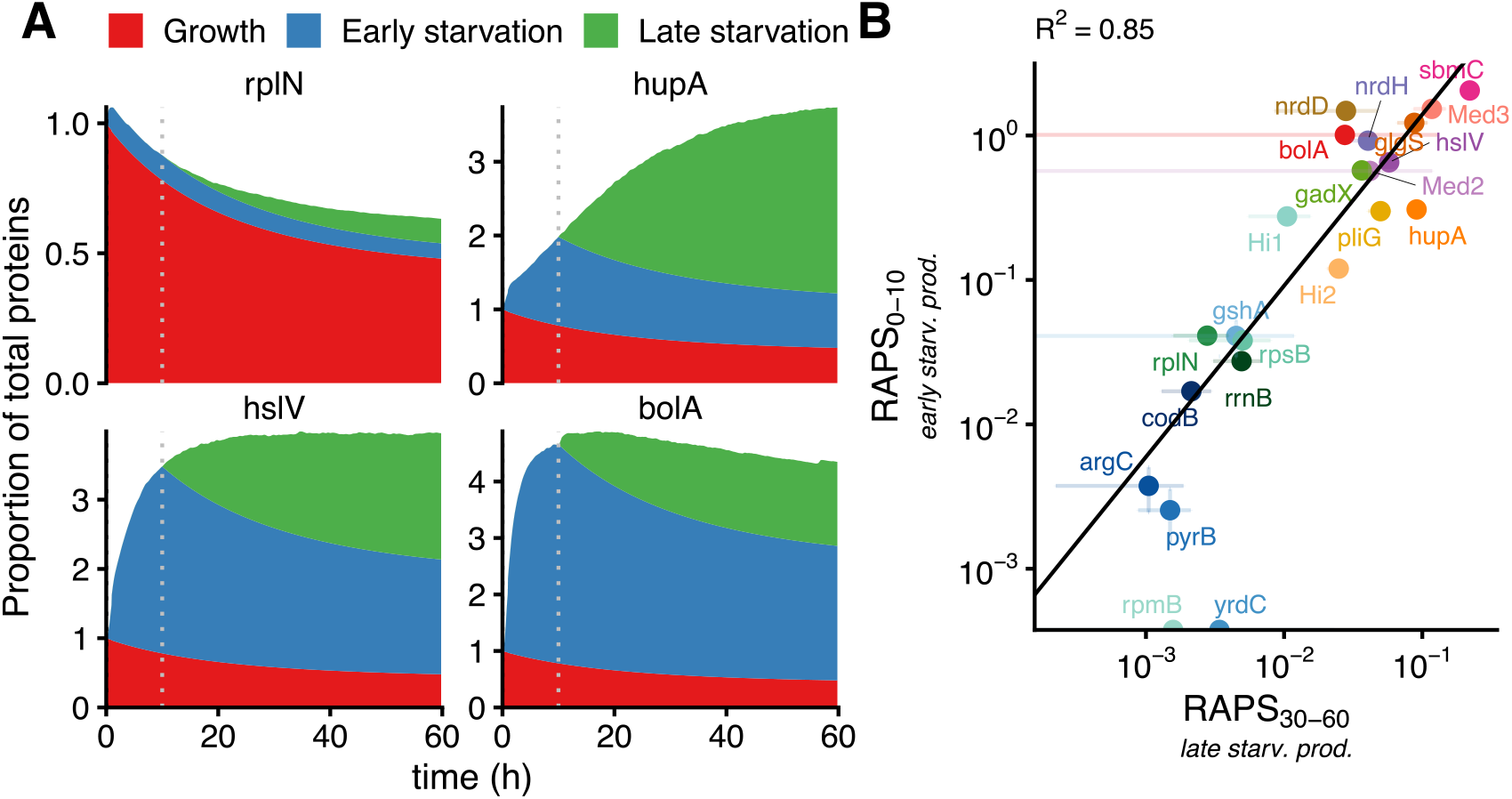
Protein concentrations late in starvation are set early on and maintained for days. (A) Amounts of total protein content coming from protein production in different time periods, as a function of time in starvation. The amount of protein coming from each time period is calculated by integrating the volumic protein production rate forward in time and accounting for protein degradation (SI section 8); “early starvation” (blue, before the vertical dotted lines) corresponds to the first 10 h of starvation while “late starvation” corresponds to 10-60 h (green, after the vertical dotted lines). The quantities are normalized by the total protein amount at time 0. (B) The level of continued protein production during late starvation, quantified by RAPS_30-60_, is correlated with the initial production during early starvation as quantified by RAPS_0-10_(Pearson correlation coefficient R^2^ = 0.85). Note that for two promoters with negligible production during early starvation, a small negative RAPS_0-10_ was estimated. These are set to 0 (bottom) and excluded from the fit.

Interestingly, we find that the ratio of the expression early in starvation (RAPS_0-10_) and the expression late in starvation (RAPS_30-60_) are strongly correlated (Pearson r^2^ = 0.85; Fig. 5B). That is, promoters that display high protein production early in starvation also tend to have sustained expression late in starvation. This suggests that the mechanism controlling gene expression during the entry into starvation might be involved in its sustainment late in starvation as well. Moreover, this correlation also implies that the changes in protein concentrations happening during the first 10 h tend to persist later on. As a consequence, the cells maintain an approximately fixed phenotypic state for days.

### The gene expression response early in stationary phase is crucial for stress tolerance later in stationary phase

We have shown that protein concentrations are remodeled early in stationary phase and that this new gene expression state is then rather stably maintained over the next two days of starvation. It is tempting to speculate that the functional role of this remodeling of gene expression early in starvation is to phenotypically prepare the cells for survival and stresses that may occur later during starvation. Indeed, as protein production rates eventually become very low, cells are likely unable to respond to environmental changes late in starvation.

To test this hypothesis, we initially aimed to investigate the role of gene expression during the first 5 h of starvation by comparing cell survival under normal conditions versus conditions in which protein translation is inhibited during the first 5 h by a low dose of chloramphenicol. However we found that, quite interestingly, the addition of chloramphenicol during the first 5 h of starvation did not prevent the burst of expression observed in RpoS regulated genes, but simply postponed it until the end of the chloramphenicol treatment (Fig. S14).

We thus decided to instead compare the effect of inhibiting protein synthesis with 16 *µ*M chloramphenicol during the entire starvation period to the effect of the same treatment starting 5 h after the entry into starvation (Fig. 6A). In the latter, only the initial production happens while the former constitutes a control where gene expression is strongly inhibited during the entire starvation period. However, we found that irrespective of translation inhibition, essentially all cells survive 1 day of starvation in our setup (Fig. S15C). Instead of testing survival to longer periods of starvation, we decided to quantify the importance of the gene expression remodeling at the onset of starvation for the stress tolerance of the cells. In particular, we compared the tolerance to a strong oxidative stress (0.65 or 0.75 mM hydrogen peroxide) from 20 to 25 h of starvation between several treatments: normal starvation, translation inhibition after 5 h in starvation, and translation inhibition during the entire starvation period (Fig. 6A).

**Figure 6.**
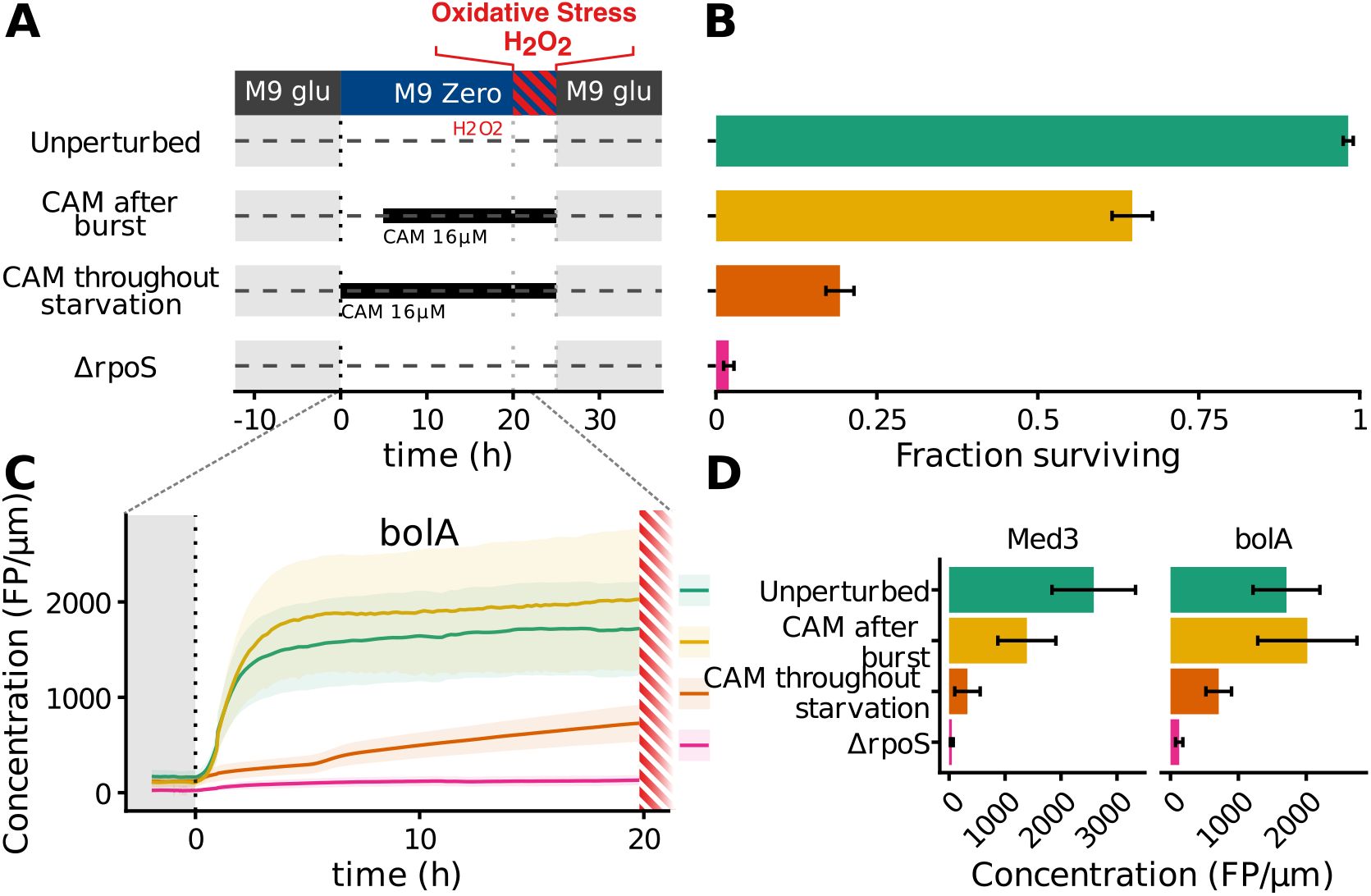
The remodeling of gene expression at the onset of starvation is essential for survival to oxidative stress later in starvation. (A) *E. coli* cells inoculated in the DIMM microfluidic chip are switched from M9 glucose to starvation in M9 zero and, after 20 h, submitted to oxidative stress during 5 h (0.65 0.75 mM H_2_O_2_), before being switched back to fresh media in order to assess survival. In addition to this default treatment (“Unperturbed”), gene expression can be inhibited using chloramphenicol, either starting after 5h of starvation (“CAM after burst”), or during the entire starvation time (“CAM throughout starvation”). Moreover, we studied a .6.rpoS mutant, in which the response to carbon starvation is strongly reduced. (B) Stress survival under the different treatments is quantified as the fraction of cells regrowing and dividing within the time of observation in fresh media (20 h). (C) Concentration profiles of the bolA fluorescent reporter (as described in Fig. 2) up to the point of the oxidative stress challenge for the various treatments (in colors). (D) Concentration levels of RpoS-regulated reporters (bolA and Med3) at the moment of the oxidative stress challenge, for the different treatments (colours). Data from 2 independent experiments and 2 similar H_2_O_2_ concentrations; replicates shown separately on Fig. S15.

We quantified stress tolerance by the fraction of cells surviving the oxidative stress, where we consider a cell to have surived if it regrows within 20 h after the termination of the stress and switch to fresh media. We find that, without translation inhibition, there is almost no mortality (98.2% survival), which indicates that most cells can handle this level of oxidative stress in absence of additional perturbations (Fig. 6B). In contrast, inhibiting gene expression with chloramphenicol during starvation increases mortality. Most importantly, the difference between the treatment with translation inhibition after 5 h in starvation and inhibiting translation from the start, demonstrates that the initial remodeling of gene expression strongly decreases the sensitivity to oxidative stress (65% *vs* 19% survival; Fig. 6B). This confirms that the remodeling of gene expression upon entry into starvation prepares the cell for stresses that can happen later during starvation.

Using the transcriptional fluorescent reporters described previously, we can also assess the effects of translation inhibition on the concentration of RpoS regulated promoters (i.e. bolA and med3; Fig. 6C and Fig. S15A,D). As expected, starting the inhibition after 5 h does not significantly affect the levels of these reporters during starvation. In contrast, upon chloramphenicol treatment during the entire starvation, their concentration is reduced more than 2-fold after 20 h. Notably, the concentrations at the time of the oxidative stress under the different treatments have the same order as the fraction surviving (Fig. 6D), suggesting that the induction of RpoS regulated genes might underlie a large part of the protective effect.

To test this further, we submitted a DrpoS strain to the same assay of resistance to oxidative stress during starvation (without translation inhibition). We find that rpoS-regulated reporters were not expressed anymore, not only during starvation but also during growth (Fig. 6C and D), strongly suggesting that their expression during growth in the wild type reflect rpoS activity during growth. Remarkably, the survival after oxidative stress dropped almost to zero and was even lower than in the wild-type strain with translation inhibition during the whole starvation period, confirming the importance of this global regulator to survive oxidative stress during starvation. The fact that deleting rpoS had a stronger impact on the sensitivity to oxidative stress than inhibiting the gene expression globally during the whole starvation could be due in part to incomplete inhibition by chloramphenicol at this concentration, but we suspect that the rpoS activity during growth, causing some expression of rpoS-regulated genes before starvation, also contributes to this effect (Fig. S9A). Overall, these experiments show that the remodeling of gene expression upon entry into starvation is crucial for resistance to stresses that may occur later in starvation.

## Discussion

Although bacteria can rapidly expand their numbers exponentially when nutrients are available, nutrients eventually always run out, forcing the bacterial cells to withstand periods of starvation, which in the wild are likely frequent and extended. Yet, very little is known about how bacteria adapt their phenotypes when faced with starvation. To address this lack of knowledge we have here tracked, with high quantitative precision and time resolution, the dynamics of growth and gene expression at the single cell level across a set of representative genes in *E coli* when cells undergo carbon starvation.

We first discuss how our findings fit in the context of previous studies. First, our main observations are that large changes in gene expression upon entry into starvation cause dramatic remodeling of proteome composition during the first 5-10 h of starvation, that this new phenotype is subsequently stable for several days, and that it is crucial for stress tolerance during starvation. These observations explain a previous report that gene expression during the first 10 h of starvation is more crucial for cell survival than expression at later times [32]. Second, it was known that, upon entry into stationary phase, the average cell size decreases due to reductive division, but this process was believed to take place as cells slow down prior to growth arrest [21, 33]. Our results show, however, that all cells immediately go into growth-arrest upon loss of the carbon source, and reductive divisions occur in this growth-arrested state (Fig. S2). In addition, we found that reductive divisions predominantly occur among cells that are further advanced in their cell cycle, confirming a previous hypothesis [22].

Third, with regard to gene expression during starvation, some previous studies have suggested that, once bacteria are growth-arrested and starving, they can no longer substantially adapt their phenotype [34]. Similarly, in another study it was proposed that adaptation to starvation only occurs when cells progressively deplete nutrients and sense the impending starvation as their growth slows down [35]. In contrast, Gefen *et al*. reported that bacteria can sustain expression, i.e. production of new proteins, over an extended period of time (≈2 days) in starvation [19], and reported an approximately constant sustained expression for a particular gene, suggesting that phenotypes can keep adapting deep into stationary phase.

Here we have shown that, contrary to the hypothesis that bacteria need to adapt their expression state before growth stops, there is dramatic remodeling of the proteome composition during the first 10 h of starvation. Moreover, the absence of dilution facilitates the increase of protein concentration for upregulated genes (Fig. 2B). Also, in contrast to the hypothesis that cells need to sense impending starvation for their adaptive response, we found that the temporal dynamics as well as the amplitude of the adaptation were not different when cells entered starvation progressively over a 45 min period as in a batch culture (Fig. 3), further highlighting the ability of bacteria to adapt during the first hours of starvation.

However, in contrast to Gefen *et al*. [19], we found that protein production systematically decreases with time and that the major changes in proteome composition occur during the first 10 h of starvation. Moreover, we show that the level of gene expression during starvation, as well as its temporal dynamics, are highly promoter-dependent, with some genes showing a transient increase in expression, and different genes exhibiting different rates of decay in protein production with time (Fig. 2B). In particular, in our sample of ≈20 promoters, the levels of protein production late in starvation (relative to that during exponential growth) span more than 2 orders of magnitude (Fig. 1E). Notably, the level of about 10% reported by Gefen *et al*. is within this range, at the upper end.

The sustained gene expression activity which we observe raises the question: how can cells maintain this energy-demanding process in the absence of a carbon source? It is known that, during starvation in *E. coli*, intracellular ATP levels drop rapidly, with 70-90% depletion occurring within minutes. The level then decreases more slowly afterwards, remaining at around 10% of the exponential growth levels for an extended period of time [11,31,36,37]. We believe that the sustained gene expression is likely fueled by different energy sources on different time scales. First, while a large fraction of protein production is allocated to production of ribosomal proteins during exponential growth [38, 39], we observe that ribosomal protein production stops immediately upon entering starvation(Fig. 1C). This sudden stop thus frees up resources that can be used for the expression of other genes during early starvation. Second, intracellular energy storage such as glycogen might transiently fuel gene expression activity very early in starvation, although it is known to be depleted within minutes at the onset of starvation [40].

It thus seems likely that the gene expression activity during prolonged starvation must be sustained mainly by endogenous metabolism [41], i.e. by the hydrolysis of abundant cellular constituents such as proteins and RNAs, which releases metabolites that can be used for producing ATP and as precursors of gene expression. Our measurements of the degradation of several fluorescent proteins during the first days of starvation (Fig. S8) shows that the rate of protein degradation *itself* decays exponentially, and that on the order of half of the proteins present at the onset of starvation are eventually degraded (Fig. 2). Assuming that other cytoplasmic proteins undergo similar degradation, up to 40% of the proteome will be degraded within the first day of starvation, which constitutes significant amounts of energy and metabolites for fueling gene expression. In addition, early experiments with mutants lacking up to 5 different proteases indicated that their activity was important for survival during sustained starvation [42], which also supports the importance of proteolysis for releasing resources during starvation. In order to further investigate the hypothesis that endogeneous metabolism fuels gene expression during starvation, it will be important to identify the proteases and nucleases active during this period in order to measure gene expression when their levels are perturbed.

It is noteworthy that our usage of microfluidic devices [20] first of all allowed us to make controlled changes in the environment and track growth and gene expression in constant starvation conditions. This is in contrast to batch cultures where, during stationary phase, bacteria continue to modify the composition of the media over time, which may have important confounding effects on the observed gene expression dynamics. Crucially, our microfluidics setup also allowed us to quantitatively track gene expression at the single-cell level. These single-cell measurements strikingly revealed that, although the common belief is that phenotypic variability is higher during stationary phase than during growth [17,18], the response to starvation which we have discovered is remarkably homogeneous across cells (Fig. 4). This homogeneity in single-cell behavior at the onset of starvation suggests the existence of a tightly controlled regulatory program that ceases energy-intensive processes and allocates available resources to reshape the cell phenotype before energy runs out completely.

By what precise regulatory mechanisms bacteria commit to this new state remains to be elucidated, but our study already provides several relevant insights. Firstly, when gene expression is inhibited with chloramphenicol during the initial 5 h of starvation, the response simply appears to occur 5 h later, and protein concentrations of induced genes still increase to levels similar to those achieved without perturbation (Fig. S14). This implies that the decrease in production rate during the first 10 h of starvation is likely the result of energy depletion by expression itself, rather than a specific regulatory mechanism at the transcriptional or translational level. Importantly, this also indicates that the total gene expression that is achieved during this period is primarily set by the amount of available intracellular resources. In order to gain more insight into how the total gene expression during starvation is set by the state of the cell when it enters growth arrest, it will likely be very informative to alter the metabolic capacity of the cells at the moment they enter growth arrest by varying the pre-starvation conditions.

The fact that different promoters show such variable time-dependent responses in protein production as a function of time raises the question as to what regulatory mechanisms implement these variable responses. For example, immediately upon entry into starvation ribosomal protein promoters are almost completely shut down. Interestingly, essentially all other promoters, including those that show a transient burst in production during the first hours of starvation, appear to also exhibit an initial very short drop in production. This suggests the existence of a general mechanism that briefly halts protein production upon entry into stationary phase. Further, our results with the knockout of rpoS (Fig. 6) show that, as expected, this sigma factor plays an important role in the expression of transiently upregulated genes. However, we suspect that other major regulators, such as the major regulator of catabolism CRP, might also be contribute to this transient upregulation.

Especially intriguing is the fact that, for promoters whose expression is sustained during prolonged starvation, the decay of the volumic production *q* happens at different rates for different promoters (Fig. S6). This indicates that the approximately exponential decay of production is not just controlled by an overall loss of expression capacity, because this would imply the production decays at the same rate for all promoters. However, we must keep in mind that the volumic production for a given promoter is the product of an overall rate of protein production and its mRNA concentration. The different rates of decay of production for different promoters thus suggest that mRNA levels continue to shift over two days of starvation. Further insight into the underlying mechanisms could perhaps be gained using dual reporters that report on the rate of production from two copies of the same promoter, or the production from two different promoters with different rates of decay. This would, for example, show whether the rates of production decay are also different within the same cell.

Perhaps the most intriguing question is what the precise functional role is of the dramatic remodeling of the proteome in the first 5-10 h of starvation. Our experiments show that the cells easily survive 25 h of starvation in our setup, even when their gene expression is inhibited throughout the whole starvation period (Fig. 6). Thus, the gene expression response is not necessary for survival on the time scale of one day, but it might well be important for survival on longer time scales. Indeed, a previous study has already reported that the first 10 h of expression during starvation are most important for survival on a longer time scale [32]. Our results further show that the gene expression occurring during the first 5 h of starvation is crucial for *E. coli* bacteria to survive a stress coming 15 h later. Thus, the gene expression response upon entry into starvation is crucial both for ensuring survival on longer time scales, and for protecting cells against stress.

In general, it might be difficult to disentangle which aspects of the expression response upon starvation are mainly to support survival and which are mainly to protect against possible stress. To learn more about how natural selection has shaped the functional role of the expression response upon entry into starvation, it would be interesting to systematically compare the response across different strains of *E. coli* as well as the differences in survival and tolerance to various types of stress across these strains.

## Acknowledgments

Data storage and image analysis calculations were performed at sciCORE (http://scicore.unibas.ch/) scientific computing center at University of Basel. Proteomics experiments were performed by the Proteomics Core Facility of the Biozentrum at University of Basel. T.G. acknowledges support from the Biozentrum Basel International PhD Program. This research was funded in part by the Swiss National Science Foundation (SNSF) [Grant 310030_197836 to T.J.], as well as by the SNF Grant PHY-1748958 and the Gordon and Betty Moore Foundation Grant 2919.02 to the Kavli Institute for Theoretical Physics (KITP).

## Notes

### Competing Interest Statement

The authors have declared no competing interest.

### Summary of Updates

ORCIDs were added for the first 2 authors.

