## Supplementary Information for "*E. coli* prepares for starvation by dramatically remodeling its proteome in the first hours after loss of nutrients"

#### 1 Bacterial strains and media

All strains used in this studies are derivatives of *E. coli* K12 MG1655 (GCSC #6300). Reporter constructs were obtained or derived from the Zaslaver library of *E. coli* native promoters [43]. Moreover, we used synthetic promoters evolved *in vitro* in the same pUA66 plasmid backbone to study the expression of unregulated promoters [24]. The sequence of promoters used in this study are listed in Table S2. We used different fluorescent reporters for different promoters; note that the same fluorescent protein is used for all strains in a given fluorescence channel, namely GFPmut2 [43], mVenus [44], and mCherry2-L [45]. All strains used in this study are listed in Table S1.

Plasmid reporters based on GFPmut2 were taken directly from the Zaslaver library. We verified that the plasmid copy number do not vary from growth to starvation, and therefore do not play a role in the observed changes of volumic production (Fig. S16C,D). Chromosomal reporters based on GFPmut2 were constructed by integrating the plasmid fragment containing the promoter, RBS, and fluorescent protein at the phage HK022 locus attB, using the *clonetegration* method [46]. The corresponding promoters are ribosomal promoters (*rplN*, *rpsB*, *rpmB*, *rrnB*), as well as two synthetic promoters (referred to as *Hi1*, *Med3*), and *gshA* and *yrdC* promoters. In addition, three strains were constructed with dual reporters (mVenus and mCerulean) for a given synthetic promoter (referred to as *Hi2*, *Med2*, *Med3*) and a mCherry reporter of the RpoS regulatory activity (using *bolA* promoter). In these strains the dual reporters are integrated with diverging orientation at the phage HK022 locus attB. In more details, each synthetic promoter was amplified from the corresponding pUA66 plasmid, together with the strong RBS sequence located downstream. Gibson assembly was used to prepare the *clonetegration* helper plasmid containing the two diverging promoter sequences separated by a random spacer, and the fluorescent proteins downstream (mVenus and mCerulean, amplified from plasmid pZS-124 [44]). Note that in the experiments reported in this article, we did not acquire the signal coming from mCerulean, but we did verify that it was not affecting other measurements (see 4.1). A similar construct of *bolA* promoter controlling the expression of a mCherry2-L protein (amplified from plasmid pEB2-mCherry2-L [45]) was integrated in these strains at the phage 186 insertion site. For all constructs integrated using the *clonetegration* protocol, the KanR resistance cassette was excised using transient FLP induction, and the final construct was checked by Sanger sequencing.

To analyse the regulatory importance of the RpoS sigma factor, $\Delta$*rpoS* strains were obtained from some of the strains mentioned above, using a P1 transduction of the *rpoS*::*kanR* loci from the Keio collection [47]. The kanamycin resistance cassette was excised afterward using the pE-FLP helper plasmid [46].

All experiments were done in M9 minimal media (Sigma-Aldrich), supplemented with 2 mM MgSO4, 0.1 mM CaCl2, and sugar as specified in the main text, typically 0.02% of glucose for the carbon limited batch culture, and 0.2% of glucose for preculture and for the growth condition in microfluidic experiments with a hard switch. To starve bacteria during these experiments, we used the same minimal media not supplemented with any carbon source (referred to as "M9 Zero"). All experiments were carried at 37°C.

| Strain | Genotype | Origin | Promoter name | Fluorescent protein | Integration site/ Plasmid |
|---|---|---|---|---|---|
| Med2 bolA | MG1655 attBHK022::(Med2p-mVenus,Med2p-mCerulean) attB186::(bolAp-mCherry2-L) | Wolf et al. 2015 Zaslaver et al. 2006 | Med2p/bolAp | mVenus/mCherry2-L | attB_HK022 attB_186 |
| Med2 bolA ΔrpoS | MG1655 attBHK022::(Med2p-mVenus,Med2p-mCerulean) attB186::(bolAp-mCherry2-L) ΔrpoS | Wolf et al. 2015 Zaslaver et al. 2006 | Med2p/bolAp | mVenus/mCherry2-L | attB_HK022 attB_186 |
| Med3 | MG1655 attBHK022::(Med3p-GFPmut2) | Wolf et al. 2015 | Med3p | GFPmut2 | attB_HK022 |
| Med3 Pl. | MG1655 | Wolf et al. 2015 | Med3p | GFPmut2 | pUA66 |
| Med3 bolA | MG1655 attBHK022::(Med3p-mVenus,Med3p-mCerulean) attB186::(bolAp-mCherry2-L) | Wolf et al. 2015 Zaslaver et al. 2006 | Med3p/bolAp | mVenus/mCherry2-L | attB_HK022 attB_186 |
| Med3 bolA ΔrpoS | MG1655 attBHK022::(Med3p-mVenus,Med3p-mCerulean) attB186::(bolAp-mCherry2-L) ΔrpoS | Wolf et al. 2015 Zaslaver et al. 2006 | Med3p/bolAp | mVenus/mCherry2-L | attB_HK022 attB_186 |
| Hi1 | MG1655 attBHK022::(pHi1-GFPmut2) | Wolf et al. 2015 | Hi1p | GFPmut2 | attB_HK022 |
| Hi1 ΔrpoS | MG1655 attBHK022::(pHi1-GFPmut2) ΔrpoS | Wolf et al. 2015 | Hi1p | GFPmut2 | attB_HK022 |
| Hi2 bolA | MG1655 attBHK022::(Hi2-mVenus,Hi2-mCerulean) attB186::(bolAp-mCherry2-L) | Wolf et al. 2015 Zaslaver et al. 2006 | Hi2p/bolAp | mVenus/mCherry2-L | attB_HK022 attB_186 |
| gshA | MG1655 attBHK022::(gshAp-GFPmut2) | Zaslaver et al. 2006 | gshAp | GFPmut2 | attB_HK022 |
| yrdC | MG1655 attBHK022::(yrdCp-GFPmut2) | Zaslaver et al. 2006 | yidCp | GFPmut2 | attB_HK022 |
| pyrB | MG1655 / pUA66-pyrBp-GFPmut2 | Zaslaver et al. 2006 | pyrBp | GFPmut2 | pUA66 |
| argC | MG1655 / pUA66-argCp-GFPmut2 | Zaslaver et al. 2006 | argCp | GFPmut2 | pUA66 |
| codB | MG1655 / pUA66-codBp-GFPmut2 | Zaslaver et al. 2006 | codBp | GFPmut2 | pUA66 |
| codB ΔrpoS | MG1655 ΔrpoS / pUA66-codBp-GFPmut2 | Zaslaver et al. 2006 | codBp | GFPmut2 | pUA66 |
| rpmB | MG1655 attBHK022::(rpmBp-GFPmut2) | Zaslaver et al. 2006 | rpmBp | GFPmut2 | attB_HK022 |
| rpsB | MG1655 attBHK022::(rpsBp-GFPmut2) | Zaslaver et al. 2006 | rpsBp | GFPmut2 | attB_HK022 |
| rpsB ΔrpoS | MG1655 attBHK022::(rpsBp-GFPmut2) ΔrpoS | Zaslaver et al. 2006 | rpsBp | GFPmut2 | attB_HK022 |
| rplN | MG1655 attBHK022::(rplNp-GFPmut2) | Zaslaver et al. 2006 | rplNp | GFPmut2 | attB_HK022 |
| rplN ΔrpoS | MG1655 attBHK022::(rplNp-GFPmut2) ΔrpoS | Zaslaver et al. 2006 | rplNp | GFPmut2 | attB_HK022 |
| rplN Pl. | MG1655 / pUA66-rplNp-GFPmut2 | Zaslaver et al. 2006 | rplNp | GFPmut2 | pUA66 |
| rrnB | MG1655 attBHK022::(rrnBp-GFPmut2) | Zaslaver et al. 2006 | rrnBp | GFPmut2 | attB_HK022 |
| rrnB ΔrpoS | MG1655 attBHK022::(rrnBp-GFPmut2) ΔrpoS | Zaslaver et al. 2006 | rrnBp | GFPmut2 | attB_HK022 |
| hupA | MG1655 / pUA66-hupAp-GFPmut2 | Zaslaver et al. 2006 | hupAp | GFPmut2 | pUA66 |
| hupA ΔrpoS | MG1655 ΔrpoS / pUA66-hupAp-GFPmut2 | Zaslaver et al. 2006 | hupAp | GFPmut2 | pUA66 |
| hslV | MG1655 / pUA66-hslVp-GFPmut2 | Zaslaver et al. 2006 | hslVp | GFPmut2 | pUA66 |
| hslV ΔrpoS | MG1655 ΔrpoS / pUA66-hslVp-GFPmut2 | Zaslaver et al. 2006 | hslVp | GFPmut2 | pUA66 |
| bolA | MG1655 / pUA66-bolAp-GFPmut2 | Zaslaver et al. 2006 | bolAp | GFPmut2 | pUA66 |
| pliG | MG1655 / pUA66-pliGp-GFPmut2 | Zaslaver et al. 2006 | ycgKp | GFPmut2 | pUA66 |
| gadX | MG1655 / pUA66-gadXp-GFPmut2 | Zaslaver et al. 2006 | gadXp | GFPmut2 | pUA66 |
| gadX ΔrpoS | MG1655 ΔrpoS / pUA66-gadXp-GFPmut2 | Zaslaver et al. 2006 | gadXp | GFPmut2 | pUA66 |
| nrdH | MG1655 / pUA66-nrdHp-GFPmut2 | Zaslaver et al. 2006 | nrdHp | GFPmut2 | pUA66 |
| nrdD | MG1655 / pUA66-nrdDp-GFPmut2 | Zaslaver et al. 2006 | nrdDp | GFPmut2 | pUA66 |
| glgS | MG1655 / pUA66-glgSp-GFPmut2 | Zaslaver et al. 2006 | glgSp | GFPmut2 | pUA66 |
| glgS ΔrpoS | MG1655 ΔrpoS / pUA66-glgSp-GFPmut2 | Zaslaver et al. 2006 | glgSp | GFPmut2 | pUA66 |
| sbmC | MG1655 / pUA66-sbmCp-GFPmut2 | Zaslaver et al. 2006 | sbmCp | GFPmut2 | pUA66 |

**Table S1.** List of strains used in this study. The paper where the constructs originate from is indicated in the column 'origin'. rpoS deletion was obtained by P1 transduction of the corresponding Keio collection strain [47]

| Promoter name | Promoter sequence |
|---|---|
| Med2p | GTTGCCCAGATCCGAGTTCGTCAGTTATAATTTGTGTATCGTTCTAGCACCACGGGTTCATCCTCTATTGCTACGGTATCGCTGCCAG |
| Med3p | CAGCGGGGTTAGTGTGCTACGCTTGATAGTGCAAAGTGGAATCGCCCTCCGATCGCTAGCCGAGGGCTTACGAGTGTTAGGTTTATTCAGGG |
| Hi1p | CAGTCAGTCCACGTGTGGCGGGAATCCCTTGGCGGCGGCCCTGCTTCTGGTGTATACTGTTATGGAAACCAAATCTACGAGCCTACAGGATTCCTATCCG |
| Hi2p | CATTCAGAGACGTAGCTGCGTGACACTGTTGTACTGGATTGTGTAAGTCTTAGGATACATCTTGTCAAGTGCTGCCTTCAGGGGTAATATAGGG |
| gshAp | CGGGATCCTCAGGGCATGATGTGGTGgcactaattgtaggcctgcacatatggtcaccattacagttatgctaattaaaacgattttgacaggcgggaggtcaatttgatcccggacgtatcacaggcgctggc<br>ctggctggaaaaacatcctcaggcgttaaaggggATACAGCGTGGGCTGGAGCTCGAGCGG |
| yidCp | CCGCTCGAGCCGACCTCGATAACCGTGctctctcggaagcgttggaaaaattatggcgccgccactgtcgcctggctcgcgggtcctgatagccctcattcgggtctatcaacgcctgattagtccgctactcgg<br>gccgcattgtcgtttcactccaacctgttcaagctacggaattgaggcattgcgcaggtttggagtgataaaaggcagttggttgacggtgaaacgcgtattaaaatgccaccctttacaccctggtggtgacg<br>atcccgtcccgcccggaccatttgataccagagaacactaacgatggattcgcaacgcaatcttttagtcatcgctttgctgttcgtgtctttcATGATCTGGCAAGCCTGGGGATCCCG |
| pyrBp | CGGGATCCGCCGTTTTTCTTCCCGTTgatcacccattcccagcccctcaatcgaggggcttttttttgcccaggcgtcaggagataaaagatggctaatccgctatatcagaaacatatcatttccataaacgac<br>cttagtcgcgatgaccttaatctggtgctggcgACAGCGGCGAAACTGAAACTCGAGCGG |
| argCp | CCGCTCGAGCTTCCGTGGCGCTTATTGaaggtgtggcaatcagagcgcggtaaatctcgataaatggcggtaatttgtttttcattgttgacacacctctggtcatgatagtatcaatattcatgcagtatttat<br>gaataaaaatacactaacgttgagcgtaataaaacccaccagccgtaaggtgaatgttttacgtttaacctggcaaccagacataagaaggtgaatagccccgatgttgaatacgctgattgtgggtgccagcg<br>gctacgctGGCGCAGAGCTAGTGACCGGATCCCG |
| codBp | CCGCTCGAGAGGACGCGATCCTGGAGatctgacgaccattgatgatcctgcgtcgttggatcagatccgccaggcgatggaagagtaggttattgtcggatgcgtcgcgcggtgcatccggcactgtgtgccgat<br>gcctgatgcgacgctgacgcgttttatcatgcctacggacctgaaccgtaggtcggataaggcgctcgcgtcgcatccgacaccatgctcagatgcctgatgcgacgctgacgcgtcttatcaggcctacccac<br>tgtttttacaccgataatttttccccccacctttttgcactcattcatataaaaaatatatttccccacgaaaacgattgctttttatcttcagatgaatagaatgcggcggatttttttgggtttcaaacagcaa<br>aaagggggaatttcgtgtcgcaagataacaACTTTAGCCAGGGGCCAGGGATCCCG |
| rpmBp | CGGGATCCTACGCGTGCTCGACCATAtcgtgattgggcgtggagagtatgtttcttttgccgaacgcggctggatttaacccgctatgcgcgatccttcgggatctttgtctgttcgggacttgagcacatcgct<br>gagtcagcgtatactacgccacctttgagaatctcgggtttggcatttgggcctggcaatcgagagttcacatagaactgcgatgaccgggctgtaaagcctgacgaggcgccaataccccatacgaagctcga<br>gctaatttgatttttggagaatagacatgtcccgagtctgccaagttactggcaagcgtccggtgaccggtaacaaccgttcccaCGCACTGAACGCGACTAACTCGAGCGG |
| rpsBp | CCGCTCGAGATCTCCAGCACTTCGGCAgccagtcggccagcgacgcgcatttttttcgatatcttctggggtcttgattgagatagccattaattctgtccatcagcgtcggtgataccgacaatatatatgtaagt<br>gccgtcaatggtatcacacccgggcaaattgagaatcattctgaatttcgccaaacgtgccactgaaggttttctataatagaaaattcgacgtctgatgctgtacacagcgccaacaattattggtgtccacga<br>cgtatttgtggtataaagcgcgccggacttccgatccatttcgtatacacagactggacggaagcgacaatctcactttgtgtaacaacacacacgtatcggcacatattccggggtgccctttggggtcggtaa<br>tatgggatacgtggaggcataaccccaacttttatatagaggttttaatcatggcaactgtttccatgcgcgacatgctcaaggctggtgttcacttcggtcaccagacccgttactggaacccgaaaatgaAGC<br>CGTTCATCTTCGGTGGGATCCCG |
| rplNp | CGGGATCCCGTCCGCTGTCCAAGACTaaatcctggacgctggttcgcgttgtagagaaagcggttctgtaatacagtacactctctcaatacgaataaacggctcagaaatgagccgtttatttttttctacccatat<br>ccttgaagcggtgttataatgccgcgccctcgatatggggatttttaacgacctgattttcgggtctcagtagtagttgacattagcggagcactaaaatgatccaagaacagactatgctgaacgtcgccgacaa<br>ctccggtgcacgtcgcgtaatgtgtaTCAAGGTTCTGGGTGGCTCTCGAGCGG |
| rrnBp | CCGCTCGAGCGCCAGGAGCTGAACAATtattgcccgttttacagcgttacggcttcgaaacgctcgaaaaactggcagttttaggctgatttggttgaatgttgcgcggtcagaaaattattttaaatttcctcttg<br>tcaggccggaataactccctataatgcgccaccactgacacggaacaacggcaaacacgccgccgggtcagcggggttctcctgagaactccggcagagaaagcaaaaataaatgcttgactctgtagcgggaagg<br>cgtattatgcacaccccgcgccgctgagaaaaagcgaagcggcactgctctttaacaatttatcagacaatctgtgtgggcactcgaagatacggattcttaacgtcgcaagacgaaaaatgaataccaagtctca<br>agagtgaacacgtaattcattacgaagtttaattctttgagcgtcaaacttttaaattgaagagtttgatcatggctCAGATTGAACGCTGGCGGGATCCCG |
| hupAp | CCGCTCGAGGCTGAGTGCGAAGAACGCgatatcgagctgataaaaggccttagggcagacctgcgtgaggcgggtgagagcaatattggtataatttttcagcaataagaccagaaaacgtgatttaacgcctgattt<br>gtcgtacctggagtcttccctttcgcccccccgtctggtctacatttgggggggcgaaaaaaagtggctatcggtgcgtgtatgcaggagagtgctattctggcatttccgtcgcactcgatgcttagcaagcgataaa<br>cacattgtaaggataacttatgaacaagactcaactgattgatgtaattgcagagaaagcagAACTGTCCAAAACCCAGGCGGATCCCG |
| hslVp | CGGGATCCAACGCAGACAGCACCCTCaatcggttgaagatggcgggtcatacaaactgcattcggctcgccgccgggggttgaaaccctcaaaatccccccccatctataattgcattatgccccgtacttttgtacggg<br>gtttgtactctgtattcgtaaccaaggggtcagctcgtgacaactatagtaagcgtacgccgtaacggccatgtggtcatcgctggtgatggtcaggcCACGTTGGGCAATACCGTCTCGAGCGG |
| bolAp | CCGCTCGAGTGGTAAAAATTCCCGCCAtcataacattgccaacggcgaggggaagtgggtaaggcatgtaaattcatcatgttgacgaaataatcgcccctggtaaaagaaacactgatgcgaggcctgtgtttcaatct<br>ttaaatcagtaaacttcatacgcttgacggaaaaaccaggacgaaacctaaatatttgttgttaagctgcaatggaaacggtaaaagcggctagtatttaaagggatggatgacatctcagcgttgtcggaggagatat<br>ttcatgatgatacgtgagcggatagaagaaAAATTAAGGGCGGCGTTCGGATCCCG |
| ycgKp | CGGGATCCGGTCTGGTCACGCTTGCTgcttcgataaatcacacaaaacatacttttactctttaagatgccaggttttggcataagctaacatcaatgaaacccgcgccacaagccagccagctgtagatttctacggttt<br>gttgagtccatgcccccttttataagtaatccccccggcaaaaatgccgcaacgtgattttaccgaacgcaggaaagccactggaattctcccgttagggattgttatattataacagttcatcgtactcattctgaacag<br>gagactaccaatgaaaatcaagagcatcaggaaggctgtattgctgcttgcactgctaacctccacGAGCTTTGCTGCGGGTAACTCGAGCGG |
| gadXp | CGGGATCCATTGCCCAGCAGAACAGCtttaaacacacctgataacataacgttgtaaaaaccgaatgcccagcctttaaaaaaaacagctgggcattcggttgcttattaatcgcaataatatattggctgtctattcatcgt<br>gttgataagatatatgttatatgaatgtttattaaatgtagcgattacctttaatgtcacgaaaagggattatttgcttactattaatttccctgtgttcttaaacacgtttttgtattataaatccagtcatcctgcccg<br>tagaacatcacacattatcatcctgttctcccgcttcgtttaaatttatttatcaatcaatttgacttaagagggcggcgtgctacattaataaacagtaatatgtttatgtaatattaagtcaactatgcaatcactaca<br>tgggaattgtctaattgcgtatgcaagacataaatatattctcaccatggttaatggtgaatatcgcTATTTTAATGGCGGTGACCTCTCGAGCGG |
| nrdHp | CCGCTCGAGGACCCTGGTGTAGCGTGGgtacagcagctgcggtaggtatttttatcggcgcactgttaagcatgcgcaaatcgtagtgcaaaaatgataataaatacgcgtctttgaccccgaagcctgtcttcggggtttc<br>tttttgcctggtgaatcacaaaaatcccccctaccccgtcacgctcatatccagggtaatttcgaccactatttgctatatattgtgtggttgaatctttttttcaactacatctagtatctctgtatcaacagagagacaac<br>ccgacgcgtatcatcgcgccgtatcttcattttaaacggaaatacgaatcatgcgcattactatttacactcgtaacgattgcgttcagtgccacgccaccaaacgggcgatggaaaacCGGGGCTTTGATTTTGAAGGATCCCG |
| nrdDp | CGGGATCCCACAACCCTGTGCCATGAatttacggccttttgaggctgtctggtggttacagaagtaaatcttccctaagccccggtaatgccggtcattccggggttttgtctgtcagtatgttgtttttgttgatttttca<br>accagcaaattcattaaaaaaatttacatatcgctgtagcgcccgtcatccgtacgctctgctttttactttgagctacatcaaaaaaaagctcaaacatccttgatgcaaagcactatatatagactttaaaatgcgtccca<br>acccaatatgttgtattaatcgactataattgctactacagctccccacgaaaaaggtgcggcgttgtggataagcggatggcgattgcggaaagcaccggaaaacgaaacgaaaaaaccggaaaacgcctttcccaattt<br>ctgtggataacctgttcttaaaaatatggagcgatcatgacaccgcatgtgatgaaacgagacggctgcaaagtgccgtttaaatcagagcgcatcaaagaagCGATTCTGCGTGCAGCTACTCGAGCGG |
| glgSp | CGGGATCCACCGCAGATAAGCGCAAAaatttctaacaggccgatcaacagtacaaaggaaatagcgaaaagataaggggtgttatagtcggcgaataaaatcatatatcactccaatgtccatttagtgacgcaatatttact<br>ttgtaatagtttccttgcttacatttatctgtgttaataataacacaggttagagaagttctggaagagaaagtcttttaatatggtttttttatttttcactaaaaagtgtgatcggggacaatatatttacgcacgttatg<br>tttaaaggcactacactgattggggaatactgaaatcagaaaaagcatatggccttaaaaggtatgcgataaccggaggatgtgcttatggatcatagtcttaattctttaaataatttcgatttcctggcgcgtagttttg<br>CCAGAATGCACGCAGAAGCTCGAGCGG |
| sbmCp | CGGGATCCTGATAAACAGGTGGCGCActggccgctggttaccctggaatctgtcggggaaggcagcatgttttctcgcctgagtgattatttccaccataaggcctgacctttcttttgcagcagactggcaggagtgcgagtc<br>tgctcgcataatcaacactcattccttgtggttttaattttgcaactatactgtatataaaaacagtatcaatggaggcgtcatgaactacgagattaagcaggaagagaaacgtaccgttgcaggtttccatctcgttggcc<br>CGTGGGAACAGACGGTAAACTCGAGCGG |

**Table S2.** Sequences of the promoters used in this study. For all native promoters coming from [43], the sequence was obtained from *in silico* PCR, using the primers provided in the articles (capital letters on the sequences)

#### 2 Microfluidic procedures

For all microfluidic experiments, bacterial cultures used for inoculation were prepared as follows, unless otherwise noted. Bacteria were streaked onto LB agar plates from frozen glycerol stocks stored at −80°C. Overnight preculture was grown from single colonies in M9 minimal medium supplemented with 0.2% of glucose. The next day, cells were diluted 100-fold into fresh medium and harvested after 4–6 h, at $OD_{600}$ 0.05-0.15.

##### 2.1 Microfluidic experiments with "hard switch" to starvation

In order to study the effect of starvation on bacterial gene expression dynamics at the single-cell level, we used a multiplexed DIMM as described previously [25]. Several independent main channels are connected to the same DaW junction, allowing to multiplex up to 8 different strains in parallel. In addition, each main channel features a low resistance overflow channel which allows loading of the strains without mixing them between channels. We used a pressure controller (OB1 mk3, Elvesys) to switch the cells from exponential growth (M9 minimal media supplemented with 0.2% of glucose) to starvation (M9 Zero).

##### 2.2 Microfluidic experiments with batch culture overflow

In order to compare the response to abrupt entry into starvation and to progressive nutrient exhaustion, we took advantage of the fact that the concentration of the limiting nutrient in a batch culture decreases due to its uptake by bacteria. When the culture reaches stationary phase, bacteria eventually stop growing after progressively exhausting nutrients. We thus continuously supplied a batch bacterial culture as media input of the microfluidic chip, such that the cells inside the chip experience the same media as the bacteria in the flask, as pioneered by others [19,21].

In more details, we seeded 10 $\mu$L of overnight culture into 100 mL of fresh M9 + 0.02% glucose. This concentration of glucose was chosen to make cell exhaust carbon before other limiting nutrients [48]. The batch culture was grown in a pressure-proof 100 mL glass bottle, with a custom cap allowing to connect tubing from the liquid to the inlet of the microfluidic chip (same as in 2.1) and to pressurise the bottle (1000 mbar during the whole experiment using an OB1 mk3 controller, Elvesys). The piece of fluoropolymer tubing used to connect the culture to the chip is chosen to minimise dead volume (ID: 0.25 mm, length ≈ 40 cm, hence ≈ 80 µL which flows in 60 to 80 min at this pressure). And the delay due to the flow should make the true difference even longer We monitor, during the whole experiment, the optical density of the culture inside the flask, using an optical fiber connected to an OD probe immersed in the liquid (Ocean Optic). The light source used for those OD measurements is a LEDD1B from Thorlabs, used at half its maximal intensity. The power is recorded using a PM100USB power meter from Thorlabs, in conjunction with Thorlabs' Power Meter Software. Because the precise time at which the transition to growth arrest happens is difficult to control experimentally, we typically ran the experiments for 120h, in order to be sure to observe at least 60h of starvation.

##### 2.3 Microfluidic experiments with oxidative stress during starvation

Applying oxidative stress at desired time during starvation and assessing survival by exposing bacteria in fresh media afterwards requires multiple media switches throughout the experiment. To do so, we took advantage of another microfluidic chip design where 8 independent DIMM are multiplexed. Those independent DIMM are similar to the original DIMM design [20], but deprived of the mixing serpentine and with all waste channels connected together for convenience. We connected to the first inlet of all the DIMM (referred to as inlets A) to tubing pieces flowing the liquid of a single reservoir through a manifold, so that the same media is flown to all. To the second inlet (referred to as inlets B), we connected tubing coming from 8 reservoirs pressurised together, so that each inlet can be flown with a different media. In order to switch four different media sequentially, the liquid of each reservoir needs to be exchanged while the corresponding inlets are inactive, as detailed in table S3.

| Current inlet | Current Media | Duration | Swap | Time of swap |
|---|---|---|---|---|
| A1 | M9 + 0.2% glucose | 17h | - | - |
| B1 | M9 Zero ±CAM | 5h | A1 → A2 | 1h |
| A2 | M9 Zero + CAM | 15h | B1 → B2 | 2h |
| B2 | M9 Zero + STRESS | 5h | A2 → A3 | 1h |
| A3 | M9 + 0.2% glucose | 30h | - | - |

**Table S3. Timing of media switches for the microfluidic experiments with oxidative stress.** The current inlet corresponds to the inlet where the highest pressure is applied, and whose media flows through the main channel. Note that in the case of B inlets, the media can be different for the 8 series. Duration corresponds to the total time cells are exposed continuously to the current media. In the example above, the experiment last in total 72 h. The column swap indicates which for which inlets the media reservoir(s) are swapped. The time of swap is indicated relative to the start of the current media.

#### 3 Microscopy and image analysis

We performed microfluidic experiments using an inverted Nikon Ti-E microscope, equipped with a motorized xy-stage and enclosed in a temperature incubator (TheCube, Life Imaging Systems). Focus was maintained using hardware autofocus (Perfect Focus System, Nikon). Images were recorded using a CFI Plan Apochromat Lambda DM ×100 objective (NA 1.45, WD 0.13 mm) and a CMOS camera (Hamamatsu Orca-Flash 4.0v3). The setup was controlled using µManager [49] and timelapse movies were recorded with its Multi-Dimensional Acquisition engine (sometimes customized using runnables). Phase contrast images were acquired using 100 ms exposure (Thorlabs MWWHL4, at 50% power). Images of mVenus fluorescence and mCherry fluorescence were acquired using a 459/526/596 nm triple-edge dichroic beamsplitter and 475/543/702 nm triple-band emission filter (Semrock), using 200ms of exposure time and 32 and 8% of excitation intensity, respectively (Lumencor SpectraX, Teal LED, excitation filter 511/16 nm, and Yellow LED, excitation filter 578/21 nm). The use of the multiband filter allows to acquire both fluorescent signals without changing the filter sets, which increases the number of positions which can be measured at a given acquisition frequency. GFPmut2 fluorescence was acquired using a 495/605 dual-edge dichroic beamsplitter, and a 527/645 dual-band emission filter (Semrock), using 200ms of exposure time and 8% of excitation intensity (Lumencor SpectraX, Cyan LED, excitation filter 470/24 nm). In growth media, images were acquired every 3 min, and every 12 min only in growth arrest conditions, in order to minimize phototoxicity. Note that acquisition was still performed every 3 min during the first hour after the switch to starvation media in order to monitor transient dynamics with high time resolution. In those conditions, the ability of cells to regrow when exiting starvation was not affected when exposed to 30h of starvation (100% of cells regrow, see Fig. S15C), but we observe a substantial increase in the non regrowing fraction after 60h of starvation, as well as a detectable increase of time to regrow (Fig. S16A,B). Even though the absence of sudden drop of fluorescent content indicates an absence of cell lysis, those non regrowing state might be associated with differences in the gene expression ability of the cells, and might therefore affect late starvation observations. Image analysis was performed using MoMA [20]. For datasets acquired in the later part of this project, we used a new version of the software developed internally (available at https://github.com/nimwegenLab/moma/releases/tag/v0.9.6, and relevant segmentation models at https://github.com/nimwegenLab/moma-model). The main difference is that segmentation hypotheses are generated using probability maps produced by a U-Net segmentation model trained for this purpose; cost functions used for tracking by global optimization were adapted as necessary. Raw images from the experiments were stored on a centralised storage, and preprocessed before being fed to MoMA. Empty growth channels, as well as those featuring structural defects or cells being growth-arrested from the start, were excluded from the analysis. Summary of the final number of cells observed during the experiments for each strains can be found in Table S4.

### 4 Estimation of fluorescent proteins levels

#### 4.1 Accounting for autofluorescence and crosstalk between fluorescence channels

For each fluorescent protein, we measured the corresponding levels of auto-fluorescence per $\mu$m cell length of our wild type strain in the same experimental conditions as used for gene expression estimation. The estimated auto-fluorescence per unit length was found to only depend on the nature of the fluorescent protein (mCherry, GFP, mVenus), and not on the conditions (growth or starvation). In our analysis, we always first subtract the auto-fluorescence from the measured fluorescent levels.

In addition, since we use multi-band fluorescence filters and since the excitation spectra of mCherry2 overlaps with the excitation spectra of mVenus, and similarly between mVenus and mCerulean, we checked whether the readout in one channel was affected by the level of the other when using dual fluorescent reporters. With *E. coli* strains transformed with a pTrc99a plasmid containing either mVenus, mCherry2, mCerulean, we could measure the amount of fluorescence coming from mVenus protein in the mCherry2 channel or mCerulean protein in the mVenus channel, and reversely. As expected from the spectra, we found no contamination coming from mVenus signal into the mCherry2 channel,

| Experiment | Strain | #Growth cell | #Starv. cell | #Transition cell | #Replicates |
|---|---|---|---|---|---|
| Hardswitch | Med2 bolA | 3745 | 496 | 307 | 4 |
| Hardswitch | Med2 bolA ΔrpoS | 617 | 79 | 50 | 1 |
| Hardswitch | Med3 | 3912 | 457 | 269 | 3 |
| Hardswitch | Med3 Pl. | 3851 | 510 | 2977 | 3 |
| Hardswitch | Med3 bolA | 5434 | 754 | 455 | 5 |
| Hardswitch | Med3 bolA ΔrpoS | 536 | 73 | 45 | 1 |
| Hardswitch | Hi1 | 1419 | 178 | 101 | 1 |
| Hardswitch | Hi1 ΔrpoS | 1339 | 149 | 103 | 1 |
| Hardswitch | Hi2 bolA | 7287 | 974 | 592 | 5 |
| Hardswitch | gshA | 790 | 106 | 63 | 1 |
| Hardswitch | yrdC | 710 | 92 | 66 | 1 |
| Hardswitch | pyrB Pl. | 3068 | 383 | 222 | 2 |
| Hardswitch | argC Pl. | 2862 | 325 | 208 | 2 |
| Hardswitch | codB Pl. | 2895 | 284 | 216 | 2 |
| Hardswitch | codB Pl. ΔrpoS | 1335 | 144 | 96 | 1 |
| Hardswitch | rpmB | 1934 | 230 | 137 | 2 |
| Hardswitch | rpsB | 707 | 82 | 57 | 1 |
| Hardswitch | rpsB ΔrpoS | 2489 | 284 | 189 | 2 |
| Hardswitch | rplN | 2244 | 314 | 195 | 2 |
| Hardswitch | rplN ΔrpoS | 376 | 61 | 28 | 1 |
| Hardswitch | rplN Pl. | 1027 | 131 | 85 | 1 |
| Hardswitch | rplN Pl. ΔrpoS | 2025 | 330 | 173 | 1 |
| Hardswitch | rrnB | 726 | 100 | 57 | 1 |
| Hardswitch | rrnB ΔrpoS | 723 | 92 | 64 | 2 |
| Hardswitch | hupA Pl. | 3355 | 413 | 225 | 2 |
| Hardswitch | hupA Pl. ΔrpoS | 1786 | 178 | 129 | 1 |
| Hardswitch | hslV Pl. | 3582 | 468 | 254 | 2 |
| Hardswitch | hslV Pl. ΔrpoS | 370 | 131 | 31 | 1 |
| Hardswitch | bolA Pl. | 2072 | 237 | 156 | 2 |
| Hardswitch | pliG Pl. | 1928 | 259 | 132 | 2 |
| Hardswitch | gadX Pl. | 3506 | 512 | 255 | 2 |
| Hardswitch | gadX Pl. ΔrpoS | 1024 | 88 | 71 | 1 |
| Hardswitch | nrdH Pl. | 3278 | 459 | 233 | 3 |
| Hardswitch | nrdD Pl. | 1839 | 272 | 136 | 2 |
| Hardswitch | glgS Pl. | 2248 | 289 | 156 | 2 |
| Hardswitch | glgS Pl. ΔrpoS | 1606 | 197 | 133 | 1 |
| Hardswitch | sbmC Pl. | 2480 | 330 | 175 | 2 |
| Overflow | Med2 bolA | 364 | 51 | 27 | 1 |
| Overflow | Med3 bolA | 486 | 77 | 38 | 1 |
| Overflow | Hi2 bolA | 2325 | 387 | 209 | 2 |
| Overflow | rpmB | 1112 | 204 | 97 | 1 |
| Overflow | rplN | 1147 | 207 | 104 | 1 |
| Overflow | rrnB | 1068 | 197 | 100 | 1 |
| Stress: Unperturbed | Med3 bolA | 2540 | 657 | 203 | 2 |
| Stress: CAM after burst | Med3 bolA | 7197 | 492 | 284 | 2 |
| Stress: CAM thr. starv. | Med3 bolA | 9769 | 623 | 289 | 2 |
| Stress: Unperturbed | Med3 bolA ΔrpoS | 2628 | 1044 | 256 | 2 |

**Table S4.** Summary of cell numbers in the different experiments. Transition cells correspond to cells undergoing the switch from M9+0.2% glucose to M9 zero and no contamination coming from mCerulean into the mVenus channel. Nevertheless, we found that the mCherry2 signal appeared in the mVenus channel at 21% of its level in the mCherry2 channel. We thus compensate for this, by subtracting this fraction of mCherry signal from the total mVenus signal, in the cases where we used dual fluorescent reporter strains. It is important to note that, due to the much higher intensity obtain with mVenus, compared to mCherry in a given experiment, the contribution of mCherry in the mVenus signal was always very small.

##### 4.2 Fluorescence signal conversion to absolute numbers

Throughout the article, we express the fluorescence levels in units of number of fluorescent proteins $n$. To obtain this number $n$, we need to convert the autofluorescence-corrected fluorescence intensity $I$ we get from our measurements (as measured by our setup, using specific illumination parameters) using a conversion factor $\alpha$, corresponding to the fluorescence emitted by a single fluorescent protein, i.e. $n = I/\alpha$.

For GFP, we have previously estimated $\alpha$ from the fluctuations of fluorescence levels in newborn sibling pairs [20]. We used this estimate $1/\alpha_{\text{Kaiser}} = 0.0361$ to infer the conversion factors in our experiments as well. In order to account for the fact that the illumination parameters of our experiments are not always the same, we assumed that $\alpha$ scales linearly with the light dose $D = T \times L \times N$ where $T$ is the exposure time in ms, $L$ is the excitation power of the light source and $N$ is the attenuation factor of the (facultative) neutral density present in the excitation light path. We therefore have, for GFP in our experiment

$$\alpha_{\text{GFP}} = \frac{D_{\text{GFP}}}{D_{\text{Kaiser}}}\alpha_{\text{Kaiser}}$$

Given our illumination settings (see SI section 3), we have $D_{\text{GFP}} = 200 \times 0.08 \times 1 = 16$ instead of $D_{\text{Kaiser}} = 2000 \times 0.16 \times \frac{1}{4} = 80$. We can thus use the conversion factor $1/\alpha_{\text{GFP}} = 0.1805$ for GFP in our experiments.

To find the conversion factor for the other fluorescent proteins (mCherry noted RFP, mVenus noted YFP), we assumed that, at steady state exponential growth, the number of fluorescent proteins present in the cells is dependent on the promoter, but independent of the reporter protein. We took advantage of sets of experiments where the same promoter drives the expression of different reporter fluorescent proteins, and write $I_{\text{RFP/YFP}}/\alpha_{\text{RFP/YFP}} = n = I_{\text{GFP}}/\alpha_{\text{GFP}}$, which in turn means

$$\frac{1}{\alpha_{\text{RFP/YFP}}} = \frac{I_{\text{GFP}}}{I_{\text{RFP/YFP}}}\frac{1}{\alpha_{\text{GFP}}}$$

Note that we first need to convert the intensity to single-reporter equivalents if one of the two reporters is expressed from a plasmid. This is due to the fact that plasmids are present in multiple copies per cell, and therefore we expect $n$ to be higher in those strains compared to chromosomal reporter strains, by a factor equal to the plasmid copy number. Using the plasmid copy number estimated in Fig. S16D, we find $1/\alpha_{\text{RFP}} = 0.495$ and $1/\alpha_{\text{YFP}} = 0.105$.

We stress that these estimations of the conversion factors $\alpha$ have limited accuracy. The main aim of reporting expression levels in protein number equivalents, as opposed to using 'arbitrary units', is that it provides a reasonable indication of the order of magnitude of the absolute number of proteins present in the cell. The relative expression levels across time of the reporters are of course not affected at all by this potential inaccuracy in the estimate of absolute levels.

#### 5 Gaussian process prior inference on the data

Our experiments provide time course measurements of total fluorescence and size of each cell in the microfluidic device, and from these measurements, we want to infer the instantaneous growth rates and volumic protein production rates of each cell as a function of time. In particular, each of the fluorescence and cell size measurements is affected by measurement noise and we want to take these into account in our inference of instantaneous growth and protein production rates, and rigorously assign error-bars to the inferred production and growth rates.

To do this, our lab has developed a Bayesian procedure which uses Gaussian process priors for both the volumic production rate and growth rate. It is implemented as a software called RealTrace, available and documented at `https://github.com/nimwegenLab/RealTrace`. The basic idea behind RealTrace is to only assume that both the growth rate and the instantaneous volumic protein production rate vary smoothly and are characterized solely by their means, variances, and correlation time. RealTrace uses a maximum entropy prior for the time-dependent growth and volumic production rates conditioned on these parameters, and additionally the (unknown) size of the noise of the fluorescence and size measurements. RealTrace first optimizes the parameters of the prior and measurement errors, i.e. finding the parameters that maximize the likelihood of all measurements in a given experiment, and then uses this maximum likelihood prior to calculate the posterior distributions over cell sizes, fluorescence concentration, instantaneous growth rate, and instantaneous volumic production rates for each cell at each time point.

The maximum entropy (i.e. Gaussian process) priors assume that key parameters such as growth and volumic production rate fluctuate around fixed averages during the experiment. For experiments, such as ours, in which growth conditions are changed dramatically during the experiments, causing large changes in the distribution of instantaneous growth and production rates, the accuracy of RealTrace inferences can be improved by using priors with different parameters for different time segments of the experiment. RealTrace therefore allows users to specify different parameters of the prior for different time segments of the experiment.

For each promoter, we used this feature of RealTrace to use one set of prior parameters for the exponential growth phase and one for the stationary phase. The prior's parameters for the exponential phase were simply fitted from all data of the experimental growth phase of the experiment while for the starvation phase, we used a slightly more complicated procedure to obtain the parameters of the prior. In particular, we observed that due to the fast and dramatic changes in volumic production at the start of the starvation phase, the algorithm had difficulty distinguishing measurement noise from rapid changes in volumic production, leading to unstable estimates of the measurement noise size. We thus first estimated the size of measurement errors, as well as the growth rate parameters, by fitting the parameters of the prior only on the later parts of the starvation phase, and then refitted all other parameters of the prior on the data from the entire starvation period.

Using these priors we then obtain from RealTrace posterior distributions for the fluorescence level in the individual cells at every time point, as well as their length, growth rate, and fluorescent protein volumic production rate. These results are the data that we use throughout this study. Finally, we found that a small fraction of data points (approximately 10 000 out of 5 682 000 data points, i.e $< 0.2\%$) were affected by segmentation errors in the DeepMoMA image analysis step that escaped attention during curation (*e.g.* two cells merging into one), and we filtered these out by hand.

### 6 Quantification of fluorescent proteins degradation

#### 6.1 Experimental protocol

We set out to measure the rate of degradation of our fluorescent proteins by performing experiments at very low exposure frequency, so as to avoid a fluorescence loss due to photobleaching. For this, we used *E. coli* strains transformed with a pTrc99a plasmid containing either mVenus, mCherry2-L, GFPmut2 or mCerulean under the control of an IPTG-inducible promoter. We conducted mother machine experiments with those strains, switching them from M9+0.2% glucose supplemented with 500 µM IPTG to M9zero (i.e. without IPTG) and back. During starvation, we acquired fluorescence images only every 3 or 6 hours, thus making the effect of photo-bleaching negligible. The decrease of fluorescence signal can therefore be assumed to be only due to protein degradation (mediated or not by proteases). Total fluorescence of single cells was recorded in this way for 30 and 60 h, respectively, with 3 hours intervals for the 30 hours time course and 6 hours intervals for the 60 hours time course, so that we typically had 10 fluorescence frames per single cell per experiment (see Fig. S8A for example time traces).

#### 6.2 Estimation of degradation rates

For each cell and each consecutive pair of time points we estimated the degradation rate by taking the discrete derivative in log-fluorescence and then averaged these across all cells at each time point. In Fig. S8B, we can see that the resulting estimated decay rates are decreasing approximately exponentially with time during the starvation period (i.e. decreasing approximately linearly on a logarithmic scale).

The degradation rate $\gamma$ is thus a time-dependent variable during starvation that decreases approximately as $\gamma(t) = \gamma_0 e^{-\omega t}$ where $\gamma_0$ is the degradation rate at entry into starvation and $\omega$ is the exponential decay rate of $\gamma$. These parameters are obtained from exponential fits of the instantaneous derivative of fluorescence along time (Table S5).

| Fluo | $\gamma_0(\mathrm{h}^{-1})$ | $\omega(\mathrm{h}^{-1})$ |
|---|---|---|
| mCherry | 0.0215 | 0.029 |
| GFP | 0.028 | 0.034 |
| mVenus | 0.0215 | 0.052 |

**Table S5.** Parameters of the time-dependent protein degradation rates $\gamma(t) = \gamma_0 e^{-\omega t}$ as estimated from control experiments with IPTG-inducible promoters.

### 7 Quantification of photobleaching rates

#### 7.1 Experimental protocol

We measured photobleaching rates of the different fluorescent proteins using the following protocol. We loaded *E. coli* cells transformed with one of the pTrc99a plasmids containing fluorescent reporters under the control of an IPTG-inducible promoter described above into the mother machine device. We then induced the expression of the fluorescent proteins with a media containing M9+0.2% glucose and IPTG 500 µM. Subsequently we switched the cells to media without inducer, either M9+0.2% glucose (to measure photobleaching during exponential growth), or M9 zero (to measure photobleaching during stationary phase). We measured the fluorescence as a function of time, using our default illumination settings (see 3).

#### 7.2 Estimation of photobleaching rates

Using the results from the above-mentioned experiments, we estimated the photobleaching rates for our different fluorescent proteins (mCherry, mVenus, GFP) in the different experimental conditions (exponential growth and starvation settings). Because the total decay in fluorescence is the result of both photobleaching and degradation of the fluorescent proteins, we need to account for the protein degradation in order to estimate the photobleaching rate. Once this is done, we use a Bayesian inference procedure to estimate the most likely photobleaching rate across all observations. This value is ultimately the one we use to run RealTrace on our data.

In order to quantify photobleaching from fluorescent traces acquired in these experiments, we calculate and subtract at every time point the amount of protein that has been degraded from the start of the observation.

The total fluorescence decreases according to the differential equation

$$\frac{dF(t)}{dt} = -\left(\beta + \gamma(t)\right) F(t),$$

with $\gamma(t) = \gamma_0 e^{-\omega t}$. The solution of this differential equation is

$$F(t) = F_0 e^{-\frac{\gamma_0}{\omega}(1-e^{-\omega t})-\beta t} \tag{S1}$$

We define $F_\beta(t)$ as the amount of fluorescence at time $t$ in the absence of protein degradation, i.e. if fluorescence decayed only due to photobleaching. Using equation S1 we have

$$F_\beta(t) = F(t) e^{\frac{\gamma_0}{\omega}(1-e^{-\omega t})} \tag{S2}$$

so that $F_\beta(t) = F_0 e^{-\beta t}$. We then use the fluorescence values $F_\beta(t)$ computed with Eq. S2 to estimate the photobleaching rate.

In particular if we define:

- $t_{ci}$ the time at which the $i$th measurement of cell $c$ took place.
- $f_{ci} = \log(F_\beta(t_{ci}))$ the logarithm of the degradation-corrected measured fluorescence value in cell $c$ at time $t_{ci}$.
- $\phi_c$ the initial log fluorescence of cell $c$.
- $\alpha$ the photobleaching rate.
- $\sigma$ the noise of the log-fluorescence measurement.

then the probability of all the fluorescence measurements given $\alpha$, the $\phi_c$ and $\sigma$ is given by

$$
\begin{aligned}
P(D|\boldsymbol{\phi_c}, \alpha, \sigma) &= \prod_{c=1}^{C} \prod_{i=1}^{N_c} \frac{1}{\sigma\sqrt{2\pi}} \exp\left[-\frac{1}{2}\left(\frac{f_{ci} - \phi_c + \alpha t_{ci}}{\sigma}\right)^2\right] \\
&\propto \frac{1}{\sigma^N} \exp\left(-\frac{1}{2}\sum_{c,i}\left(\frac{f_{ci} - \phi_c + \alpha t_{ci}}{\sigma}\right)^2\right),
\end{aligned} \tag{S3}
$$

where $\boldsymbol{\phi_c}$ is the vector of $\phi_c$, $C$ is the number of cells, $N_c$ is the number of data points in cell $c$, and $N = \sum_c N_c$ is the total number of data points in our data set.

Marginalizing over all $\phi_c$ using uniform priors and over $\sigma$ using a scale-prior $P(\sigma) = \frac{1}{\sigma}$ we obtain:

$$
P(D|\alpha) \propto \left(\sum_{c=1}^{C} N_c \text{var}(f_c + \alpha t_c)\right)^{-\frac{N-C}{2}},
$$

where $\text{var}(f_c + \alpha t_c)$ denotes the variance of the quantity $f_{ci} + \alpha t_{ci}$ across the $N_c$ timepoints of cell $c$.

The optimal bleaching rate $\alpha_*$ and its error-bar (i.e. standard-deviation of the posterior) $\sigma_\alpha$ can be expressed in terms of the weighted sums of variances

$$
V_f = \sum_{c=1}^{C} N_c \text{var}(f_c), \tag{S4}
$$

$$
V_t = \sum_{c=1}^{C} N_c \text{var}(t_c), \tag{S5}
$$

and the weighted sum of covariances

$$
V_{ft} = \sum_{c=1}^{C} N_c \text{var}(f_c, t_c). \tag{S6}
$$

| Fluo | $c_{step}$ | $\beta(\mathrm{h}^{-1})$ | Correction coeff. |
|---|---|---|---|
| GFP | Growth | 0.026 | 1.04 |
| mCherry | Growth | 0.20 | 1.29 |
| mVenus | Growth | 0.28 | 1.4 |
| GFP | Starvation | 0.0041 | - |
| mCherry | Starvation | 0.031 | - |
| mVenus | Starvation | 0.044 | - |

**Table S6.** Experimentally measured bleaching rates and corresponding correction constant to estimate the total concentration from the observed concentration during exponential growth

In particular, we find

$$\alpha^* = -\frac{V_{ft}}{V_t} \tag{S7}$$

with error-bar

$$\sigma_\alpha = \sqrt{\frac{V_f V_t - V_{ft}^2}{V_t^2(N-C)}}. \tag{S8}$$

The resulting photobleaching rates $\alpha_*$ used for the inference with RealTrace are reported in Table S6.

#### 8 Calculation of production traces and concentration traces

##### 8.1 Inference of true instantaneous production

To infer a volumic production rate $q_{\inf}(t)$, RealTrace assumes that the total fluorescence $F(t)$ of the cell evolves according to

$$\frac{dF(t)}{dt} = q_{\inf}(t)V(t) - \beta F(t), \tag{S9}$$

where $\beta$ is the inferred bleaching rate. That is, RealTrace neglects the loss of fluorescence due to degradation of (unbleached) fluorescent proteins. The true volumic production $q(t)$ evolves according to

$$\frac{dF(t)}{dt} = q(t)V(t) - (\beta + \gamma(t))F(t), \tag{S10}$$

where $\gamma(t)$ is the time-dependent degradation rate that we inferred as described in section 6. Comparing these two equations we see that true production is given in terms of the inferred production as

$$q(t) = q_{\inf}(t) + \gamma(t)\frac{F(t)}{V(t)}. \tag{S11}$$

We use this equation to infer the true production $q(t)$ as shown in, *e.g.*, Fig. 1C and Fig. S5, using the estimates of $q_{\inf}(t)$, $F(t)$ and $V(t)$ given by RealTrace, and the degradation rate $\gamma(t)$ that we estimated in section 6. Finally, we note that for observations during the exponential growth phase we assume that the degradation rate $\gamma(t)$ equals the degradation rate $\gamma_0$ at the start of the starvation phase. Notably, since in exponentially growing cells the role of degradation is negligible compared to dilution due to growth, we expect the relative error introduced by this assumption to be very small.

##### 8.2 Integration of concentration traces

We want to calculate the true concentration of proteins $C(t)$ as a function of time $t$ in stationary phase, i.e. as plotted in Fig. 2A and Fig. S12A. Notably, this concentration includes both unbleached and bleached proteins. For that, we need to integrate the concentration dynamics consisting of the true volumic production at rate $q(t)$ and protein degradation at rate $\gamma(t)$ during stationary phase. The dynamics reads,

$$\frac{dC(t)}{dt} = q(t) - \gamma(t)C(t).$$

Here, $q(t)$ is the true production rate which is corrected for bleaching and degradation as described in the last section.

**Correcting the concentration for photobleaching during exponential growth** Note that this integration approach requires estimating the total concentration of fluorescent proteins $C(0)$ at the start of stationary phase $t = 0$, i.e. including both unbleached and bleached proteins. However, our measurements only give us estimates of the concentration $C_u(0) = F(0)/V(0)$ of unbleached proteins at the start of stationary phase. To estimate $C(0)$, we thus need to estimate the ratio of bleached and unbleached proteins at the start of stationary phase. Note that during exponential growth, the concentration $C(t)$ evolves according to

$$\frac{dC(t)}{dt} = q(t) - (\lambda(t) + \gamma_0)C(t), \tag{S12}$$

where $\lambda(t)$ is the instantaneous growth rate and $\gamma_0$ is the degradation rate during exponential growth. Similarly, the concentration of unbleached proteins $C_u(t) = F(t)/V(t)$ evolves according to

$$\frac{dC_u(t)}{dt} = q(t) - (\lambda(t) + \gamma_0 + \beta)C_u(t). \tag{S13}$$

Both the volumic production $q(t)$ and the growth-rate $\lambda(t)$ will fluctuate around averages $q$ and $\lambda$ respectively, and if we approximate them as approximately constant, we find steady-state values $C = Q/(\lambda + \gamma)$ and $C_u = Q/(\lambda + \gamma_0 + \beta)$. We thus find that approximately,

$$C = C_u \frac{\lambda + \gamma_0 + \beta}{\lambda + \gamma_0} \approx C_u \frac{\lambda + \beta}{\lambda}, \tag{S14}$$

where the last approximation follows from the fact that the degradation rate $\gamma_0$ is much smaller than the dilution rate $\lambda$. With an average growth rate of approximately $\approx 0.7$ h$^{-1}$ and the bleaching rate estimated as described in the previous section, we obtain the correction coefficients displayed in Table S6.

We use this correction to estimate the concentration at the beginning of the cell cycles during growth and at the first observation in starvation. In the case of reductive division, where the cell cycle starts after the beginning of growth arrest, we cannot use this factor. In this case, we simply use the last (corrected) concentration of the parent cell as a starting observation.

**Integrating instantaneous production from initial fluorescence value**
In order to obtain the level of proteins in the cells at time $t$, integrating the true production and taking into account the degradation of both already-bleached and unbleached proteins, we need to integrate

$$\frac{dC(t)}{dt} = q(t) - \gamma(t)C(t) \tag{S15}$$

There is no easy analytical solution, and we therefore decided to integrate it numerically instead, using the following iterative scheme :

$$C(i+1) = C(i) + [q(i) - \gamma(i)C(i)]\,\Delta_t(i) \tag{S16}$$

At time $t = 0$, we use the value $C(0)$ calculated from the bleaching correction coefficient if the cells are coming from exponential growth (see Table S6), or the last calculated concentration of the parent cells if they come from a reductive division during starvation.

#### 8.3 Dynamics of the protein amount produced during a given time window

We now want to estimate at each time point the fraction of fluorescent proteins coming from *e.g.* a burst of expression (during the entry in starvation). For this, we need to track the evolution of a pool of proteins produced during a given time window. As in the previous section, this requires to rewrite the dynamics of $F(t)$ in terms of the true production and degradation, rather than using the rate $q_{\text{inf}}$.

As for equation S15, we want to integrate the following:

$$\frac{dF(t)}{dt} = q(t)V(t) - \gamma(t)F(t) \tag{S17}$$

We do not use an analytical solution, but, similar to S16, we integrate it numerically instead, using an iterative scheme:

$$F(i+1) = F(i) + [q(i)V(i) - \gamma(i)F(i)]\,\Delta_t(i) \tag{S18}$$

From this, we can calculate, at any given time point, the amount of protein coming from a given period of time $t_1 \to t_2$. At time $t$, the amount of protein that remains and has been produced during this period of time $F_{1\to 2}(t)$ can be calculated with :

$$F_{1\to 2}(i) = \begin{cases} 0 \quad ; \quad i < t_1 \\ F(i) + [q(i)V(i) - \gamma(i)F(i)]\,\Delta_t(i) \quad ; \quad t_1 \leq i < t_2 \\ F(i) - \gamma(i)F(i)\Delta_t(i) \quad ; \quad i \geq t_2 \end{cases} \tag{S19}$$

We can now follow, the fraction $F_{\text{frac}}(t)$ of the total protein at time $t$ that have been produced between $t_1$ and $t_2$.

$$F_{\text{frac}}(t) = \frac{F_{1\to 2}(t)}{F(t)}$$

Calculating this fraction for three time windows : Growth ($t \leq 0$ h), Early Starvation (0 h $< t \leq 10$ h), and Late Starvation ($t > 10$ h), we obtain the Muller plots represented on Fig. 5A and Fig. S13.

#### 9 Dynamics of volumic production in late starvation

In contrast to the previously reported Constant Activity in Stationary Phase, we observe no period during starvation where gene expression remains constant. From the maximal level of production (either the level during exponential growth or the top of the peak when there is a burst), volumic production initially decreases fast for all promoters, after which it decreases more slowly. The overall dynamics of volumic production over time is reasonably well fitted by a bi-exponential decay (Fig. S6B, fitting procedure described below), although this fit does not extend late in starvation for most promoters because the production is estimated to be negative at some time points due to measurement errors. Note that when production is very low, fluctuations of the autofluorescence and of the degradation of fluorescent proteins, as well as measurement errors of the total fluorescence level, will all visibly affect the accuracy of the estimation of the instantaneous production.

**Exponential fit of the production decay** In order to fit the decay of production with a bi-exponential function, we express $q_i$, the "true" $q$ at time $t_i$, as

$$q_i = (b-a)e^{-\lambda_1 t_i} + ae^{-\lambda_2 t_i}$$

which can be rewritten

$$q_i = q_0 \left((1-\delta)e^{-\lambda 1 t_i} + \delta e^{-\lambda 2 t_i}\right)$$

where $q_0$ is the production at $t=0$ and $\delta = \frac{a}{q_0}$ is the fraction of $q_0$ getting decayed by the second process.

We assume that the measured production $q_i$ has Gaussian noise of variance $\sigma_i^2$. We can therefore write

$$P(q_i|t_i, \sigma_i, q_0, a, \lambda_1, \lambda_2) = \frac{1}{\sigma_i\sqrt{2\pi}} e^{-\frac{1}{2}(\frac{q_i - q_i}{\sigma_i})^2}$$

In turns, this means that the likelihood of the data given the parameters can be expressed as

$$P(q|t, \sigma, q_0, a, \lambda_1, \lambda_2) = \prod_{i=0}^{N} \left(\frac{1}{\sigma_i\sqrt{2\pi}}\right) \times e^{-\frac{1}{2}\sum_{i=0}^{N}(\frac{q_i - q_i}{\sigma_i})^2}$$

where $q = \{q_1, q_2, ...q_N\}$, $t = \{t_1, t_2, ...t_N\}$, $\sigma = \{\sigma_1, \sigma_2, ...\sigma_N\}$, and $N$ is the number of observations.

This likelihood is maximized when the exponential term in minimized, which means that we need to find the set of parameters $\{a, q_0, \lambda_1, \lambda_2\}$ that minimize $\sum_{i=0}^{N}(\frac{q_i - q_i}{\sigma_i})^2$.

$q_i$ is actually the mean of $q_{ci}$ measured for different cells, that itself has some error bars $\sigma_{ci}$. We therefore have :

$$\sigma_i = \frac{\sqrt{\sum_{c=0}^{N_{ci}} \sigma_{ci}^2}}{N_{ci}}$$

where $N_{ci}$ is the number of cells we observe at time $i$.

**Assessment of sustained gene expression during prolonged starvation** Because no carbon source is provided in our experiments, bacteria only rely on their reserves to fuel their metabolism. Since an exponential decay corresponds to an infinite process, one would like to know how long the second exponential decay observed at low production levels persists before gene expression drops to zero. Because it is not possible to address this directly due to measurement errors, we decided to focus on the how the average volumic production evolves between 30 and 60 hours of starvation, in time windows of 10 hours (Fig. S6A). Taking the simple criteria that volumic production can physically not be negative, one concludes that the our production estimate is reliable down to ≈ 5 FP/$\mu$m/h (the observation at -4 for rpsB is considered to be an outlier). Notably, the standard errors are typically much smaller than that, which indicates that measurements across consecutive time points within 10 hours windows are very stable, but that some parameters change in an uncontrolled manner between promoters and/or experiments.

Of the 15 promoters above this accuracy threshold, only 1 shows no clear sign of decrease in volumic production from 30 to 60 h. Moreover, 10 out of 15 maintain a detectable production, while 4 decrease from an initially detectable level to levels below the threshold. Altogether, this sketches a picture where sustained gene expression during starvation is ubiquitous across promoters but will eventually drop to levels which are too low to be reliably measured, and hence probably to be functionally relevant.

#### 10 Regulators of synthetic promoters

The synthetic promoters used in this study have been selected to achieve a given expression level during exponential growth ( [24]). Importantly, prediction of transcription factors binding on their sequences indicates only binding sites for the sigma factor RpoD. Based on this, we selected them to assess how the global gene expression capacity of the cell change upon entry into starvation, in absence of specific regulation. Importantly, it is well established that the two sigma factors RpoD and RpoS recognize very similar motifs. Hence promoters transcribed under the control of one of them will also be transcribed due to the activity of the other, albeit with different intensity [50]. Given that RpoS is the main sigma factor active during early starvation, it is important to characterize the strength of the regulation by RpoS for our constitutive promoters.

Using ΔrpoS mutants, we were able to demonstrate that the concentration of the two medium expressors (Med2 and Med3) is dramatically decreased in this genetic background compared with the wild type levels, both during growth and starvation (Fig. S9A). Moreover, the increase of concentration observed for those promoters in the wild type upon switching from exponential growth to starvation is also abolished in the ΔrpoS background (Fig. S9B). Together, those observations strongly suggest that the main regulator of the medium expressors is RpoS, meaning that those synthetic promoters can be used as reporters for RpoS activity. On the contrary, the concentration and fold change of the high expressor Hi1 was only mildly affected in the ΔrpoS background compared to the wild type. This implies that Hi1 is primarily regulated by RpoD and we interpret it as a reporter of "unregulated" gene expression.

### 11 Quantitative proteomics measurements of protein levels

#### 11.1 Sample Preparation

In order to compare the temporal dynamics of the protein levels inferred from the GFP content of the cells and the actual levels of the native proteins produced by our promoter of interest, we used quantitative proteomics. In addition, we compared our results to the fold change between exponential growth and stationary phase conditions found in publicly-available proteomics datasets.

We conducted a low time-resolution experiment with several strains in parallel. Based on the concentration profiles measured in microfluidic experiments (Fig. 4A), we selected reporters covering the range of fold-change observed between exponential growth and starvation (10 out of 18 strains from the Zaslaver library): "pyrB", "codB", "rplN", "pliG", "gadX", "sbmC", "nrdD", "hslV", "hupA", "bolA". We collected 5 mL samples ($\approx 5.10^8$ cells in early exponential phase, at OD600 = 0.1) at 0 h, 1 h, 4 h, 24 h, and 48 h, starting from a 50 mL culture so that half of the initial volume remained at the end of the experiment.

For this, two days before the experiment, we inoculated a 2 mL culture in M9 + 0.2% glucose from individual colonies for each strain. After overnight growth, each culture was resuspended in 10 mL of fresh M9 + 0.2% glucose media at very low density (16000x and 32000x dilution), in order to have cultures still growing exponentially on the next day. For each strain, one culture still growing exponentially ($0.1 \leq$ OD600 $\leq 0.5$) was then diluted in a final volume of 50 mL of M9 + 0.02% glucose to an OD of 0.025 (time - 2h). When the culture reached OD600 = 0.1 (0h), we sampled 5 mL of the culture, washed 2 x with 3 mL of ice cold PBS, resuspended the pellet in 3 mL ice cold PBS, centrifuged at 4°C and flash-froze the pellet at -80ºC after carefully removing all supernatant. We repeated this sampling step at every time points.

Proteins were purified and digested with the SP3 approach [51] using a Freedom Evo 100 liquid handling platform (Tecan Group Ltd., Männedorf, Switzerland). In brief, Speed Beads ™(#45152105050250 and #65152105050250, GE Healthcare) were mixed 1:1, rinsed with water and diluted to the 8 µg/$\mu$L stock solution. Samples were adjusted to the final volume of 90 $\mu$L and 10 $\mu$L of the beads stock solution was added to them. Proteins were bound to the beads by addition of 100 $\mu$L of 100% acetonitrile to the samples, which were then incubated for 8 min at RT with a gentle agitation (200 rpm). After, samples were placed on a magnetic rack and incubated for 5 min. Supernatants were removed and discarded. The beads were washed twice with 160 $\mu$L of 70 % (v/v) ethanol and once with 160 of 100% acetonitrile. Samples were placed off the magnetic rack and 50 $\mu$L of digestion mix (10 ng/$\mu$L of trypsin in 50 mM triethylammonium bicarbonate) was added to them. Digestion was allowed to proceed for 12 hours at 37°C. After digestion samples were placed back on the magnetic and incubated for 5 min. Supernatants containing peptides were collected and dried under vacuum.

#### 11.2 LC-MS analysis

Dried peptides were resuspended in 0.1% aqueous formic acid and subjected to LC–MS/MS analysis using an Orbitrap Eclipse Tribrid Mass Spectrometer fitted with an Ultimate 3000 nano-LC (both Thermo Fisher Scientific) and a custom-made column heater set to 60°C using block randomization. Peptides were resolved using a RP-HPLC column (75 µm × 30 cm) packed in-house with C18 resin (ReproSil-Pur C18–AQ, 1.9 µm resin; Dr. Maisch GmbH) at a flow rate of 0.3 µL/min. The following gradient was used for peptide separation: from 2% B to 12% B over 5 min to 30% B over 40 min to 50% B over 15 min to 95% B over 2 min followed by 11 min at 95% B. Buffer A was 0.1% formic acid in water and buffer B was 80% acetonitrile, 0.1% formic acid in water. The mass spectrometer was operated in DIA mode with a cycle time of 3 seconds. MS1 scans were acquired in the Orbitrap in centroid mode at a resolution of 60,000 FWHM (at 200 m/z), a scan range from 350 to 1200 m/z, normalized AGC target set to 250% and maximum ion injection time mode set to 50 ms. MS2 scans were acquired in the Orbitrap in centroid mode at a resolution of 15,000 FWHM (at 200 m/z), precursor mass range of 400 to 900, quadrupole isolation window of 12 m/z with 1 m/z window overlap, a defined first mass of 120 m/z, normalized AGC target set to 1000% and a maximum injection time of 22 ms. Peptides were fragmented by HCD (Higher-energy collisional dissociation) with collision energy set to 33% and one microscan was acquired for each spectrum.

##### 11.3 Protein Identification and Quantification

To determine changes in protein expressions across samples, a spectral library free DIA analysis was carried out. Therefore, the generated raw files were searched against an *E. coli* database (downloaded from Uniprot on 20220222) and 392 commonly observed contaminants (total of 4764 protein sequences) using DIANN (v1.8.1) [52] and default settings with a few changes: MBR was activated, Protein names were selected for Protein inference and Neural network classifier were set to Double-pass mode. In the additional options box "–report -lib -info" was added. Quantitative results obtained from the report file were further analyzed using the MSstats R package v.4.7.3 [53, 54]. All mass spectrometry files associated with this manuscript are accessible at MassIVE (ftp://massive.ucsd.edu/v08/MSV000095680/).

##### 11.4 Proteomics fold change calculation from literature

For all datasets and all genes, the log fold change in sample i $F_i$ is calculated by taking the natural logarithm of the ratio of raw mass spectrometry signal in starvation condition(s) $M_{si}$ over the raw mass spectrometry signal in steady state exponential growth $M_g$ : $F_i = log(\frac{M_{si}}{M_g})$ . When replicates are available the average is used to do this calculation. For all datasets, the precise samples that are used and data sources are detailed below :

- Caglar et al. 2017 [55] : sample names, gene names dictionary and protein counts are obtained from github repository `https://github.com/umutcaglar/ecoli_multiple_growth_conditions/tree/master/`, downloading the files `Protein_reads_06_16_2016/sampleInformation.csv`, `generateDictionary/nameDictionary_RNA&Protein.csv`, and `Protein_reads_06_16_2016/Protein_Count_GlucoseTimeCourse.csv` respectively. Samples taken in exponential phase at 3, 4 and 5 hours are averaged and used as $M_g$. Fold change at 24, 48, 168 and 336h are calculated and represented on Fig. S7 as sample 1,2,3 and 4, respectively.

- Mori et al. 2021 [39] : description of the samples and protein counts are taken from Supplementary Tables EV2 and EV8. The sample named "LB mid log" is used as $M_g$. Fold change in late log phase in LB, as well as in stationary phase (following growth in LB) are calculated using the samples named "LB late log", "LB stat 1" and "LB stat 2", and represented on Fig. S7 as sample 1,2 and 3 respectively. Note that the two latter samples are repetitions of the experiments, and expected to give the same results.

- Schmidt et al. 2016 [56] : Table with combined global absolute abundance estimations from both datasets are taken from Supplementary Table S6. The counts in the "Glucose" sample are used as $M_g$. Fold change after 1 and 3 days are calculated from samples "Stationary phase 1 day" and "Stationary phase 3 days", and represented on Fig. S7 as sample 1 and 2 respectively.

- Proteomics data from this study : in order to calculate the fold change of native proteins, samples 1 and 2 (Exponential growth) for all 10 strains -that only differ by the plasmid they carry- are averaged and used as $M_g$. Fold change after 4h, 24h and 48h are calculated from the 10 replicates of samples 3, 4 and 5 (Starvation 1,2,3) and represented on Fig. S7 as sample 1, 2 and 3 respectively.

  In order to calculate the fold change of the GFPmut2 fluorescent proteins, we proceed as described above, with the exception that this time, all strains have to be treated as independent experiments -and not as replicates- since the promoter controlling the expression of the protein differs from strain to strain.

##### 11.5 Interpreting the changes of concentration

Reliably measuring the temporal dynamics of native protein levels during starvation or using related data from the literature proved to be challenging for individual proteins. In order to verify our ability to compare the fluorescence microscopy data and the quantitative proteomics data, we compared the dynamics of GFPmut2 levels expressed from 10 promoters that we previously used in our microfluidics experiments. Because those two methods measure variables that should be proportional to the same quantity (the concentration of GFPmut2 inside the cells), we expect their temporal dynamics to be similar. We observed that, even though the overall trend agrees relatively well between those two measurements, substantial differences were observed for some promoters such as bolA, pliG or sbmC (see Fig. S7, "Proteo exp. GFP" and "MM exp."). Comparing those dynamics to those of native protein levels revealed hard-to-interpret differences (see Fig. S7, "Proteo exp."). In only two promoters did the trend go in the same direction, namely for argC and pliG. For most promoters the signal in our proteomics data was very weak, suggesting instead that the concentration levels remain unchanged upon entry into starvation, which is at odd with our microfluidic experiments. Finally, two promoters showed a trend that is opposite to that of our previous results, with *e.g.* codB going down very sharply. Comparing those results to data available from the literature cautioned the interpretation of proteomics data for this type of comparison (Fig. S7). Although for some promoters there was a good agreement between datasets (e.g. bolA, pliG, sbmC, rpmB and rpsB), in most cases we observed diverging signal between or within the datasets, and in almost all cases highly fluctuating or missing values.

#### 12 A simple mathematical model capturing the diversity of concentration dynamics

The diversity of the observed protein concentration dynamics, can be captured by a simple mathematical model which is characterized by only 3 effective parameters. Moreover, we here show that the fold-change in protein concentration between exponential and late stationary phase can be calculated analytically as a function of these effective parameters.

First, we define $t = 0$ as the time of the switch to starvation and assume that the growth-rate drops instantaneously to zero, i.e.

$$\lambda(t) = \begin{cases} \lambda & ; \quad t < 0 \\ 0 & ; \quad t \geq 0 \end{cases} \tag{S20}$$

where $\lambda$ is the average growth rate before starvation. Note that in this simple model we track the average concentration dynamics as determined by the average growth rate, average protein production, and average protein degradation rate, i.e. we are not modeling single-cell variation.

As already discussed above, the growth-rate drops exponentially during stationary phase according to.

$$\gamma(t) = \begin{cases} \gamma_0 & ; \quad t < 0 \\ \gamma_0 e^{-\omega_\gamma t} & ; \quad t \geq 0 \end{cases} \tag{S21}$$

where $\gamma_0$ is the degradation rate during exponential phase and at entry into starvation, and $\omega_\gamma$ the decay rate of the degradation rate.

Finally, we need to define the protein production rate $q(t)$ as a function of time in stationary phase. Empirically, we have observed that, at the switch, protein production rate either immediately undergoes a sharp drop in production, followed by more slow decay, or shows an initial burst in production, followed by slow decay. Moreover, we have seen that the slow decay in production is often well described by a sum of two exponential decay functions.

We here simplify this observed diversity in production dynamics by assuming that, at the switch, production undergoes an instantaneous jump from $q_0$ during exponential growth, to $bq_0$ immediate after the switch. That is, the parameter $b$ denotes the fold change in production upon the switch to starvation. Although our model can be easily extended to have production decay as the sum of two exponentials, we will for simplicity describe the decay of $q(t)$ by a single exponential. That is, the protein production $q(t)$ is described by

$$q(t) = \begin{cases} q_0 & ; \quad t < 0 \\ bq_0 e^{-\omega_Q t} & ; \quad t \geq 0 \end{cases} \tag{S22}$$

where $w_Q$ is the rate at which production decays.

At the end of the exponential growth phase, the protein concentration $c_g$ will be given by

$$c_g = \frac{q_0}{\lambda}, \tag{S23}$$

where, for simplicity, we have neglected the degradation rate $\gamma_0$ since it is so much smaller than the dilution rate $\lambda$.

The dynamics of the protein concentration $c(t)$ during stationary phase is described by the differential equation

$$\frac{dc(t)}{dt} = bq_0 e^{-\omega_q t} - \gamma_0 e^{-\omega_\gamma t} c(t). \tag{S24}$$

We are particularly interested in the concentration $c_s$ late in stationary phase, i.e. as $t \to \infty$. Solving the differential equation (S24) we find that $c_s$ is given by the following integral

$$c_s = c_g e^{-\gamma_0/\omega_\gamma} + bq_0 \int_0^\infty \exp\left[-\omega_q t - \frac{\gamma_0}{\omega_\gamma} e^{-\omega_\gamma t}\right] dt. \tag{S25}$$

This integral can be solved using the substition $x = e^{-\omega_\gamma t}$, which turns the integral into the integral definition of an incomplete gamma-function. Moreover, we are especially interested in the fold-change $f = c_s/c_g$ in protein concentration.

We find for this fold-change

$$f = e^{-\gamma_0/\omega_\gamma} + \frac{b\lambda}{\gamma_0}\left(\frac{\omega_\gamma}{\gamma_0}\right)^{\omega_q/\omega_\gamma - 1}\left[\Gamma\left(\frac{\omega_q}{\omega_\gamma}\right) - \Gamma\left(\frac{\omega_q}{\omega_\gamma}, \frac{\gamma_0}{\omega_\gamma}\right)\right]. \tag{S26}$$

Importantly, the fold change $f$ depends on only 3 effective parameters that correspond to ratios of rates. That is, if we define $r_\gamma = \gamma_0/\omega_\gamma$, which controls how much degradation there is overall, $r_q = b\lambda/\gamma_0$ which sets the ratio in fold-changes of production and degration, and $r_t = \omega_q/\omega_\gamma$ which sets the ratio of time scales at which production and degradation decay. In terms of these variables we have

$$f = e^{-r_\gamma} + r_q (r_\gamma)^{1-r_t} [\Gamma(r_t) - \Gamma(r_t, r_\gamma)] \tag{S27}$$

**Lower limit on fold-change** To obtain the fold-change $f_0$ resulting from an instantaneous arrest of production, we just have to set $b = 0$ in equation (S26). We then find

$$f_0 = e^{-\gamma_0/\omega_\gamma}. \tag{S28}$$

Using the degradation rates measured in calibration experiments for the different fluorescent proteins (Table S5), we obtain lower limits in the range of 44 - 66% (Table S7).

| Fluo | $f_0$ |
|---|---|
| mCherry | 0.48% |
| GFP | 0.44% |
| mVenus | 0.66% |

**Table S7.** Predicted lower bounds on the fold-change $f_0$ in concentration for the different fluorescent reporters.

**Examples of the diversity in protein dynamics** To give some examples of the concentration dynamics resulting from different time-dependent protein production functions, we use a value of $\lambda$ corresponding to a doubling time of 1 h$^{-1}$, and set the initial concentration $c_g$ arbitrarily to 70 FP $\cdot\,\mu$m$^{-1}$. The initial degradation rates $\gamma_0$ and its decay rate $\omega_\gamma$ are set at 0.03 and 0.05 h$^{-1}$ respectively. $q_0$ is set at 50 FP $\cdot\,\mu$m$^{-1}\cdot$ h$^{-1}$. The other parameters used for the main text figure are displayed in TableS8

| Production dynamics | b | $\omega_q$ |
|---|---|---|
| Instantaneous arrest | 0 | $\infty$ |
| Gradual decrease (slow) | 1 | 0.1 |
| Gradual decrease (fast) | 1 | 0.2 |
| Burst and gradual decrease (fast) | 2 | 0.2 |
| Reduction and gradual decrease (fast) | 0.5 | 0.2 |

**Table S8.** Parameters of the concentration dynamics model

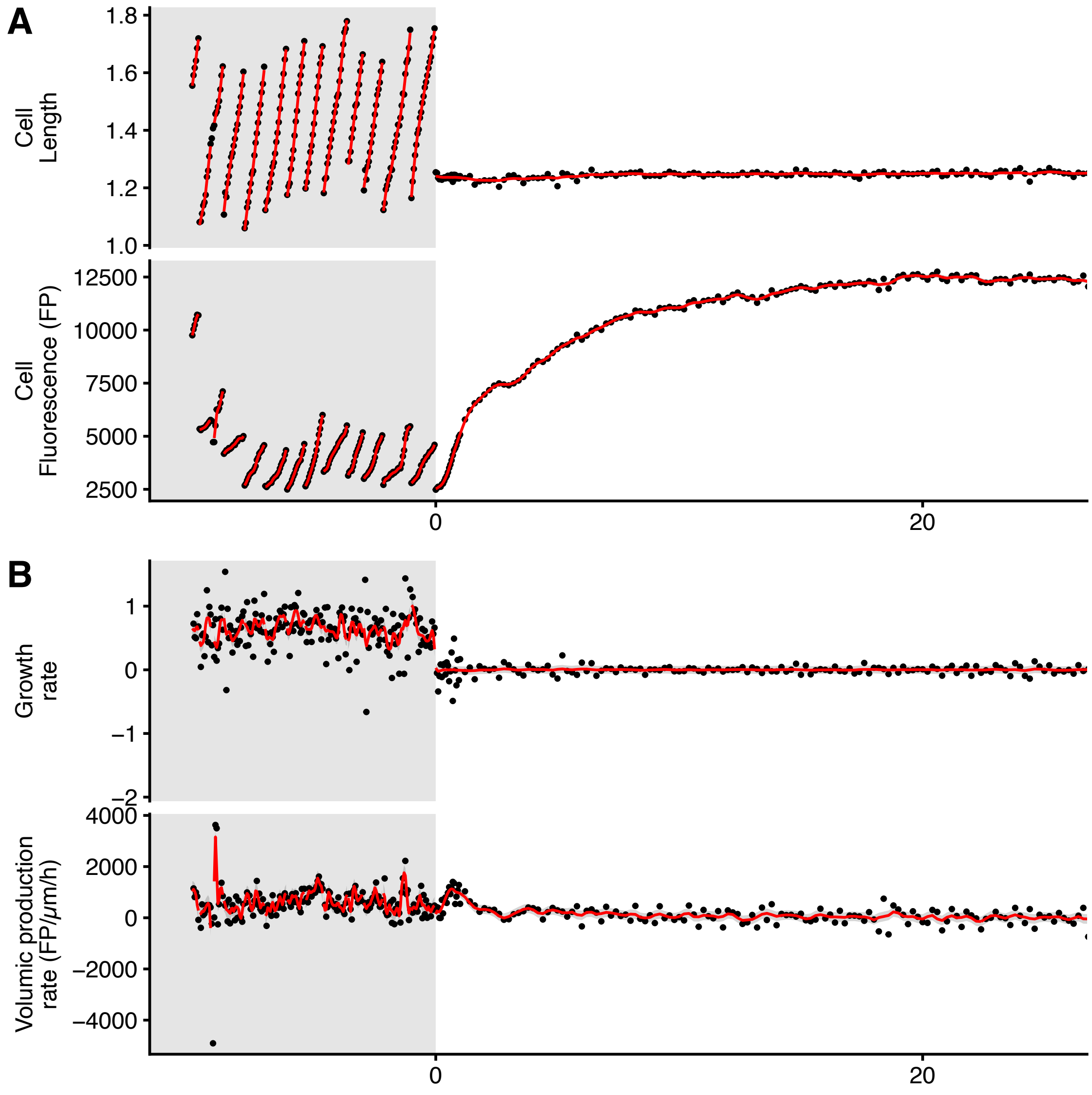


**Figure S1. Inference of instantaneous growth rate and production rate from noisy traces of cell length and fluorescence.** A model based on Gaussian prior process assuming that growth rate and production rate fluctuate as OU processes was used to predict their best estimate at each time point. (A) Raw length and fluorescence measurements (black dots) along with their inferred "true" values (red lines) from a randomly picked mother cell lineage. (B) Instantaneous derivatives calculation of the corresponding growth and production rates (black dots) and the corresponding inferred rates (red line). In A and B the grey ribbons surrounding the red line show the errors predicted by the inference.

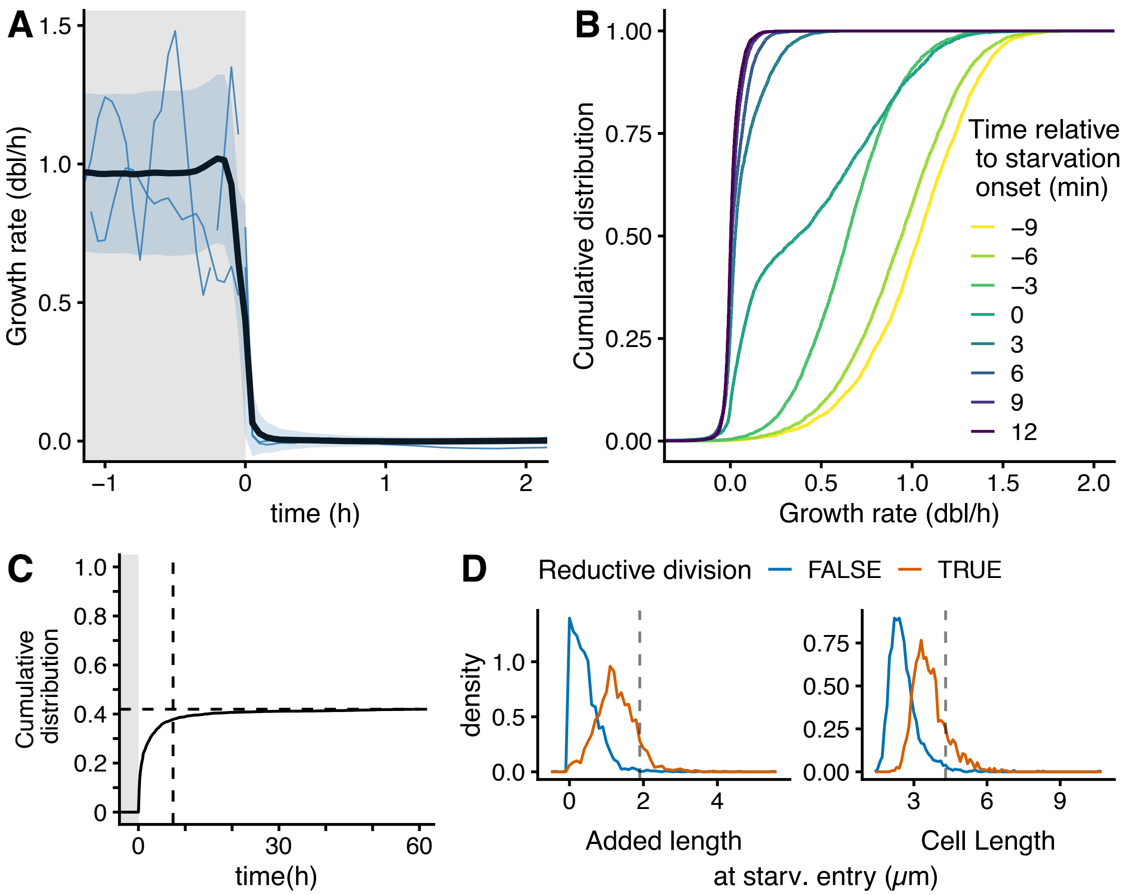


**Figure S2. Growth dynamics upon entry into starvation.** (A) After switching to "M9 zero" (defined as time 0), elongation stops almost instantaneously. The black line and ribbon show mean and mean ± s.d. of the instantaneous growth rate across all single cells, respectively; blue lines show randomly picked single cell traces. (B) Cumulative distribution of instantaneous growth rates across cells at each time point around the switch. All cells stop growing within 6 min. (C) Cumulative distribution of division across time, relative to the number of cells entering starvation. The horizontal dashed line correspond to the final fraction of cell that divided (42.6%), and the vertical dashed line indicate the moment where 90% of the divisions have happened. (D) Distribution of added length since last division and of cell length, measured at the point of starvation entry. Colors indicate whether cells undergo a division during the first 10 hours of starvation (called "reductive division").

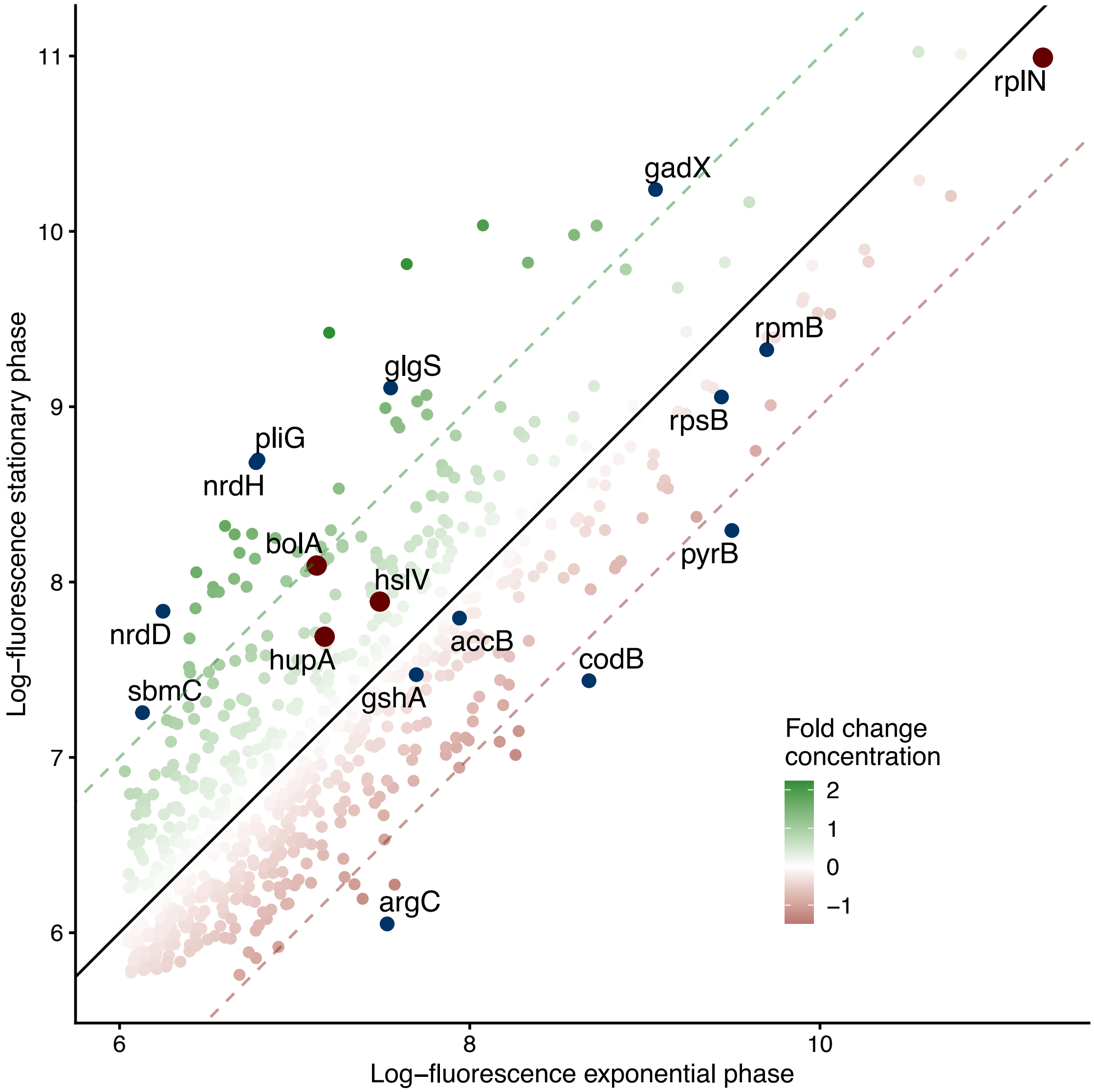


**Figure S3. Selection of native promoters.** Log of mean fluorescence measured in stationary phase cultures (after 2 days) as function of that measured in exponentially growing cutures (data from [23]). Every point corresponds to a promoter from the Zaslaver library [43], with the fold change of concentration between the two conditions indicated in color. Promoters are chosen so as to cover a broad range of fluorescence fold change and absolute levels (dark points; dark red points indicate promoters used as examples in the figures).

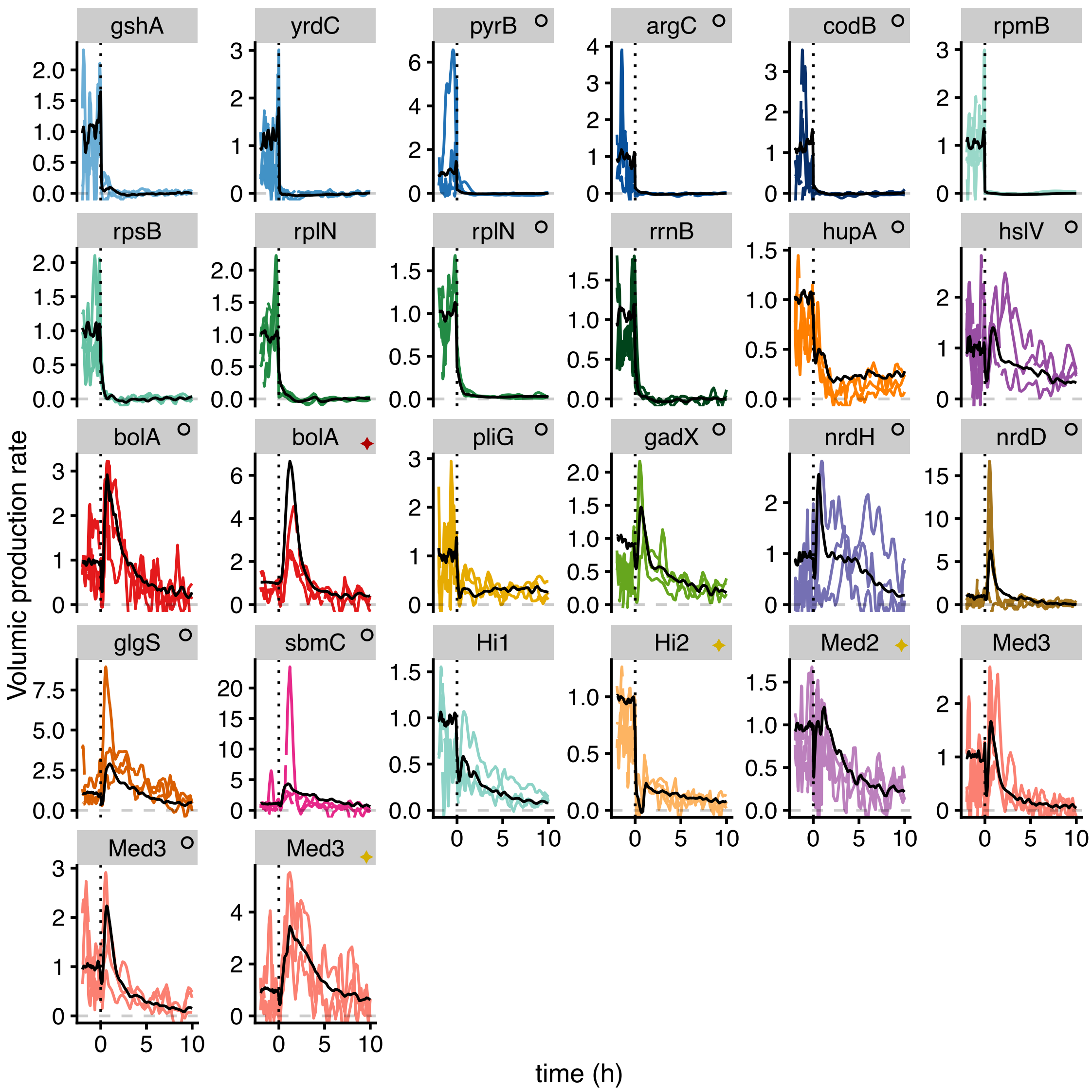


**Figure S4. Production dynamics of individual cells for all promoters at the onset of starvation.** Sampled single-cell traces of volumic production $q$ for the different promoters used in this study show a consistent response across the population upon entry into starvation for most promoters. The dark line represents the average volumic production rates of the various promoters. An empty circle next to the promoter name indicates a plasmidic reporter. The fluorescent protein is GFPmut2 by default; red or yellow diamonds indicate mCherry and mVenus, respectively.

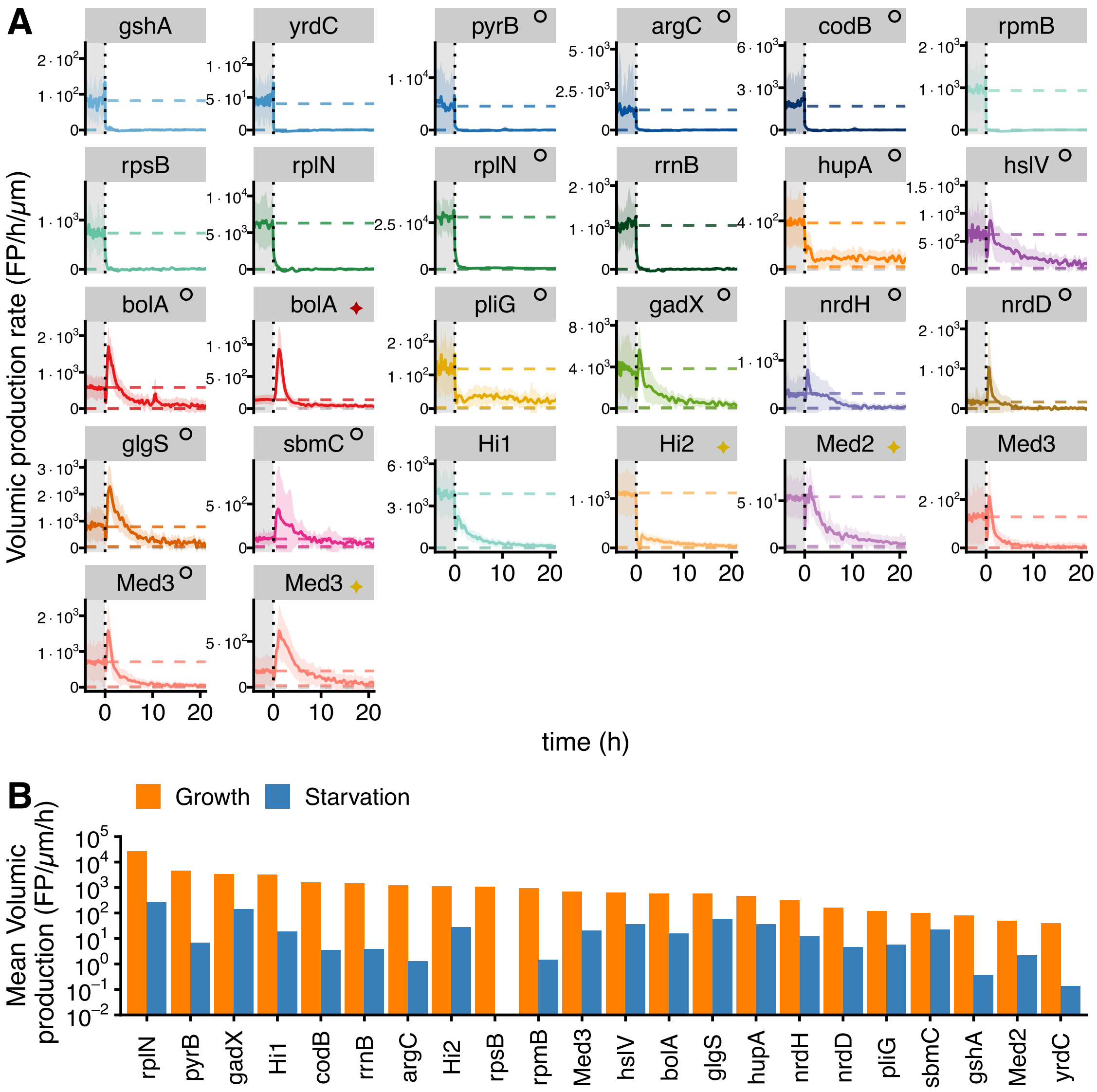


**Figure S5. Production dynamics of all promoters at the onset of starvation.** (A) Mean volumic production rate across time (only the first 20 hours of starvation are shown out of 60 hours). Plain lines correspond to average across all individual cells while ribbons correspond to mean ± sd. The average of volumic production during exponential growth is represented as the upper dashed line. Average volumic production for the last 30h of starvation is represented as the lower dashed line. An empty circle next to the promoter name indicates a plasmidic reporter. The fluorescent protein is GFPmut2 by default; red or yellow diamonds indicate mCherry and mVenus, respectively. (B) Average volumic production is computed during exponential growth, and during the last 30h of starvation. Note that the production does not always reach a steady state during the last 30h of starvation (see fig. S6).

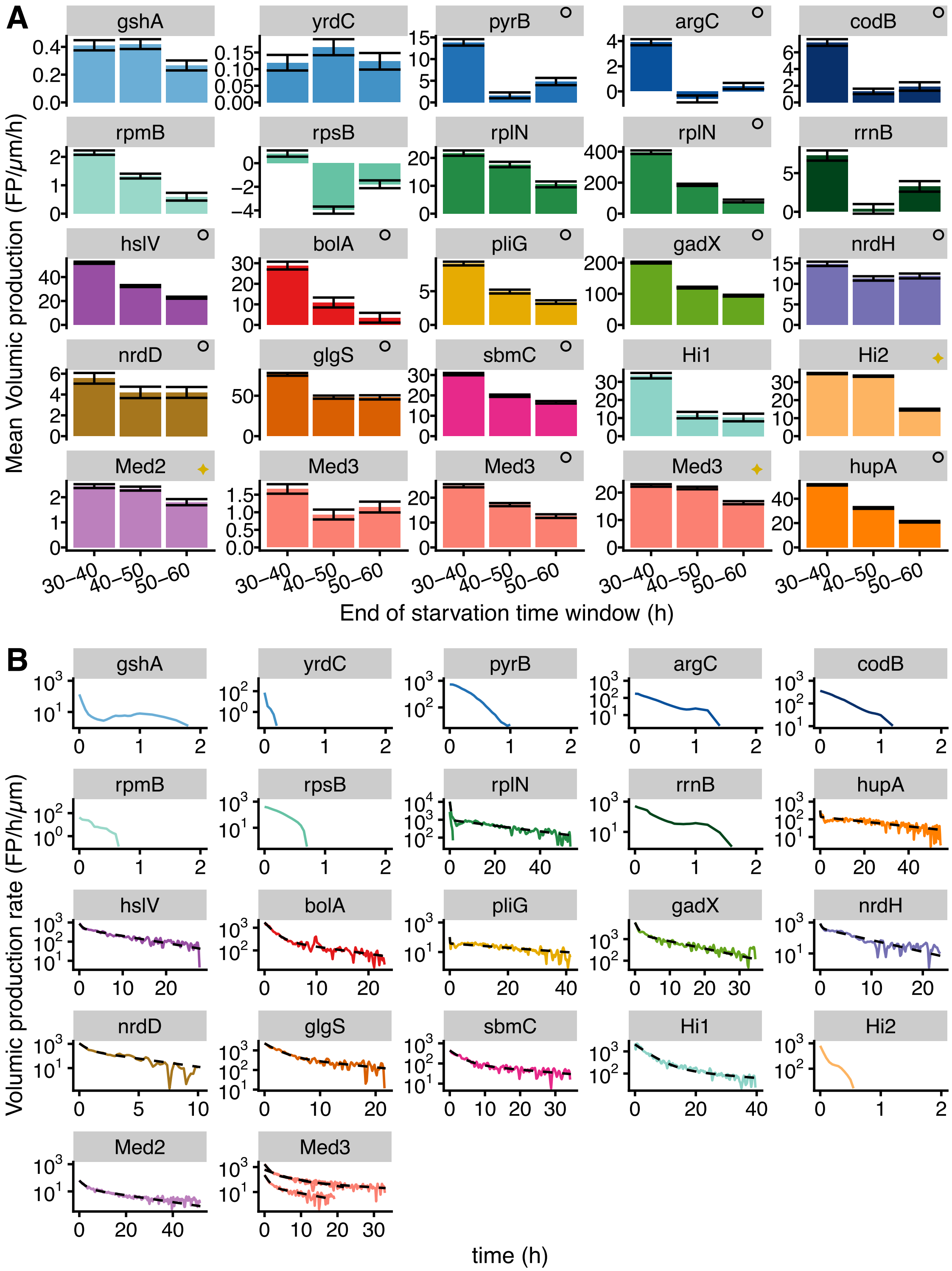


**Figure S6. The late dynamics of volumic production is promoter dependent.** (A) Average mean production over three 10 hours time windows covering the last 30 hours of starvation (±standard error of the mean). The type of reporter is indicated next to the names, as in S5. (B) volumic production rates on a logarithmic scale, between the maximal value (defined as time 0) and the first time point where it fluctuates below 0. Solid colored lines correspond to the average of volumic production across the population, and the black dashed lines correspond to fits to a sum of two exponentials. Only promoters with more than 30 observations were fitted.

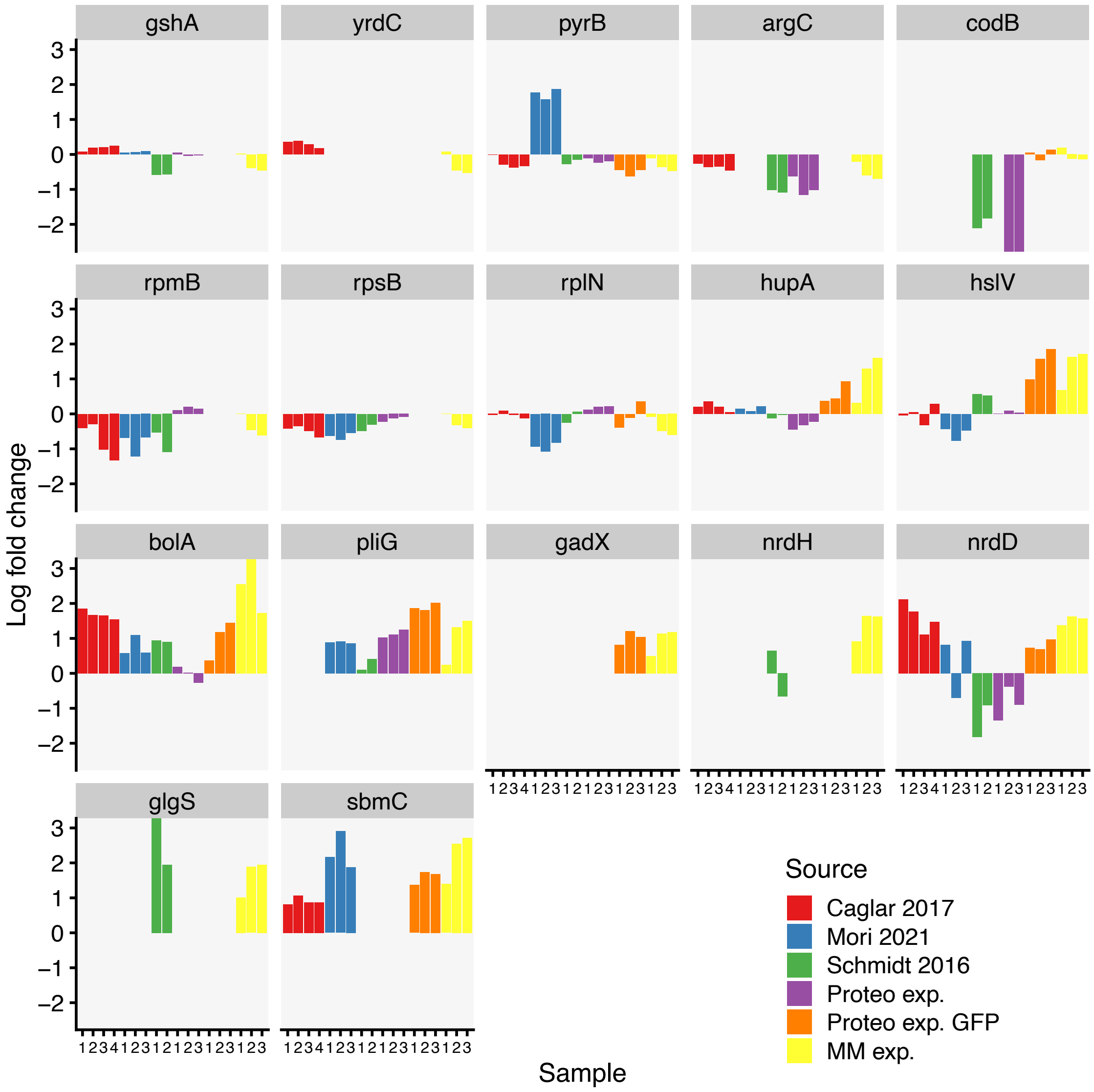


**Figure S7. Analysis of protein concentration changes between growth and starvation using quantitative proteomics.** In order to investigate the effects of protein-specific degradation on concentration levels, we performed quantitative proteomics experiments and compared the results with publicly available datasets [55, 56] [39]. Our experiments have been performed with strains from the Zaslaver library [43], allowing us to track levels of the native proteins (Proteo exp.), and levels of GFP whose expression is controlled by the promoters of those native genes (Proteo exp. GFP). In addition, we report the fold change observed in the mother machine experiments when using those reporters (MM exp.). We compare the log fold change of normalized mass spectrometry signal between exponential growth conditions and their corresponding starvation conditions. When multiple measures have been made in starvation conditions, we report them with increasing sample number (x axis). Note that some protein can not be quantified in some datasets due to technical limitations. For more details, see Section 11

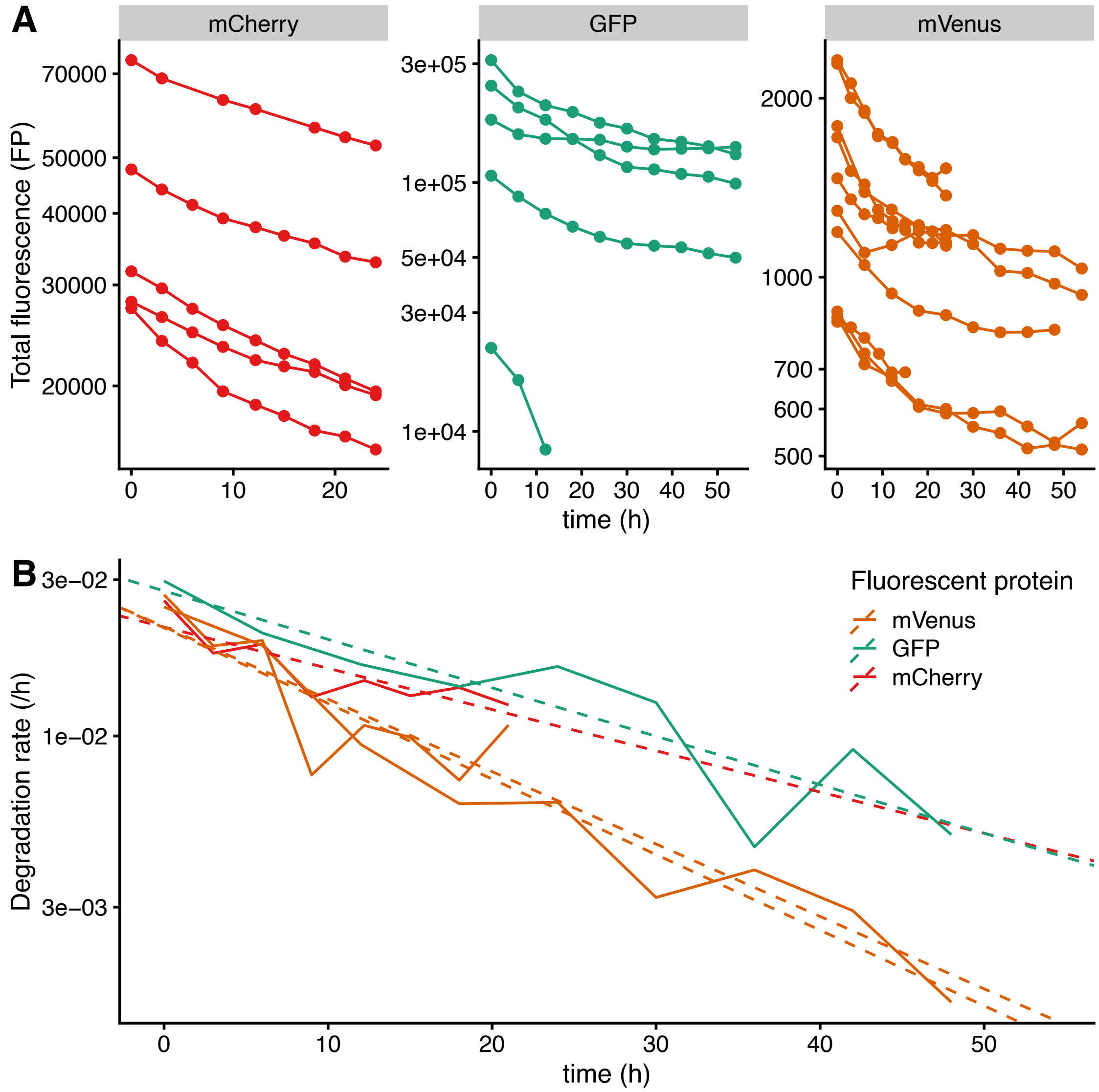


**Figure S8. Estimation of the degradation rates of fluorescent proteins.** Independent calibration experiments are performed with low acquisition frequency in order to ensure that photobleaching is negligible. (A) Random selection of traces of raw total fluorescence along time measured for starved bacteria carrying an inducible promoter, for which the inducer has been removed at t=0 (entry in starvation). The acquisition frequency was every 3 hours for mCherry2 and one mVenus replicate, and every 6 hours for the others. (B) Average instantaneous exponential degradation rate, defined as $\langle \frac{1}{F_c(t)} \frac{dF_c(t)}{dt} \rangle$ where $F_c(t)$ is the total fluorescence in cell $c$ at time $t$. The dashed line correspond to an exponential fit of this value as a function of time. $R^2$ of the fit are between 0.7 and 0.95. The initial values of degradation rates ( $0.03h^{-1}$) are comparable to the rates reported at vanishing growth [28].

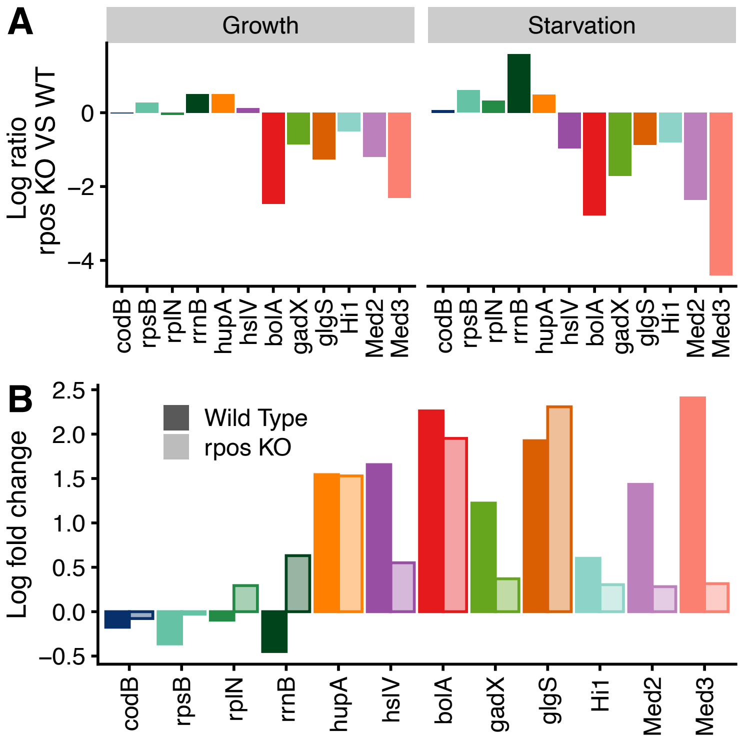


**Figure S9. Expression levels of all reporters in the ΔrpoS mutant.** (A) Natural log ratio of fluorescence concentration during the last 5 hours of growth or starvation (vertical panels) in the ΔrpoS mutant *versus* the wild type background. (B) Fluorescence concentration log fold-change from exponential growth to starvation in the ΔrpoS mutant (semi-transparent bars) and the wild type background (solid bars)

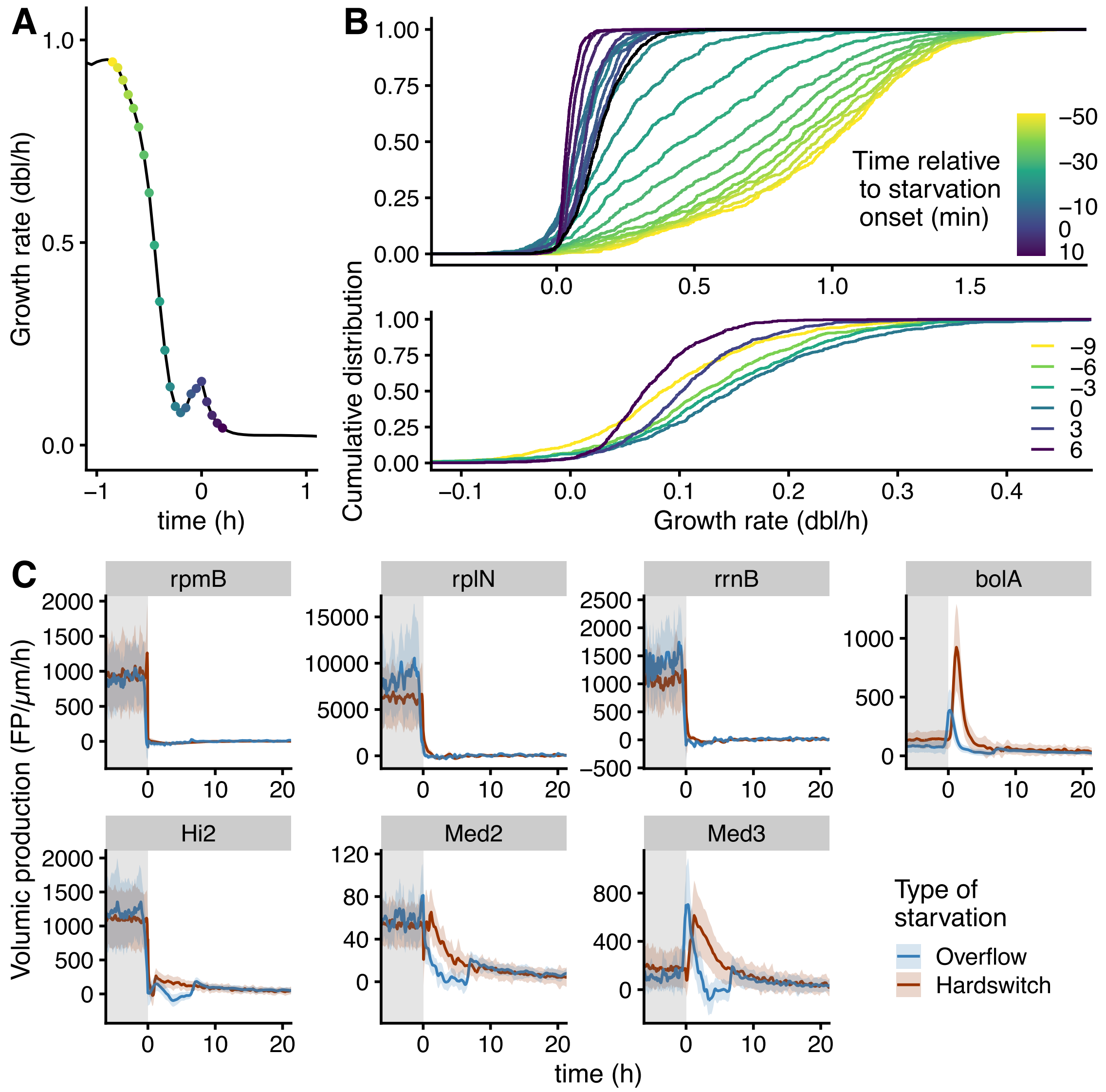


**Figure S10. Comparison of a hard switch with a progressive entry into starvation.** (A) The average instantaneous exponential growth rate of cells in the chip as a function of time in the overflow experiments. Individual time points during the sharp decrease in average growth rate, at 3 minute resolution, are indicated with different colored dots. The color gradient is aligned with time. (B) Cumulative distribution of instantaneous growth rates across cells at each time point from panel A. All cells stop growing within 60 min (top panel). Around time 0, the growth rate shows a transient increase and then decreases to 0 for all cells (bottom panel, zoom). (C) Mean volumic production rate versus time for every promoter that has been measured in the two microfluidic setups (indicated in colours). Only the first 20 hours are shown although the experiments lasted between 60 hours and 100 hours.

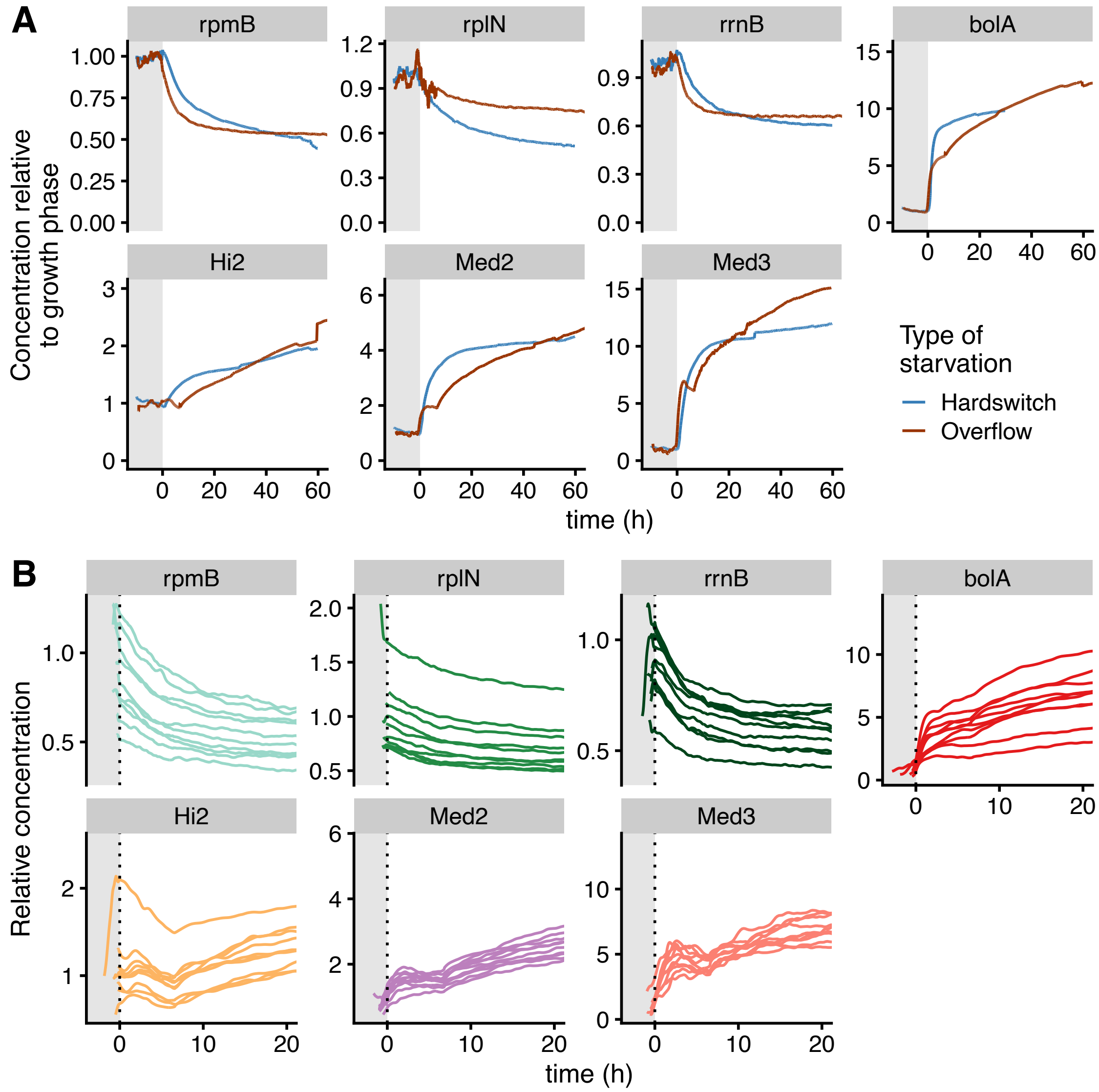


**Figure S11. Comparison of protein concentration dynamics for the hard switch and progressive entry into starvation.** (A) Average concentration relative to the average concentration during exponential growth as a function of time for every promoter that was measured for both the hard switch and culture overflow experiments (indicated in colors), calculated as in Fig. 2 (B) Relative protein concentration as a function of time for a random selection of single cells entering starvation. Each panel corresponds to a promoter used in the overflow experiments. The protein concentration of the single cells is scaled to the average concentration across the whole population during exponential growth. Only the first 20 hours out of >100 hours of starvation are shown.

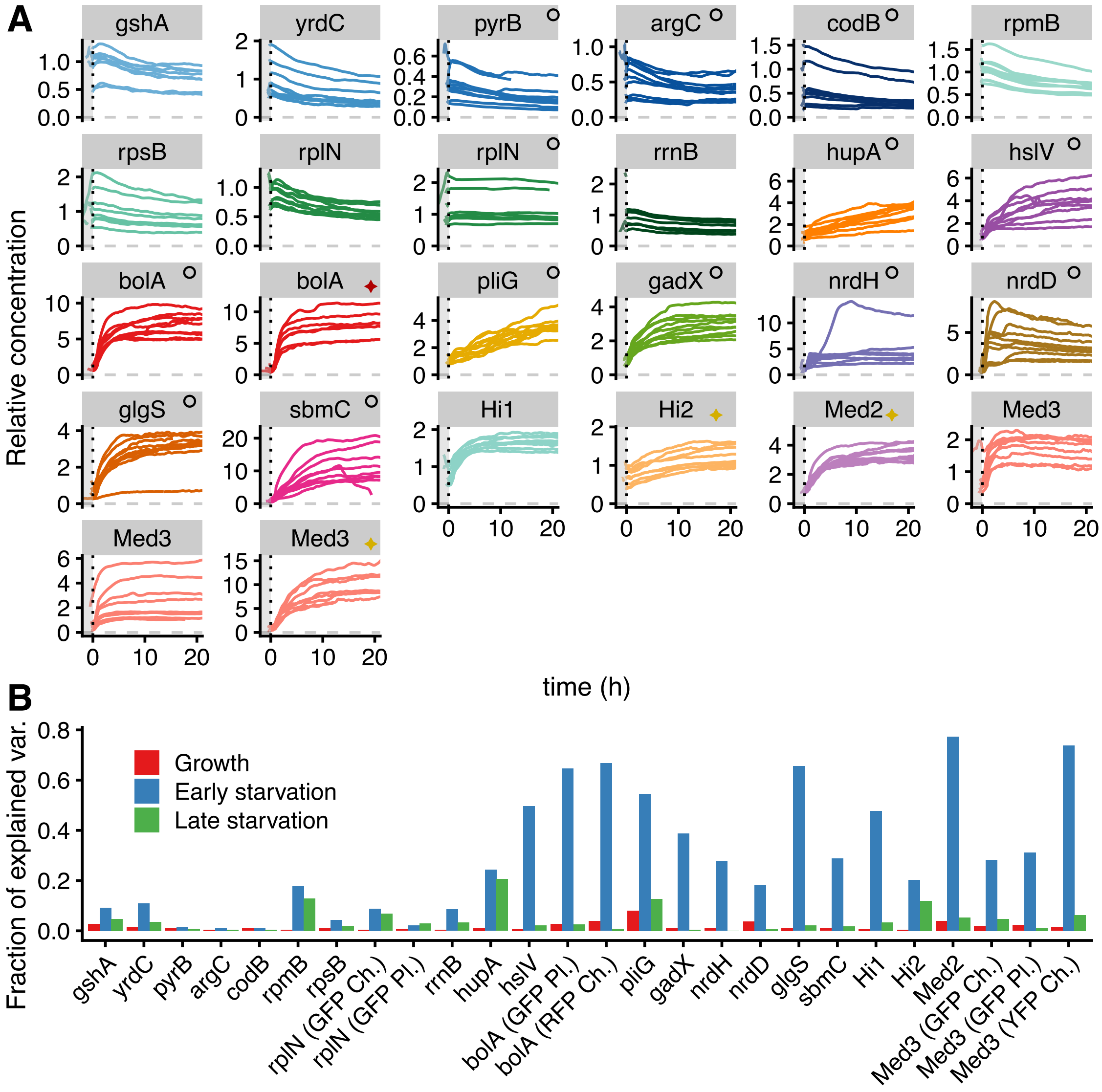


**Figure S12. Changes of concentration upon entry into starvation are consistent across individual cells.** (A) Protein concentration traces for a random selection of individual bacteria entering starvation. The concentration in single cells is normalized by the average concentration across the whole population during exponential growth. Only the first 20 out of 60 hours of starvation are shown. The type of reporter is indicated next to the name, as in Fig. S5. (B) The fraction $\rho$ of total variance captured by the variance in average concentration (computed as explained in the main text) is systematically higher during the first 10h of starvation (blue), compared to exponential growth (red) and late starvation (green). When a given promoter was measured with multiple reporters, the fraction of variance measured with each are shown separately. In this case, "Pl." indicates that the reporter was plasmidic, while "GFP", "YFP" and "RFP" indicate that the fluorescent protein used was GFPmut2, mVenus and mCherry2-L respectively.

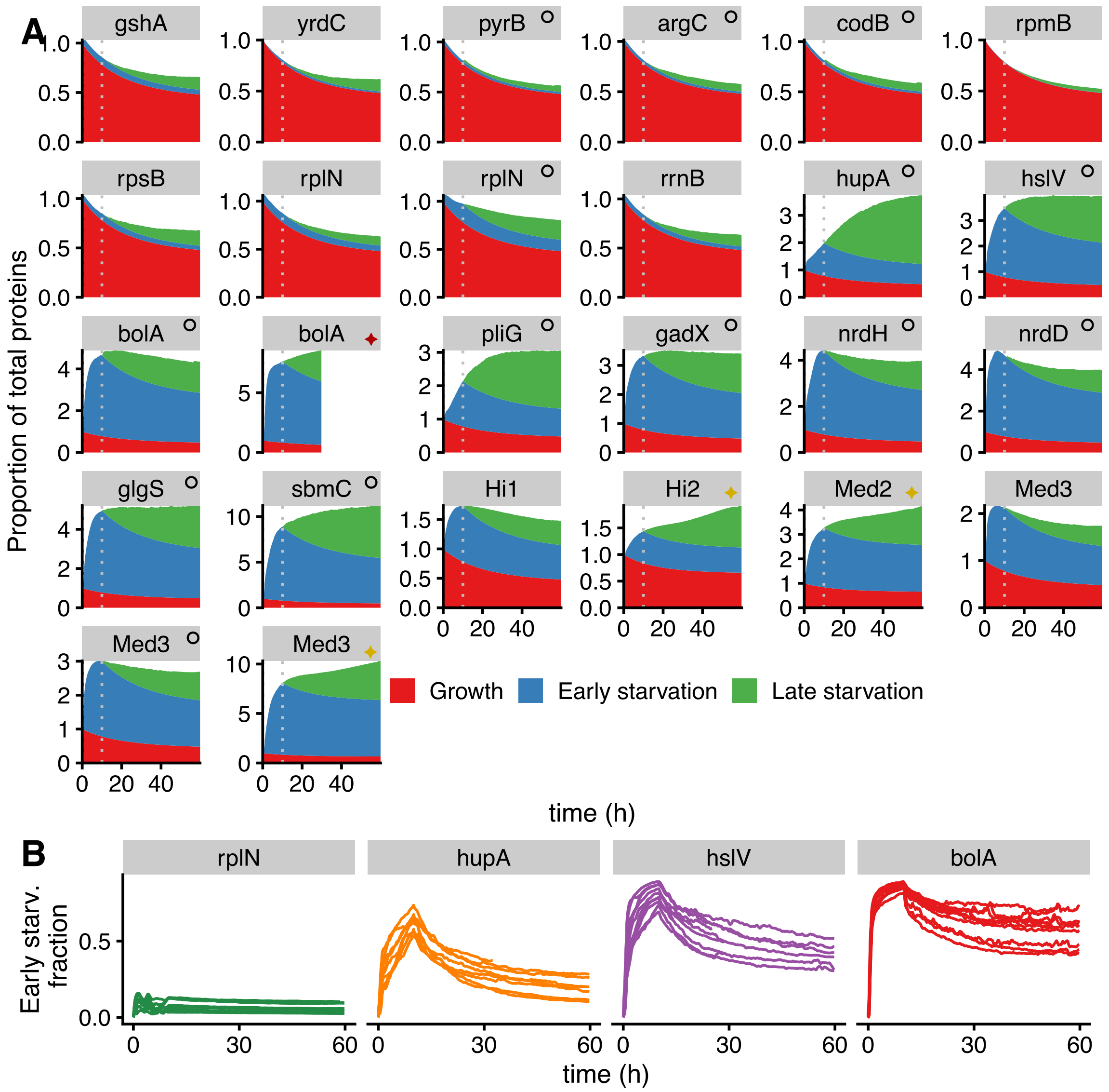


**Figure S13. Long-term impact of early gene expression dynamics.** (A) Proportion of total fluorescent proteins coming from defined periods of fluorescence production, as a function of time (as in Fig. 5A for all promoters). The type of reporter is indicated next to the promoter name, as in Fig. S5. (B) Fraction of the total fluorescent proteins coming from production during the first 10 hours of starvation, as a function of time, for a randomly selected subset of individual bacteria. Only the 4 promoters we focus on in the main text are shown here.

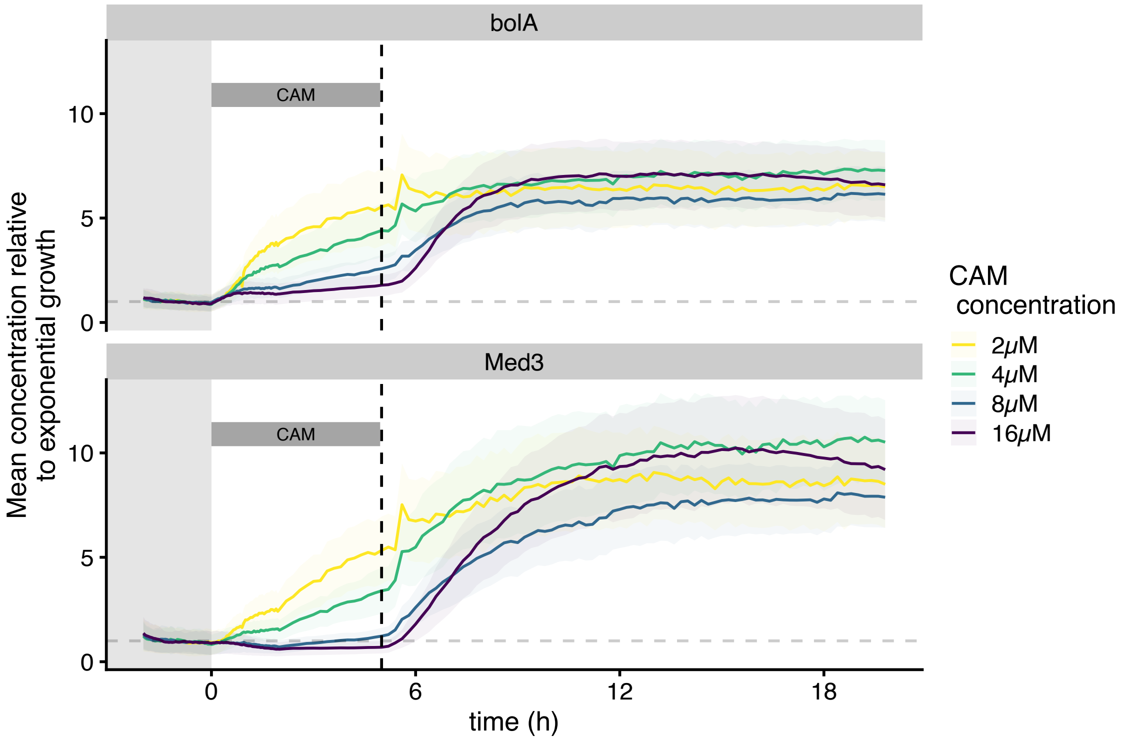


**Figure S14. Dynamics of gene expression upon entry into starvation with increasing inhibition of protein synthesis.** Chloramphenicol (CAM) was added at different concentrations (colors) at the start of starvation and removed after 5 hours (indicated by the vertical dashed line and CAM box). Solid lines correspond to the mean concentration across time normalized by the average concentration during exponential growth; ribbons correspond to mean ± sd.

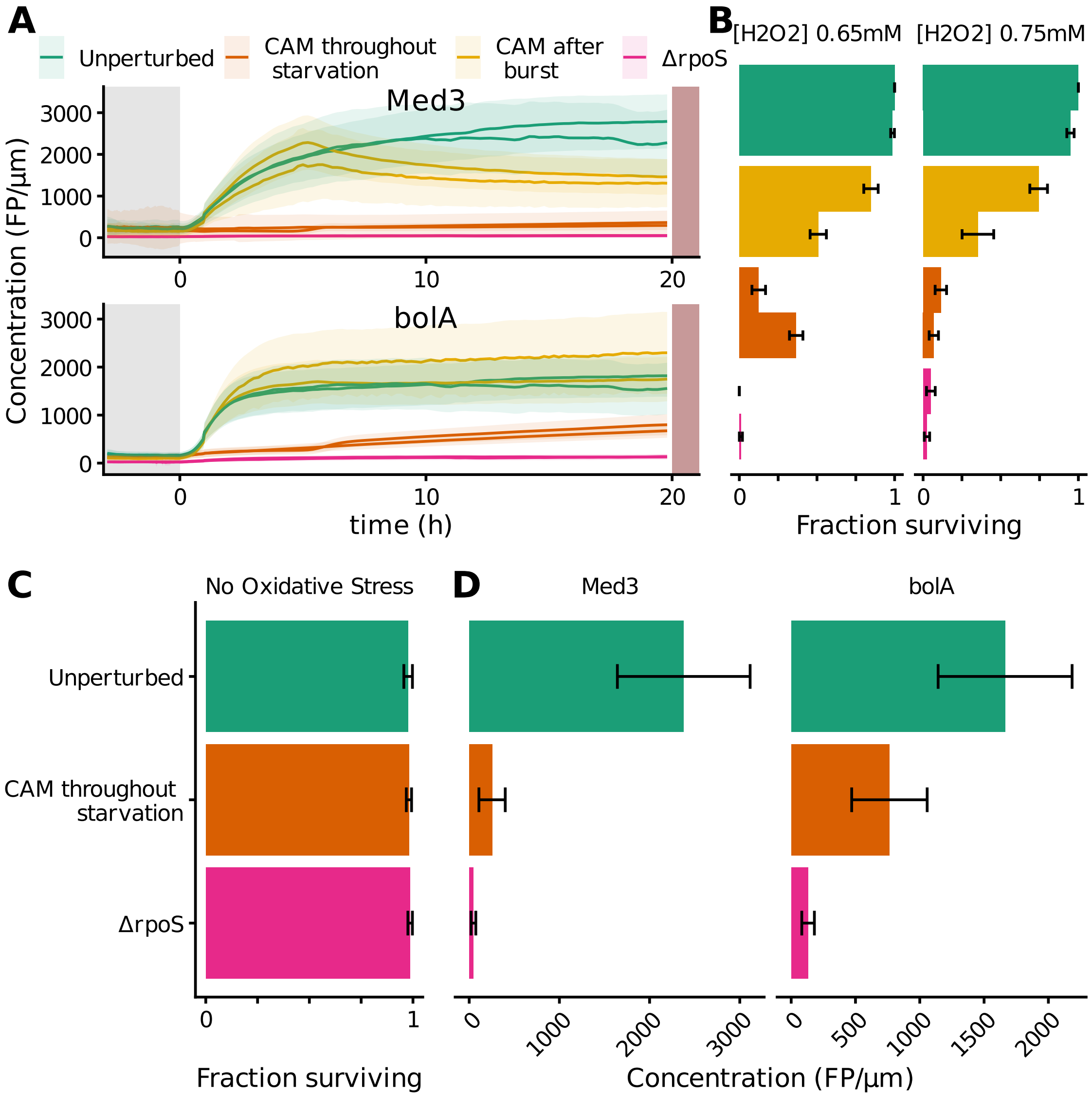


**Figure S15. Impact of gene expression perturbations on survival to oxidative stress during starvation.** (A) Average concentration levels for the two promoters used in the oxidative stress challenge experiments. Ribbons indicate mean ±s.d. The different colors correspond to different inhibition, as explained in Fig. 6 (2 replicates per promoter). (B) Fraction of surviving cells for all peroxide concentrations and replicates, as calculated in Fig. 6B. (C) Fraction of surviving cells for all control conditions, as calculated in Fig. 6B. (D) Average concentration after 20 hours of starvation in control experiments where no stress was used. C and D show that adding chloramphenicol or having rpoS knocked-out does not in itself affect the survival of starved cells on this timescale.

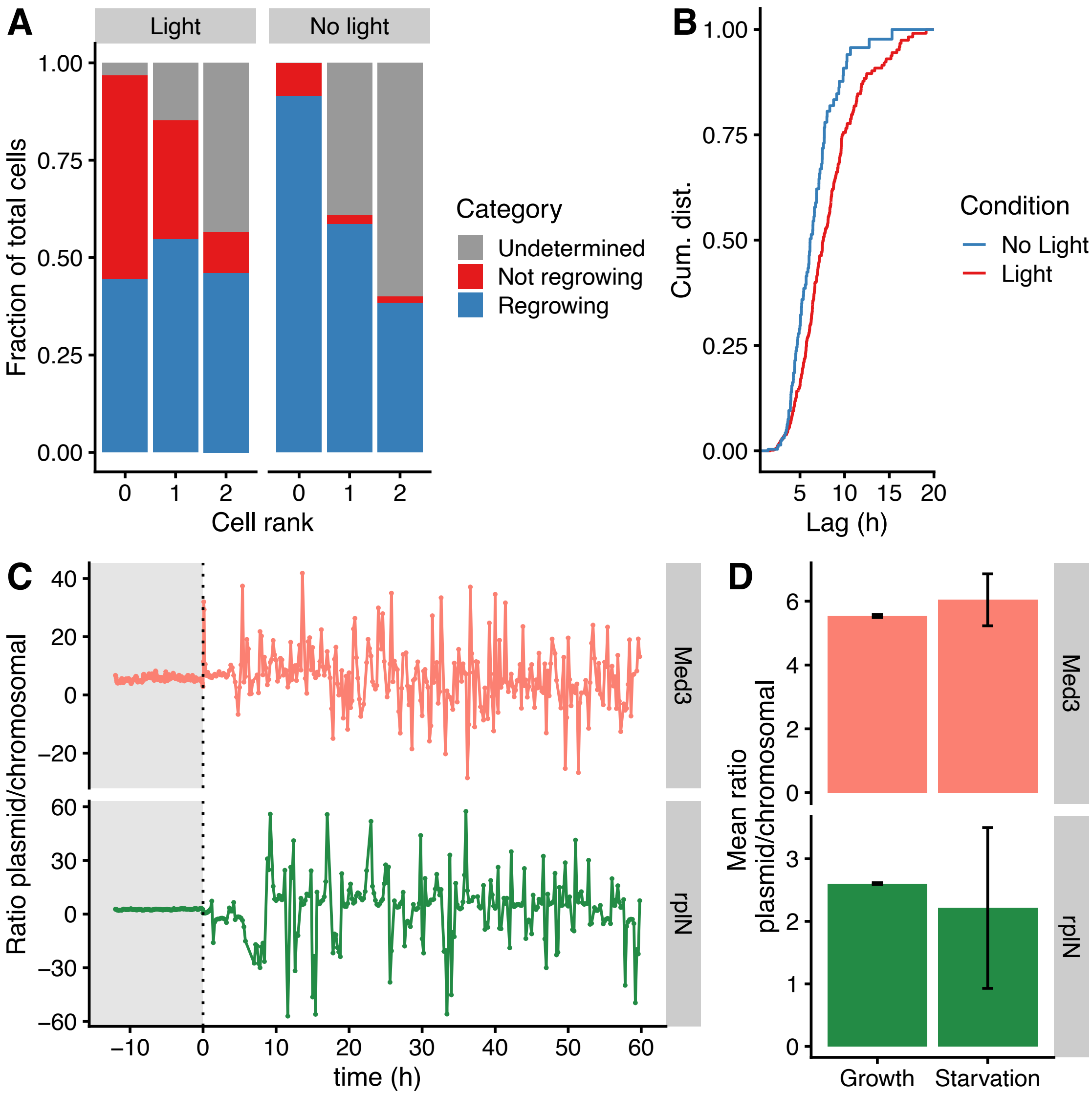


**Figure S16. Experimental setup controls for the hard switch experiments.** (A) The fraction of cells regrowing (observed dividing), not regrowing (observed lysing, or observed for more than 20 hours without regrowth), and undetermined (e.g. cells that are not observed long enough because they get pushed out of the growth channel) is plotted as a function of the cell rank in the growth channel, for the experiments performed with or without fluorescence illumination ("Light" vs "No light"). (B) Cumulative distribution of lag of regrowth for cells that divide after being re-exposed to fresh media. The lag is estimated as the delay between the switch and the point when the exponential fit on the last 5 time points of a cell cycle crosses the mean length at the exit of starvation. (C) Plasmid copy number is inferred by comparing the ratio of average Q of the plasmidic reporter and that of the chromosomal reporters for 2 promoters. During starvation, the promoter activity approaches 0 and becomes noisier, which explains the increase of the fluctuations in ratio amplitude. (D) Average of volumic production ratio between plasmidic and chromosomal reporters, during growth and starvation. Because the ratio can be very noisy during starvation, certain extreme points are driving the mean to artifactually high/low values. Those statistics have therefore been computed after removing the 2.5% highest and lowest ratio values. The difference of ratio between promoter is of unknown origin, and, even though it does not affect the changes of activity observed from growth to starvation, calls for caution when comparing absolute production levels between promoters.
